# A hybrid approach combining a phylogenetic method and Approximate Bayesian Computation Random Forest for phylogenetic network inference: application to the rice domestication process in Asia

**DOI:** 10.64898/2026.07.13.738130

**Authors:** Charles-Elie Rabier, Vincent Berry, Jean-Christophe Glaszmann

**Affiliations:** Institut Montpelliérain Alexander Grothendieck (IMAG), Université de Montpellier, CNRS, Montpellier, France; Univ Angers, Institut Agro, INRAE, IRHS, SFR QUASAV, F-49000 Angers, France; Laboratoire d’Informatique, de Robotique et de Microélectronique de Montpellier (LIRMM), Université de Montpellier, CNRS, Montpellier, France; CIRAD, UMR AGAP, F-34398 Montpellier, France; Amélioration Génétique et Adaptation des Plantes méditerranéennes et tropicales (AGAP), Université de Montpellier, CIRAD, INRAE, Institut Agro, Montpellier, France

## Abstract

Asian rice is one of the best documented crops in terms of genetic diversity. The domestication process, that probably started 9000 years ago in China, remains difficult to infer since the main vertical signal is blurred by horizontal signals related to gene flow among cultivars and wild relatives. Consequently, a large number of hypotheses on the domestication process of rice have been published. Besides, most of the methods used to infer these scenarios do not model all the known biological phenomena at stake.

Here, we present a methodological study based on a rich stochastic model, that incorporates introgression events, incomplete lineage sorting, and mutations that happen over time. The global evolutionary scenario is represented by a phylogenetic network. Furthermore, each locus scenario is modeled according to a locus tree through the *Multispecies Network Coalescent*. More importantly, for inferring the phylogenetic network, we propose a new hybrid approach combining a phylogenetic network method and a machine learning technique. In particular, our hybrid approach, named Snarf, benefits from advantages of a mathematical phylogenetic method, SnappNet, and from the potential of a powerful machine learning classifier, <u>i.e.</u> Approximate Bayesian Computation Random Forest (ABC-RF). These two methods are complementary since SnappNet reconstructs network accurately, whereas ABC-RF is able to handle a large amount of data. The originality is twofold. First, prior distributions required for ABC-RF are calibrated thanks to SnappNet’s estimates. Secondly, ABC-RF relies on summary statistics inspired by phylogenetic network literature.

We show, on simulated data, that the Snarf hybrid approach enjoys very good performances. On rice real data, it infers a scenario with a unique domestication (that of Japonica), followed by three reticulation events involving early Japonica. It highlights two introgression events at the origin of Indica and cAus, and one admixture event responsible for the emergence of cBas.

**Author summary:** Today, in genomics, there is a real need for methods able to infer phylogenetic networks. A phylogenetic network is a directed graph representing events like hybridization, introgression, and horizontal gene transfer. Understanding these complex biological phenomena, essential for crop adaptation, can help breeders when facing challenges like climate change and population growth.

Genome-wide diversity analysis thus requires network methods scaling for large data volumes and incorporating fundamental biological phenomena. In this context, we present a new hybrid approach, Snarf, that benefits from the potential of a powerful machine learning classifier, Approximate Bayesian Computation Random Forest, and from advantages of a mathematical phylogenetic method, SnappNet. Consequently, Snarf is able to handle large data-sets thanks to machine learning and is also based on a deep mathematical theory.

On simulated data, our hybrid approach performs very well. When applied to real rice genomic data, it supports a scenario with a single domestication event, that of Japonica. The analysis further highlights the role of early Japonica in the origin of both Indica and circumAus. Finally, it identifies an ancient admixture event, involving circumAus in the emergence of circumBasmati. Together, these findings confirm the importance of early rice history along the Himalayan region.

## 1 Introduction

### The rice domestication in Asia

Asian rice is one of the best documented crops in terms of genetic diversity thanks to a wealth of germplasm and sequence data. Current understanding of rice domestication features a long and complex process that probably started around 9000 years ago in China, in the Yangtze valley (e.g. [53, 43]). Next, it expanded its foundation by profuse gene exchanges with wild populations and other early domesticates along migrations routes throughout Asia. This foundation is yet far from being homogeneous since some cultivar groups are well-known while some may be still uncovered (cf. rayada group, [102, 69]). Moreover, detailed molecular genotyping studies reveal many introgressions among cultivar groups (e.g. [147, 30, 113, 47, 148]) and between wild and cultivated forms (e.g. [129, 51]).

Rice domestication being a complex process, various viewpoints have been expressed in the literature to explain the current diversity among varietal groups (see [26] for a review). Authors suggest either a unique domestication ([61, 140, 27, 53, 138, 51]) or multiple domestications [56, 29, 130, 146, 69] featuring up to three lineages corresponding to Japonica, Indica and circumAus (cAus). The various scenarios largely depend on the respective contributions of the three lineages in terms of domestication alleles. These alleles are responsible for new features that differentiate a crop from its wild ancestors.

The single-domestication scenario attributes the fundamental domestication syndrome to the Japonica lineage. It happened before its dissemination, accelerating the domestication of other forms. In contrast, the multiple domestication scenario considers that each of the lineages underwent human-directed transition from wild to crop. Next, crop feature exchanges lead to other lineages. However, some materials classified as wild can be the result of hybridizations involving some cultivars and producing predominantly weedy forms ([44, 80, 129, 139]). Generally speaking, the domestication process is hard to infer since the main vertical signal is blurred by horizontal signals related to gene flow, such as admixture or introgressions. Various materials may bear witness to various individual histories, challenging the extraction of predominant phylogenetic patterns.

Most of the methods (e.g. Neighbour-Joining tree [112] in [69, 51], Principal Component Analysis in [69], TreeMix [98] in [51]) used to infer the rice evolutionary scenarios, model only part of the biological phenomena that are known for being responsible for the differentiation of genomes. Moreover, these studies do not benefit from the last advances on reticulated evolution in phylogenetics. As a consequence, we present in this manuscript, an approach complementary to existing ones. We propose to tackle here the reticulate evolution of rice subpopulations thanks to a rich stochastic model and to a new hybrid approach combining a phylogenetic network method and machine learning.

Our model incorporates simultaneously introgression events, mutations happening over time, and Incomplete Lineage Sorting (ILS), an important source of discordance between gene trees (see [37]). Because we are able to deal with a wide range of materials, this enables us to actively contribute to untangling the species complexes often generated by the domestication process.

### Phylogenetic inference

In phylogenetics, species tree inference has been studied extensively for many years, giving rise to a large corpus of theory and methods [76, 33] that mostly rely on implicit or explicit mathematical models of evolution. More recently, machine learning techniques have also been proposed to infer species trees (e.g. Convolutional Neural Networks (CNN) [131, 123], residual neural networks [152], generative adversarial networks [122], random forests [9, 54] and reinforcement learning [10]).

There exist various contexts in which a tree does not faithfully describe the evolution of studied species. For instance, a species tree is unable to model complex biological events such as horizontal gene transfers (e.g. among prokaryotes [73], but also eukaryotes [124]), hybridization events (see [84], e.g. for yeast [91], wheat [46], citrus [137], fish [31]) introgressions (e.g. rice [30], sea bass [35], butterfly [38], sunflower [62]) or recombinations (mostly in viruses [128, 92]). By contrast, phylogenetic networks are able to capture all of these mechanisms due to their reticulate edges (e.g. [14]).

Phylogenetic network inference is currently a very active research field (e.g. [6, 8, 11]). Recent network inference methods rely on mathematical models of evolution and they produce explicit networks, that is networks in which reticulation events explicitly correspond to an evolutionary event like a lateral transfer or an hybridization. Most of these methods model ILS and hybridization simultaneously.

Note that in this network context, ILS is usually modeled according to the *Network Multiple Species Coalescent* (NMSC) [75, 89, 141], a variant of the well-known *Multiple Species Coalescent* (MSC) model [103, 72] heavily used in species tree inference.

Methods inferring explicit phylogenetic network can be divided in several categories: Maximum Likelihood (ML), Pseudo Likelihood (PL), Combinatorial Approaches (CA) and Bayesian Methods (BM). The studies of [142] and [143] are examples of ML approaches handling ILS. In [142], Yu et al. derived the probability of a gene tree topology within a phylogenetic network, using multilabeled trees. Later, they added the possibility to account for branch lengths [143]. In contrast, the ML method called NetRAX [82] presents the drawback of not dealing with ILS. When facing large genomic data sets, ML methods are intractable, so studies usually turn to approximate solutions, as provided by PL methods. The latter rely on a two-step approach. In a first step, the phylogenetic network of interest is divided in sub-networks with a small number of taxa, and the likelihood of each sub-network is computed. In a second step, these likelihoods are combined in order to obtain a “pseudo” likelihood of the entire network. For instance, the PL methods named SNaQ [121] and PhyNEST [74] rely on quarnets (4-leaf level-one network) whereas that of [150] focused on trinets (3-leaf level-one network). CA methods are a quite different kind of methods. They consist in enumerating the number of possible outcomes without computing a likelihood. For instance, TINNIK [6], implemented within the MSC-quartets [109] R Package, is a fast method that can handle many taxa without being restricted to level-one networks, unlike SNaQ.

Finally, many BM methods have been proposed over the last decade. They have the advantage of providing a distribution over networks, from which the uncertainty of the inferred clades can be explored. Wen et al. [132] introduced the first Bayesian method. Note that since this method takes gene trees as input, gene trees needs to be estimated in a preliminary step, as for SNaQ and TINNIK. Next, the SpeciesNetwork [144] and MCMCBiMarkers [149] methods were proposed, respectively implemented in Beast 2 [18] and PhyloNet [133]. Over the previous method, they offer the advantage to take sequence alignments as input, hence avoiding estimation errors in the intermediary step inferring gene trees [125]. More recently, we introduced the SnappNet [101] BM method, implemented within the Beast 2 framework [18]. SnappNet is based on the same mathematical model as MCMCBiMarkers, except that it considers a different network prior distribution. Both methods can infer a network of any level. The advantage of SnappNet over MCMCBiMarkers, lies in the fact that the likelihood computation is extremely faster, thanks to the mathematical treatment. In particular, whereas MCMCBiMarkers is based on an exhaustive combinatorial approach, SnappNet computes joint probability distributions. The maximum number of random variables involved in these joint probabilities is governed by a new concept, called the <u>scanwidth</u> of the network [13]. This approach has opened new perspectives on directed acyclic graphs (see [60]) and has a link with belief propagation in graphical models (see [127]).

Despite the progress made in the studies cited above, computing a likelihood in phylogenetic networks is still a time-consuming operation given the considered evolutionary models.

With regard to artificial intelligence, very few works proposed to apply machine learning methods to reticulate evolution. Some concentrate on introgression detection either through CNN ([117, 16]) and multilayer perceptron ([22]) or through classifiers trained on Summary Statistics (SS) (see [106, 58]).

To date, in phylogenetics, the superiority of machine learning methods over traditional methods has not yet been established yet [114, 90, 151], and there is no guarantee that this will be the case in the years to come. From a general perspective, although deep learning methods are becoming increasingly popular, they suffer from a lack of robustness [151] and the black box problem: since it is difficult to understand how decisions are made by the algorithm, this leads to a lack of interpretability and it is challenging to determine how to reuse the same tools in another context or on new data.

A current challenge is to combine both approaches by devising phylogenetic network inference methods that are based on a solid theoretical background, while also benefiting from the potential of machine learning.

Few pioneering works explored this track. For instance, Bernardini et al. [12] proposed a heuristic method combining the random forest machine learning and the CA classical approach. More precisely, their method aims at finding a network that represents a set of input trees while minimizing the number of reticulation events. In the current paper, we also propose a hybrid approach to infer phylogenetic networks, but based on a BM method. More precisely, our method combines a mathematical phylogenetic method, SnappNet, and a powerful machine learning classifier, Approximate Bayesian Computation Random Forest (ABC-RF). These two methods are complementary, as SnappNet offers good performances in terms of network reconstruction and parameter estimation, while ABC-RF is capable of processing large amounts of data.

### The concept of ABC-RF

Let us first recall the Approximate Bayesian Computation (ABC) approach. ABC is a family of simulation-based methods for approximating posterior distributions, which in turn can enable to estimate parameters or to select a model among several ones, see [85] and [119] for reviews. ABC consists in simulating numerous data sets and comparing them to the real data. The comparison is based on summary statistics (SS), which act as a proxy for the information contained in the data. Simulated data whose statistics are evaluated to be close to that of the original data, can then be used to approximate the posterior distribution. ABC methods [111] present the advantage of being likelihood free. In other words, the likelihood computation is not required, which makes ABC particularly useful when the likelihood cannot be expressed analytically or is computationally intractable due to a complex statistical model. ABC methods were proposed to address problems in biology [126, 99] and have since become very popular in population genetics (e.g., [66, 17, 87, 68]). They have also been used in phylogenetics (e.g., [67, 115, 57]).

The ABC-RF variant, introduced by [100, 40, 107], differs from ABC since it is not necessary, with ABC-RF, to estimate the posterior probability distribution to select the most probable model.

More precisely, ABC-RF consists in building a classifier on the basis of the very popular concept of random forests (RF) [19, 118] and of a reference table containing a large number of SS. In a nutshell, RF is a learning method that consists in aggregating results of multiple randomized decision trees. It helps to create a more robust classifier, <u>i.e.</u> reducing overfitting. Once the ABC-RF classifier is built, it can be applied to the real data. ABC-RF is more flexible than ABC since the choice of SS is not as crucial as in latter. In classical ABC, the choice of bad SS leads to a large number of rejections, and consequently, a huge number of simulations is required to approximate the posterior distribution [100]. In contrast, ABC-RF does not rely on an accept-reject method, and we can easily fill the reference table thanks to simulations from prior distributions. Note that with ABC-RF, it is also possible to approximate the posterior probability of the selected model. The ABC-RF approach has been fruitful in several contexts (e.g. [25, 45]).

### The SNARF hybrid approach

As mentioned before, our new hybrid approach relies both on SnappNet and on ABC-RF. The originality is twofold. First, prior distributions required for ABC-RF are calibrated thanks to SnappNet’s estimates. These estimates are obtained by running SnappNet in a penalized likelihood framework, a novelty compared to [101]. Secondly, ABC-RF relies on SS inspired by the phylogenetic network literature. As far as we are aware, this is the first time that the ABCRF approach is used for phylogenetic networks in a probabilistic framework. Up to now in the phylogenetics literature, we denote only two studies, [16] and [12], requiring to ABC-RF. Bernardini et al. [12] present an algorithmic approach whereas Blischak et al. [16] compare the performances of their devised CNN with those of a RF classifier based only on several phylogenetic statistics.

In our current ABC-RF study, the SS are specific to phylogenetic networks and computed under the NMSC. These SS have been implemented within the SimSnappNet simulator [101], companion of the SnappNet inference method. A large proportion of the SS we selected relies on phylogenetic invariants ([24, 78]). Recall that these invariants are polynomials in the frequencies of site patterns in idealized data (see [7]): they present the particularity to be equalling zero for a specific tree or network topology. Invariants are an extensive area of research in phylogenetics [41, 23], with recent works being focused on networks [49, 50, 136, 11]. The method we propose here relies on the invariants proposed by Kubatko et al. [77] and Pease et al. [97]. These works both consider the presence of ILS and introgressions (or hybridization) in their models. They provide statistical tests in order to test whether the discordance between a gene tree and a species tree is simply due to ILS, or whether it can be attributed to introgression events. More precisely, Kubatko et al. concentrate on the model of [89, 75] based on parental trees displayed by the phylogenetic network (<u>i.e.</u>, trees obtained from the network), whereas Pease et al. investigate a model that allows different kinds of introgressions (e.g. <u>inter-group</u>, <u>intra-group</u>, <u>ancestral</u>).

### Roadmap on rice evolution

In the present study, we apply the new method described above to the case of rice evolution. As input, we consider data sets formed from the multiple sequence alignments (MSA) from [61] and [130]. We consider from one to ten varieties per rice subpopulation and up to a maximum of 64 varieties in a same data set.

Our analysis is organized in three steps. First, we freely explore the whole space of evolutionary scenarios, thanks to SnappNet in a Bayesian framework. In order to cover a large genetic diversity, we consider a large number of small rice data sets, but with only one individual per subpopulation. This leads to multiple evolutionary histories inferred by SnappNet on our different data sets. Most of the inferred topologies are networks rather than trees: this highlights the importance of genetic exchanges in rice and the importance of later considering more than one variety per subpopulation.

Next, the goal of the second and third steps is to obtain a consensus scenario by adding more genetic material. The second step can be viewed as an initial step for calibrating our new hybrid approach, named Snarf, that will be applied in third step. More precisely, the second step consists in focusing on 16 rice evolutionary scenarios of interest. These networks were chosen on the basis of features found in the Bayesian analysis of our first inference step. We studied these 16 networks on the basis of larger data sets (<u>i.e.</u> more MSAs). With the help of SnappNet in a penalized likelihood framework, we evaluate the likelihoods of the 16 phylogenetic networks and we select the six best networks among them, according to the AIC criterion.

Lastly, in a third step, we concentrate on ABC-RF using priors calibrated thanks to the six evolutionary scenarios obtained at step two. For this third step we increase the sample size in order to exploit the potential of ABC-RF to handle large datasets. To begin with, we train our ABC-RF classifier on simulated data. Next, we use this classifier on real rice data to discriminate between the six scenarios. We show that our hybrid approach achieves very good performances on simulated data, regardless of the prior under study. On real data, we come to the conclusion that Asian rice underwent a unique domestication, that of Japonica. These results are in accordance with numerous papers on rice evolution ([61, 140, 27, 53, 138]), suggesting a unique domestication. But our method provides much more details: our number one ranked scenario shows an initial differentiation between Japonica and its wild ancestor, followed by three reticulation events involving early Japonica. The first two reticulations refer to introgressions from Japonica for the emergence of Indica and cAus, respectively. Note that these introgressions involve different groups of wild ancestors. The estimated inheritance probabilities reflect very small contributions of Japonica to Indica and to cAus. These contributions must be restricted to genome segments bearing alleles of the domestication syndrome. Last, the third reticulation event at the origin of cBasmati, depicts admixture between early cAus and early Japonica. This scenario reveals a more balanced composition between pre-existing cultivated lineages.

Overall, this study highlights the capacity of the Snarf hybrid approach we propose to account for various modes of crop evolution, and yet stresses the importance of the sample size for retrieving a pertinent global picture.

## 2 Material and Methods

### 2.1 SnappNet’s stochastic model

Let us briefly recall the stochastic model behind SnappNet.

#### 2.1.1 Phylogenetic network

A phylogenetic network is a directed acyclic graph (e.g., the 7 species network in Figure 1). Network branches are directed from top to bottom: they go forward in time. A phylogenetic network contains two kinds of internal nodes. A node is either (a) a <u>speciation</u> node, that is a node with one incoming branch and two outgoing branches, or (b) a <u>reticulation</u> node, that is a node with two incoming branches and one outgoing branch. The oldest ancestor is the node at the top of the network, while the <u>leaves</u> are nodes with no descendant at the bottom of the tree, representing extant taxa. In SnappNet’s model, each network branch has both its own length, measured in expected number of mutations, and its own scaled population size.

**Figure 1.**
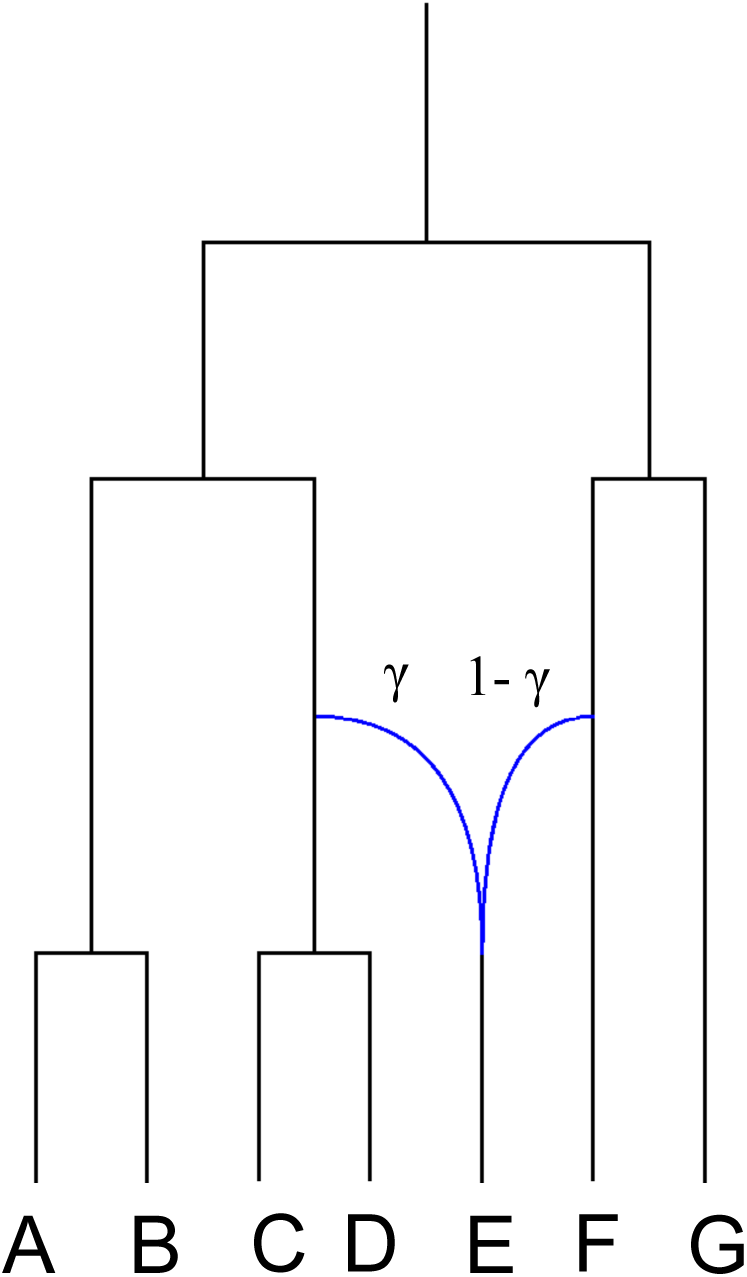
A phylogenetic network including 7 species denoted A through G. Branches are directed from top to bottom: they go forward in time. The oldest ancestor is represented at the top, and the the leaves are represented at the bottom. Each branch has its own length measured in expected number of mutations and has its own scaled population size. The reticulation node is the internal node with one outgoing branch leading to species E and two incoming edges in blue. Above the latter, *γ* and 1 *− γ* indicate the inheritance probabilities from these two ancestors for the individuals in the outgoing branch.

#### 2.1.2 Gene tree model

For each site, the associated gene tree is obtained according to the network multispecies coalescent (NMSC) model. The process starts at the leaves of the phylogenetic network and goes backward in time, <u>i.e.</u>, up the network, until all lineages coalesce. At the beginning, in the branches rising up from the leaves, coalescence occurs only between lineages that belong to the same species. Two given lineages coalesce at rate 2*µ/θ* where *µ* and *θ* refer respectively to a mutation rate (cf. Section 2.1.3 below) and to the scaled population size parameter of that species. Assuming that *k* lineages belong to that species, the first coalescent time follows an exponential distribution with parameter *k*(*k −* 1)*µ/θ*, since the coalescence of each combination of 2 lineages is equiprobable. When *k* = 2, the expected coalescent time is *θ/*(2*µ*), *θ* then being the average number of mutations separating two individuals. When the lineages (that fail to coalesce) enter a speciation node, the remaining lineages of the two species are considered as belonging to a same population. Then, they are allowed to coalesce within the branch ancestor to that speciation node according to the *θ* value associated to that branch. When the lineages (that fail to coalesce) enter a reticulation node, each lineage chooses independently its ancestral origin, among the two branches ancestral to that node (cf. model of [141]). Any given lineage goes either through the left reticulation branch with probability *γ* or through the right reticulation edge with probability 1 *− γ*. In what follows, *γ* will be called the inheritance probability. Last, above the root of the phylogenetic network, coalescence occurs among all lineages and the process ends when only one ancestral lineage remains.

Note that as its cousin Snapp [21], SnappNet considers *µ* = 1 (see the explanations below).

#### 2.1.3 Mutation model

As in Snapp, SnappNet considers biallelic markers, with *red* and *green* colors representing the two alleles. Markers evolve along the gene tree branches, according to a continuous time Markov chain. In this context, let *u* and *v* denote the instantaneous mutation rates from red to green, and from green to red respectively. Under this model, on a branch of length *T*, there are on average 2*uvT/*(*u* + *v*) mutations. Then, the mutation rate *µ* on the branch equals 2*uv/*(*u* + *v*). Imposing the constraint 2*uv/*(*u* + *v*) = 1 (i.e. *µ* = 1), enables to measure branch lengths in substitutions per site (i.e. genetic distance).

### 2.2 Real data

We sampled from the largest sources of data [61, 130] (see Figure 2A). We refer to potential wild ancestors as components of *O. rufipogon*, in line with [51]. The three included components are named Or3, Or1I and Or1A and are closely related to japonica, indica and aus, respectively. They correspond respectively to Or-IIIa, Or-Ia and Or-1b in [51]. For denoting cultivated materials we use the classification scheme proposed in [130] and we use the terms Jap (for GJ, Geng-Japonica), Ind (for XI, Xien-Indica), cAus (for *circum*Aus) and cBas (for *circum*Basmati). In view of the proven complexity of rice domestication, we pay specific attention to the representation of taxonomic units, by considering different representatives for each unit.

**Figure 2.**
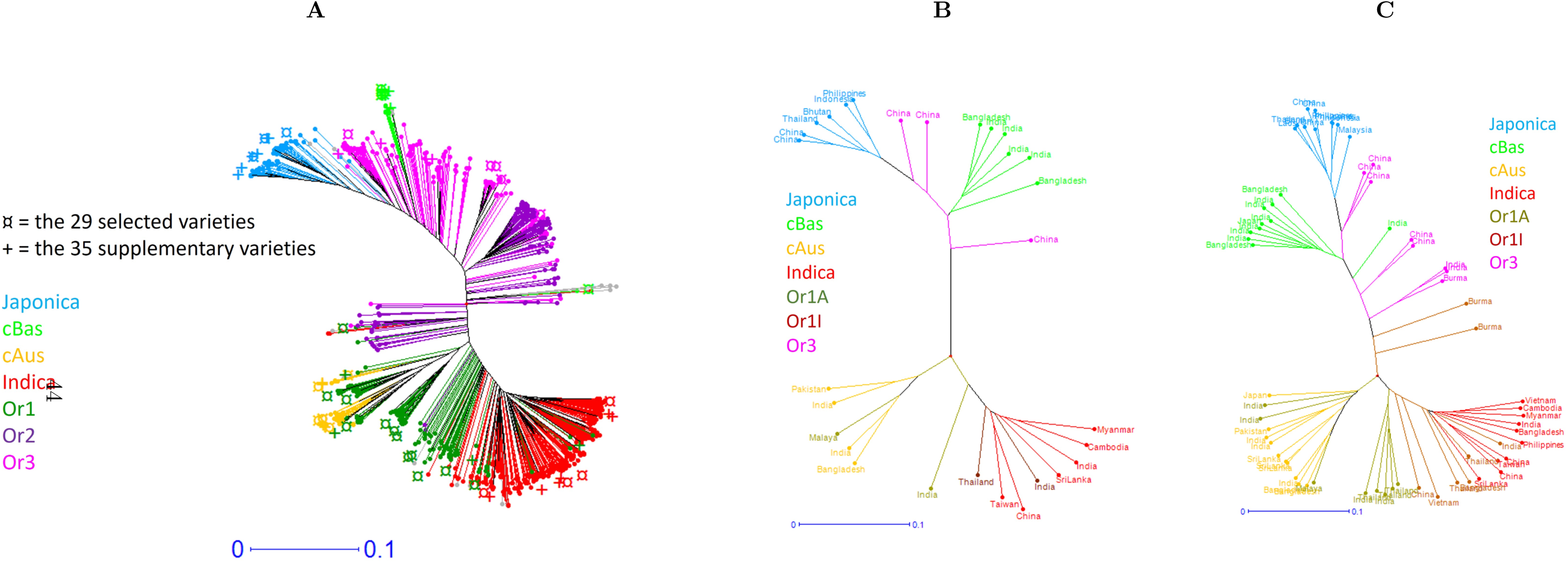
Summary of rice molecular diversity used for selecting our sample of rice cultivated varieties and wild types. A: unweighted neighbour joining (UWNJ) tree reflecting dissimilarities among 899 accessions based on 2.48 million SNPs as described in [130]; the accessions are colored according to their classification into wild population types or cultivar groups. B: UWNJ tree using the same data for the 29 accessions we selected. C: UWNJ tree using the same data for the 64 accessions we selected.

First, we considered 29 varieties (cf. Table 1 in Supplementary Material) that are either representative of the four main rice cultivar subpopulations (Jap, Ind, cAus and cBas) or representatives of the three wild subpopulations (Or3, Or1I and Or1A), considered as progenitors of cultivated rice. We refer the reader to Figure 2B and also to Figure 1B in Supplementary Material for more details. We built several random data sets from the same set of varieties (cf. Table 1 in Supplementary Material), in order to explore the diversity among the varieties. Data sets 1-17 contain only one variety per subpopulation (see Table 2 in Supplementary Material) and were analyzed in the first inference step we detailed previously. Data sets 18-21 contain either two or three varieties per subpopulation (see Tables 3 and 4 in Supplementary Material) and served as a basis for the second step of our inference process. Last, data set 22 was formed by adding 35 varieties of diverse origin to those of the previous data sets, reaching a total of 64 varieties (cf. Figure 2C and also in Supplementary material Figure 1C and Tables 5 and 6). In more details, data set 22 includes 10 varieties for each of the four cultivar subpopulations and 8 varieties for each of the three wild subpopulations. When creating this data set, we made sure to diversify the geographic origin of the Or3 samples, extending them to Myanmar (Burma) and India.

**Table 1.** Informations obtained according to the Tracer software, when data set 4 was analyzed with SnappNet. Two different samplings of 12,000 SNPs were considered, and also two chains for each sampling.

|  |  | First Sampling |  | Second Sampling |  |
| --- | --- | --- | --- | --- | --- |
|  |  | Chain 1 | Chain 2 | Chain 1 | Chain 2 |
| **LogPosterior** | mean | -23527.3838 | -23527.3273 | -23743.1762 | -23741.4521 |
|  | stdev | 6.3969 | 6.4014 | 6.2398 | 6.1792 |
|  | median | -23526.8828 | -23526.7721 | -23742.6857 | -23740.9926 |
|  | auto-correlation time | 17435.1626 | 11035.5755 | 14964.0538 | 12787.8975 |
|  | effective sample size | 516.3 | 815.6 | 9001 | 9001 |
| **LogLikelihood** | mean | -23338.6348 | -23338.2496 | -23553.0738 | -23551.7665 |
|  | stdev | 4.408 | 5.2966 | 2.5575 | 2.8963 |
|  | median | -23338.7152 | -23338.5423 | -23552.8238 | -23551.6634 |
|  | auto-correlation time | 19298.6346 | 33317.1566 | 21328.0454 | 24161.7779 |
|  | effective sample size | 466.4 | 270.2 | 422 | 372.5 |
| **LogPrior** | mean | -188.749 | -189.0777 | -190.1024 | -189.6856 |
|  | stdev | 6.2051 | 6.6087 | 5.6346 | 5.4366 |
|  | median | -188.0584 | -188.2521 | -189.5716 | -189.2066 |
|  | auto-correlation time | 14740.2916 | 21491.0215 | 8145.1581 | 12655.6205 |
|  | effective sample size | 610.6 | 418.8 | 1105.1 | 711.2 |
| ***u*** | mean | 0.5545 | 0.5545 | 0.5559 | 0.5562 |
|  | stdev | 9.0107E-4 | 8.8289E-4 | 8.9439E-4 | 9.1308E-4 |
|  | median | 0.5545 | 0.5545 | 0.5559 | 0.5562 |
|  | auto-correlation time | 2219.589 | 2212.8031 | 2124.4036 | 2444.8495 |
|  | effective sample size | 9001 | 4067.7 | 4237 | 3681.6 |
| ***v*** | mean | 5.089 | 5.0886 | 4.9753 | 4.9508 |
|  | stdev | 0.0759 | 0.0743 | 0.0717 | 0.0723 |
|  | median | 5.0879 | 5.0886 | 4.976 | 4.9493 |
|  | auto-correlation time | 2224.845 | 2215.9913 | 2125.5838 | 2452.4402 |
|  | effective sample size | 4045.7 | 4061.8 | 4234.6 | 3670.2 |
| ***d*** | mean | 5.7257 | 5.7197 | 5.6741 | 5.4883 |
|  | stdev | 2.7562 | 2.829 | 2.679 | 2.6499 |
|  | median | 5.3521 | 5.3404 | 5.3281 | 5.1385 |
|  | auto-correlation time | 1622.6145 | 1729.5125 | 1393.50 | 1258.5231 |
|  | effective sample size | 5547.2 | 5204.4 | 6459.3 | 7152 |
| ***r*** | mean | 0.234 | 0.2381 | 0.2233 | 0.241 |
|  | stdev | 0.1647 | 0.1703 | 0.1607 | 0.1686 |
|  | median | 0.1965 | 0.1974 | 0.184 | 0.2023 |
|  | auto-correlation time | 2022.0597 | 2135.5715 | 1674.1636 | 2123.6676 |
|  | effective sample size | 4451.4 | 4214.8 | 5376.4 | 4238.4 |

**Table 2.** Scanwidth and average number of iterations in six months of computation as a function of the network under study. The 16 networks are illustrated in Figures 3.

| Network | Scanwidth | Average number of iterations |
| --- | --- | --- |
| 1 | 2 | 692,153 |
| 2 | 2 | 579,543 |
| 3 | 2 | 612,413 |
| 4 | 2 | 1,688,528 |
| 5 | 2 | 2,030,07 |
| 6 | 2 | 397,061 |
| 7 | 2 | 3,338,825 |
| 8 | 3 | 15,666 |
| 9 | 3 | 28,795 |
| 10 | 3 | 16,292 |
| 11 | 3 | 102,125 |
| 12 | 3 | 14,651 |
| 13 | 3 | 51,700 |
| 14 | 3 | 134,270 |
| 15 | 3 | 303,303 |
| 16 | 3 | 56,913 |

**Table 3.**
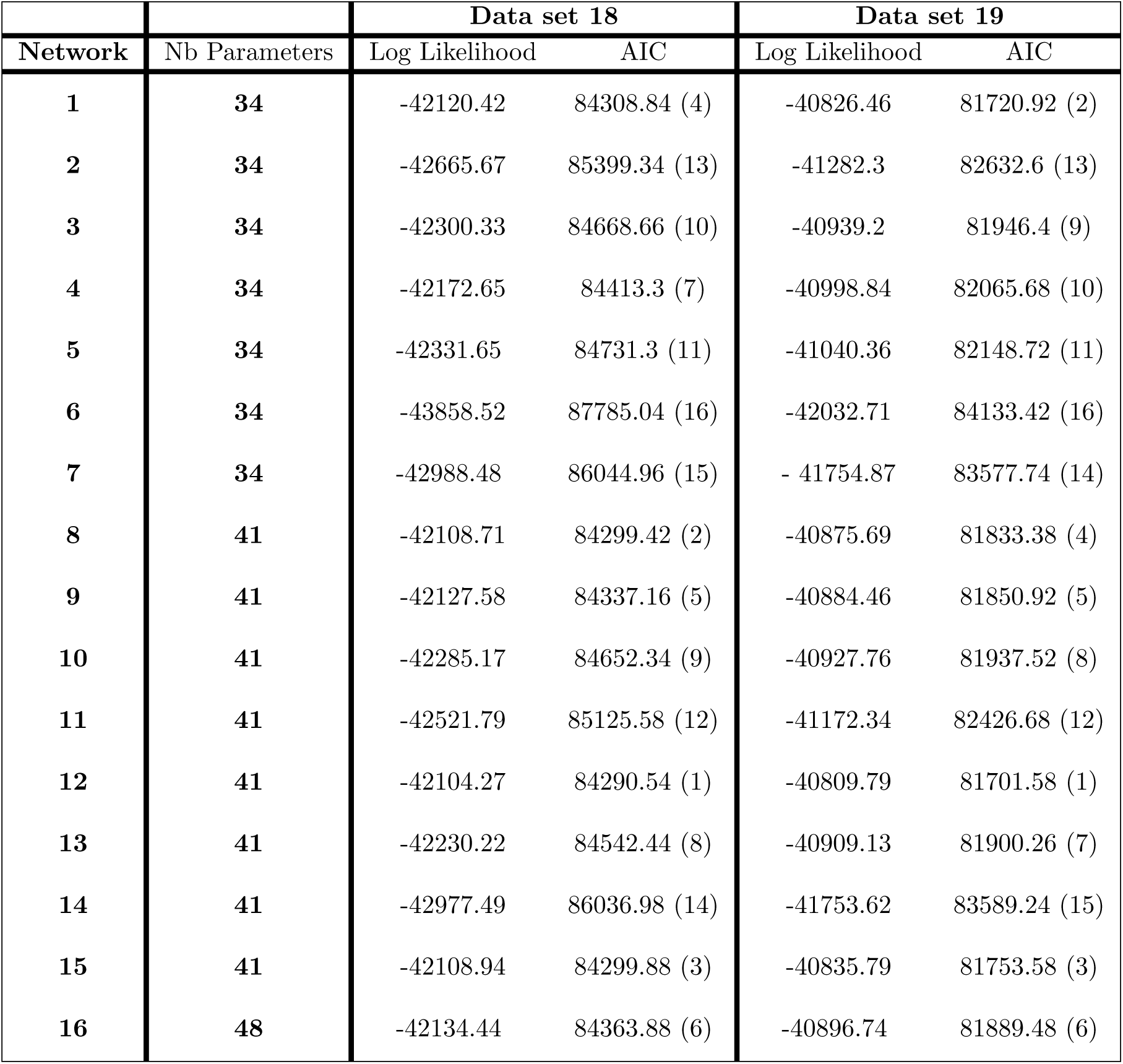
Log Likelihood and AIC values, computed on data sets 18 and 19 (each comprising m=12,000 SNPs distributed across the rice genome) for the 16 phylogenetic networks described in Figure 3. The network ranking, given into brackets, is obtained by minimizing the AIC criterion.

| Network | Nb Parameters | Data set 18 |  | Data set 19 |  |
| --- | --- | --- | --- | --- | --- |
|  |  | Log Likelihood | AIC | Log Likelihood | AIC |
| 1 | 34 | -42120.42 | 84308.84 (4) | -40826.46 | 81720.92 (2) |
| 2 | 34 | -42665.67 | 85399.34 (13) | -41282.3 | 82632.6 (13) |
| 3 | 34 | -42300.33 | 84668.66 (10) | -40939.2 | 81946.4 (9) |
| 4 | 34 | -42172.65 | 84413.3 (7) | -40998.84 | 82065.68 (10) |
| 5 | 34 | -42331.65 | 84731.3 (11) | -41040.36 | 82148.72 (11) |
| 6 | 34 | -43858.52 | 87785.04 (16) | -42032.71 | 84133.42 (16) |
| 7 | 34 | -42988.48 | 86044.96 (15) | - 41754.87 | 83577.74 (14) |
| 8 | 41 | -42108.71 | 84299.42 (2) | -40875.69 | 81833.38 (4) |
| 9 | 41 | -42127.58 | 84337.16 (5) | -40884.46 | 81850.92 (5) |
| 10 | 41 | -42285.17 | 84652.34 (9) | -40927.76 | 81937.52 (8) |
| 11 | 41 | -42521.79 | 85125.58 (12) | -41172.34 | 82426.68 (12) |
| 12 | 41 | -42104.27 | 84290.54 (1) | -40809.79 | 81701.58 (1) |
| 13 | 41 | -42230.22 | 84542.44 (8) | -40909.13 | 81900.26 (7) |
| 14 | 41 | -42977.49 | 86036.98 (14) | -41753.62 | 83589.24 (15) |
| 15 | 41 | -42108.94 | 84299.88 (3) | -40835.79 | 81753.58 (3) |
| 16 | 48 | -42134.44 | 84363.88 (6) | -40896.74 | 81889.48 (6) |

**Table 4.**
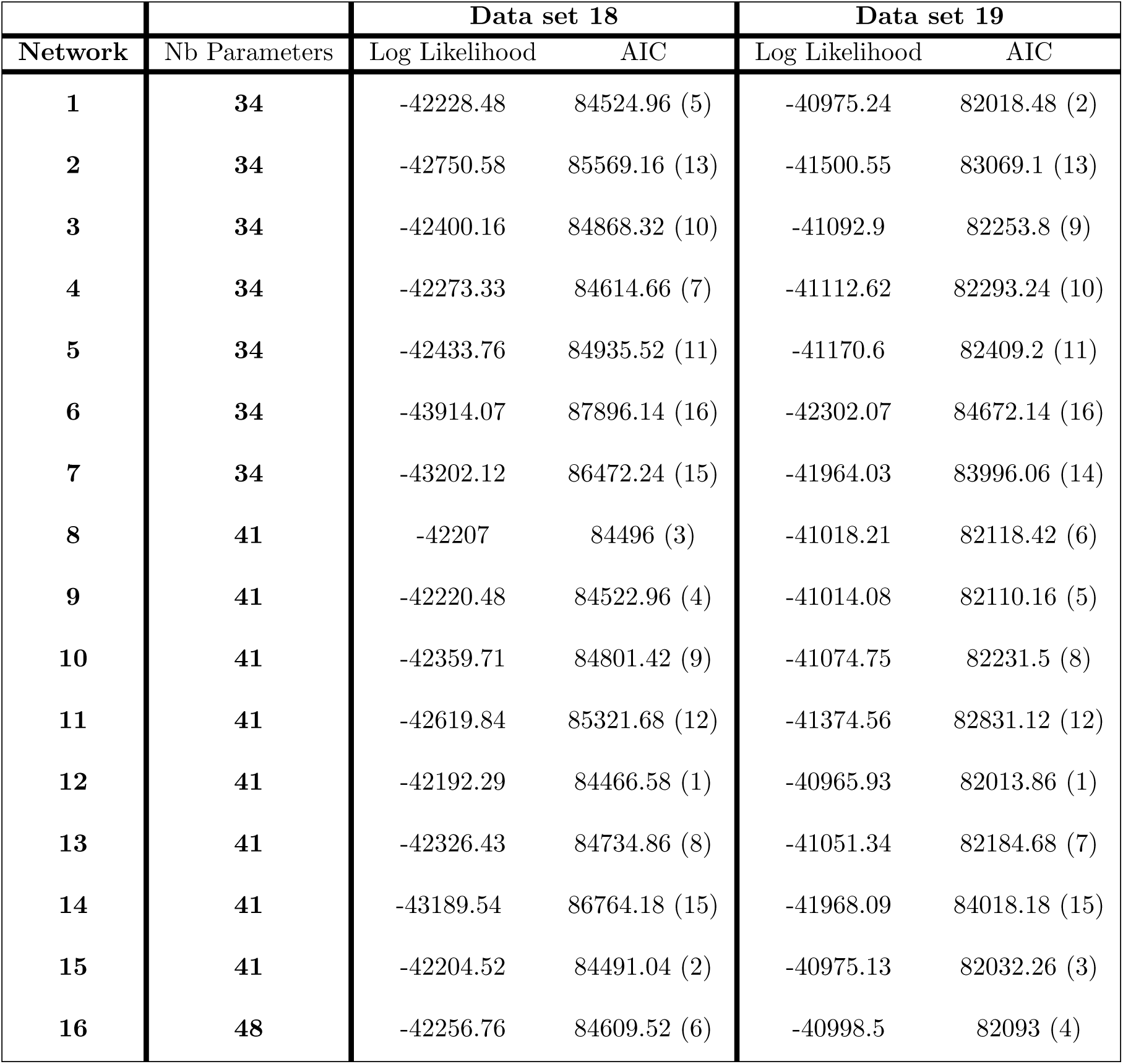
Same as Table 3 except that the analysis relies on m=12,000 other SNPs sampled along the genome. The selected rice varieties remain unchanged for the two data sets.

**Table 5.** Log Likelihood and AIC evaluations, based on data sets 20 and 21, for the 16 phylogenetic networks described in Figure 3. The network ranking, given into brackets, is obtained by minimizing the AIC criterion. The two data sets consist in m=12,000 SNPs spread out along the rice genome.

| Network | Nb Parameters | Data set 20 |  | Data set 21 |  |
| --- | --- | --- | --- | --- | --- |
|  |  | Log Likelihood | AIC | Log Likelihood | AIC |
| 1 | 34 | -41516.49 | 83100.98 (4) | -41094.29 | 82256.58 (2) |
| 2 | 34 | -42036.80 | 84141.60 (13) | -41532.51 | 83133.02 (13) |
| 3 | 34 | -41713.1 | 83494.20 (10) | -41197.32 | 82462.64 (8) |
| 4 | 34 | -41590.45 | 83248.9 (7) | -41225.04 | 82518.08 (9) |
| 5 | 34 | -41777.74 | 83623.48 (11) | -41267.18 | 82863.06 (11) |
| 6 | 34 | -43210.17 | 86488.34 (16) | -42121.69 | 84311.38 (16) |
| 7 | 34 | -42368.11 | 84808.22 (15) | -42073.26 | 84214.52 (15) |
| 8 | 41 | -41492.67 | 83067.34 (2) | -41122.56 | 82327.12 (3) |
| 9 | 41 | -41509.35 | 83100.70 (3) | -41126.92 | 82335.84 (5) |
| 10 | 41 | -41687.63 | 83457.26 (9) | -41165.59 | 82413.18 (7) |
| 11 | 41 | -41871.39 | 83824.78 (12) | -41443.48 | 82968.96 (12) |
| 12 | 41 | -41488.59 | 83059.18 (1) | -41083.76 | 82249.52 (1) |
| 13 | 41 | -41657.74 | 83397.48 (8) | -41382.7 | 82847.4 (10) |
| 14 | 41 | -42367.97 | 84817.94 (15) | -42050.02 | 84182.04 (14) |
| 15 | 41 | -41514.53 | 83111.06 (5) | -41125.53 | 82333.06 (4) |
| 16 | 48 | -41559.09 | 83214.18 (6) | -41137.33 | 82370.66 (6) |

**Table 6.** Same as Table 5 except that the analysis relies on m=12,000 other SNPs sampled along the genome. The selected rice varieties remain unchanged for the two data sets.

| Network | Nb Parameters | Data set 20 |  | Data set 21 |  |
| --- | --- | --- | --- | --- | --- |
|  |  | Log Likelihood | AIC | Log Likelihood | AIC |
| 1 | 34 | -41606.31 | 83280.62 (5) | -41210.38 | 82488.76 (2) |
| 2 | 34 | -42125.59 | 84319.18 (13) | -41727.88 | 83523.76 (13) |
| 3 | 34 | -41792.93 | 83653.86 (10) | -41312.23 | 82692.46 (9) |
| 4 | 34 | -41668.81 | 83405.62 (7) | -41315.54 | 82699.08 (10) |
| 5 | 34 | -41856.99 | 83781.98 (11) | -41362.86 | 82793.72 (16) |
| 6 | 34 | -43307.6 | 86683.2 (16) | -42360.3 | 84788.6 (15) |
| 7 | 34 | -42550.22 | 85168.44 (14) | -42246.05 | 84560.10 (14) |
| 8 | 41 | -41575.9 | 83233.8 (1) | -41210.05 | 82502.1 (3) |
| 9 | 41 | -41591.37 | 83264.74 (3) | -41231.13 | 82544.26 (5) |
| 10 | 41 | -41762.87 | 83607.74 (9) | -41263.86 | 82609.72 (8) |
| 11 | 41 | -41975.10 | 84032.20 (12) | -41607.8 | 83297.6 (12) |
| 12 | 41 | -41580.22 | 83242.44 (2) | -41183.84 | 82449.68 (1) |
| 13 | 41 | -41737.70 | 83557.4 (8) | -41477.76 | 83037.52 (11) |
| 14 | 41 | -42550.59 | 85183.18 (15) | -42232.43 | 84546.86 (6) |
| 15 | 41 | -41593.38 | 83268.76 (4) | -41222.17 | 82526.34 (4) |
| 16 | 48 | -41644.58 | 83385.16 (6) | -41226.79 | 82549.58 (7) |

MSAs were obtained for all chosen variety sets, by sampling 12,000 SNPs along the whole genome of rice. On each chromosome, the 1,000 SNPs were chosen as far as possible since SnappNet [101] assumes independence between sites. In order to evaluate the influence of character sampling, a second sampling of 12,000 SNPs along the whole genome was also considered for each variety set.

### 2.3 SnappNet in a Bayesian framework

Our first step consists in running SnappNet in a Bayesian setting. We concentrate on rice data sets 1-17, which contain one variety per subgroup. The same priors as those incorporated by default within SnappNet, were considered [101]. Recall that SnappNet incorporates the same network prior as in SpeciesNetwork ([145]): the network is modeled according to a birth hybridization process (see [70]). For the MCMC analysis, on each data set, two markov chains per sampling were considered. Each chain ran during 10 millions iterations. The two chains were combined after checking they add converged (discarding the first 10% observations considered as their burn-in phase). The convergence was assessed by the Effective Sample Size (ESS) criterion ([81]), thanks to the Tracer software ([104]). Recall that MCMC samples are correlated observations, and as a consequence, the ESS determines the number of independent observations that can be drawn from the sample obtained by MCMC. For all parameters, a minimal ESS value of 200 was imposed as advised by the beast community (https://beast.community/ess_tutorial). The posterior distribution was estimated from observations sampled every 1,000 iterations. To limit the computational burden, the number of reticulations was bounded by 2.

### 2.4 SnappNet in a penalized likelihood framework

#### Preliminaries

The first exploratory step revealed an important variability in scenarios obtained while widely exploring the topological space of possible networks (see Results section below).

In view of this, our second inference step focuses on a few evolutionary scenarios of interest which we analyzed in greater depth, based on large datasets. We focused on 16 networks (cf. Figure 3), each representing a different rice evolutionary scenario. These networks were built by assembling topological traits emerging from networks obtained in step one. Moreover, most of these networks reflect patterns and pathways reported in the wide range of studies on rice diversity and evolution [147, 56, 61, 140, 29, 27, 130, 53, 146, 47, 69, 138, 51, 148].

**Figure 3.**
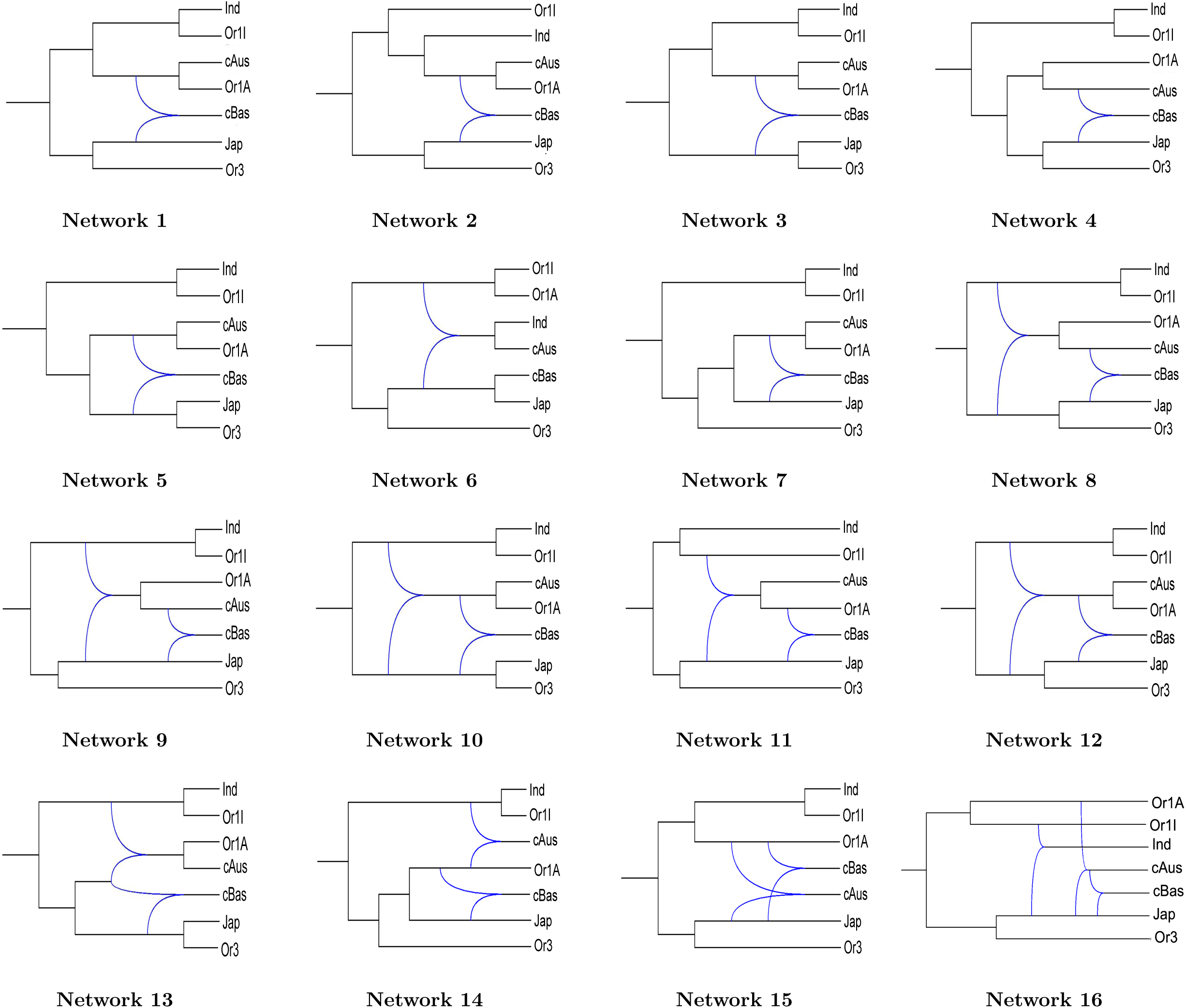
The 16 studied phylogenetic networks at step two in the rice data analysis. Each network illustrates a different rice evolutionary scenario. Networks 4, 5, 6 and 7 represent unlikely scenarios.

In particular, most of the scenarios examined in the second step are consistent with the main characteristics of the species’ genetic structure, i.e.:

- across species, a correspondence between wild subpopulations and cultivated subpopulations (Jap/Or3, Ind/Or1I, cAus/Or1A)
- among cultivars, a greater similarity between Indica and cAus on the one hand, and between Japonica and cBas on the other.
- cBas occupies an intermediate position between the Jap/Or3 and the cAus/Or1A lineages.

Note that a small number of the 16 studied networks correspond to unlikely scenarios. This allowed us to verify whether the data analysis is able to differentiate them from the most likely scenarios.

#### Main differences among the 16 networks

In a nutshell, Networks 1, 2, 3, 4, 5, 7, 8, 9, 10, 11, 12, 13, 14, 15 present multiple domestications (Japonica, Indica and cAus, or only Japonica and Indica). Most of these multiple domestication scenarios were mentioned in [56, 29, 130, 146, 69]. In contrast, Networks 6 and 16 illustrate a single domestication scenario (Japonica), as highlighted in the studies of [61, 140, 27, 53, 138, 51]. The most commonly proposed scenarios in the whole-genome surveys era are illustrated by Network 1 with three independent domestications, and by Network 16 with one domestication (Japonica from Or3) followed by hybridization with other wild forms. In contrast, four of the studied networks (Networks 4, 5, 6 and 7) depart from the structure features listed above and represent unlikely scenarios.

#### Differences in terms of hybridizations

The 16 studied networks incorporate either one, two or three hybridizations. In particular, seven networks present one hybridization whereas eight networks exhibit two hybridations. A single network (Network 16) displays three hybridations. While the most common hybridization is the one that leads to cBas, seven networks highlight the contribution of domesticated Jap to the emergence of other cultivated types (networks 6, 9, 11, 15 and 16) or even to wild type Or1A (networks 9 and 11). A few other networks explore original hypotheses. For instance, Network 13 depicts a scenario with 3 separate domestications (Japonica, Indica, cAus) and a novel hybridization. Indeed, the peri-himalayan types cAus and cBas have a hybrid origin mobilizing a cryptic wild type that left no pure descendant.

#### Differences in terms of topologies

Network 16 exhibit 3 reticulation nodes, whereas networks 8-12 present similar topologies with two reticulation nodes on top of each other. Networks 1-7 present a single reticulation node. Most of these networks are embedded within more complex networks. Indeed, Networks 1 and 4 are respectively <u>displayed</u> [65, 93] by networks 12 and 8, <u>i.e.</u>, are sub-networks of the latter. Finally, Networks 3 and 5 are both displayed by Network 10.

#### Data analysis with SnappNet

For each of the 16 evolutionary scenarios, we computed the likelihoods associated to data sets 18-21, thanks to SnappNet. The likelihood optimization was based on 11 operators among the 16 original operators incorporated within SnappNet. Indeed, since we kept the network topology as fixed, we did not consider the 5 topological operators originating from [144]. The procedure used to optimize the likelihood is described in Section 4 of SnappNet’s guide at https://github.com/rabier/MySnappNet/tree/master/workspace-Package-Beast/SnappNet/doc. We let the optimization run during six months for each of the 16 networks under study. Note that SnappNet is only used here to compute likelihoods of fixed networks. For datasets as large as those studied here it is not feasible to explore networks using the full SnappNet Bayesian inference process. This would also be impossible with another current method such as MCMCBiMarkers [149].

As in [116], to penalize models with more parameters, the AIC [1] criterion was adopted using the following expression: AIC = 2*p −* 2 ln(L) where *p* refers to the number of parameters.

### 2.5 The SNARF approach

Step two lead us to select six phylogenetic networks, namely Networks 12, 8, 1, 15, 9 and 16, that we study in more details in Step three. Recall that each network, selected among the 16 networks according to the AIC criterion, represents the evolution of 7 rice subpopulations.

The third inference step on rice data involves a combination of the SnappNet method and a new implementation of the ABC-RF framework, dedicated to the inference of phylogenetic networks. This approach, that we call Snarf, is general in the sense that it is applicable to other organisms than rice.

Before describing the priors and the summary statistics (SS), it is necessary to highlight the fact that this hybrid approach was always built on SnappNet in our experiments, in the sense that all investigated network had branch lengths estimated by SnappNet. The importance of branch lengths in ABC approaches was shown in [115]. Note also that instantaneous rates *u* were always calibrated thanks to SnappNet. Details are given below.

#### 2.5.1 Priors

In this study, we assigned a dedicated prior to (a) each inheritance probability associated to a reticulation node and to (b) each scaled population size *θ* associated to a branch. Two kinds of priors were considered for each *γ*: the prior is either (a) specific or (b) general. The specific prior relies on SnappNet, whereas the general prior is obtained independently from SnappNet. For the specific prior, the inheritance probability was drawn from a uniform distribution *U* (max(0.05*, γ̃ − δ*), min(0.95*, γ̃* + *δ*)), where *γ̃* denotes the maximum likelihood estimate (MLE) of the inheritance probability obtained with SnappNet, and where *δ* refers to a small real value. This prior allows to avoid nested models. In practice, *δ* was set to 0.1 in all experiments. In contrast, the general prior consists in sampling the inheritance probability from a uniform distribution *U* (0.05, 0.95).

In the same way, we investigated two kinds of population size priors. Under the first prior, for each network branch, the scaled population size *θ* was drawn independently from a gamma distribution, Γ(1*, θ̃*). Here, *θ̃* refers to the MLE from SnappNet in the prior step considering fixed topology networks. For instance, on rice data, the estimates where computed from data sets 18-21. For the second prior, the scaled population sizes *θ* were drawn from Γ(1, 0.1) distributions, like in Snapp.

As prior for the instantaneous rate *u*, we concentrated exclusively on a uniform distribution *U* (max(0*, ũ −* 0.0005)*, ũ* + 0.0005) where *ũ* refers to MLE of the rate obtained with SnappNet (cf. Table 7 in Supplementary Material). We also studied the possibility to use no prior for *u* at all: in this case, we directly set *u* to the MLE value *ũ*. Last, for changing the branch lengths, a specific algorithm, inspired from [144], was considered. It is described as Algorithm 1 in Supplementary Material. As a starting point, branch lengths were set to MLE values from SnappNet. Table 16 in Supplementary Material summarizes all the prior combinations investigated in our study.

**Table 7.** Rankings for each of the 16 different networks illustrating a different rice evolution scenario. The criterion is the sum over the rankings obtained for the 8 datasets (cf. rankings in Tables 3, 4, 5 and 6). The final ranking is given into brackets.

| Network | Sum of all rankings |
| --- | --- |
| 12 | 9 (1) |
| 8 | 24 (2) |
| 1 | 26 (3) |
| 15 | 28 (4) |
| 9 | 35 (5) |
| 16 | 47 (6) |
| 10 | 67 (7) |
| 13 | 67 (8) |
| 4 | 67 (9) |
| 3 | 75 (10) |
| 5 | 93 (11) |
| 11 | 96 (12) |
| 2 | 104 (13) |
| 14 | 109 (14) |
| 7 | 116 (15) |
| 6 | 127 (16) |

#### 2.5.2 Summary statistics

Recall that ABC-RF relies on simulated data, that are compared to observed data thanks to SS (see [85]). In this study, we propose three kinds of SS, all made available within the SimSnappNet simulator (https://github.com/rabier/SNARF).

The first kind of SS we consider (denoted SS 1-5) represents basic statistics inspired by population genetics, but that can be adapted when dealing with multiple species:

1. **Average number of pairwise differences** between the sequences of two individuals sampled **from all species**. This average is computed over all sites and all combinations of two individuals.
2. **Average number of pairwise differences** between two individuals sampled **within a given species**. This average is computed over all the sites and over all combinations of two individuals from the species of interest.
3. **Average number of pairwise differences between** one individual from **a given species**, **and** one individual from **all other species**. This average is computed over all the sites and over all combinations of two individuals.
4. **Average** linkage disequilibrium (**LD**) over consecutive sites and **over all species**. Pearson correlation was used as the LD measure.
5. **Average** linkage disequilibrium (**LD**) over consecutive sites, **for a given species**. Pearson correlation was used as the LD measure.

Our second set of SS relies on the theoretical work of Kubatko et al. on phylogenetic invariants [77]. These authors proposed statistics whose analytical expression was obtained under the model described in [89, 75]. In our case, data will be generated with SimSnappNet, according to SnappNet’s model. This model discrepancy is of no consequence in ABC-RF. Recall that the key point is to get a table containing a large panel of summary statistics allowing to compare observed data with simulated data. We denote SS 6-18 our statistics inspired by those of Kubatko et al.

For describing this set of SS, let us focus on the phylogenetic network represented on the left side of Figure 4 (a reproduction of [77, Figure 1]). This network is composed of **4 species** (named *O*, *P*_1_, *H*, *P*_2_) and only one reticulation event. Assuming only one lineage per species, let *p_ijkl_*denote the probability P(*O* = *i, P*_1_ = *j, H* = *k, P*_2_ = *l*) at a given site, where (*i, j, k, l*) *∈ {−*1, 1*}*^4^. The *−*1 and +1 values respectively refer to the green and red alleles in SnappNet’s model. The assignment (*i, j, k, l*) to the tips of the network can be viewed as a site pattern (see [77]).

**Figure 4.**
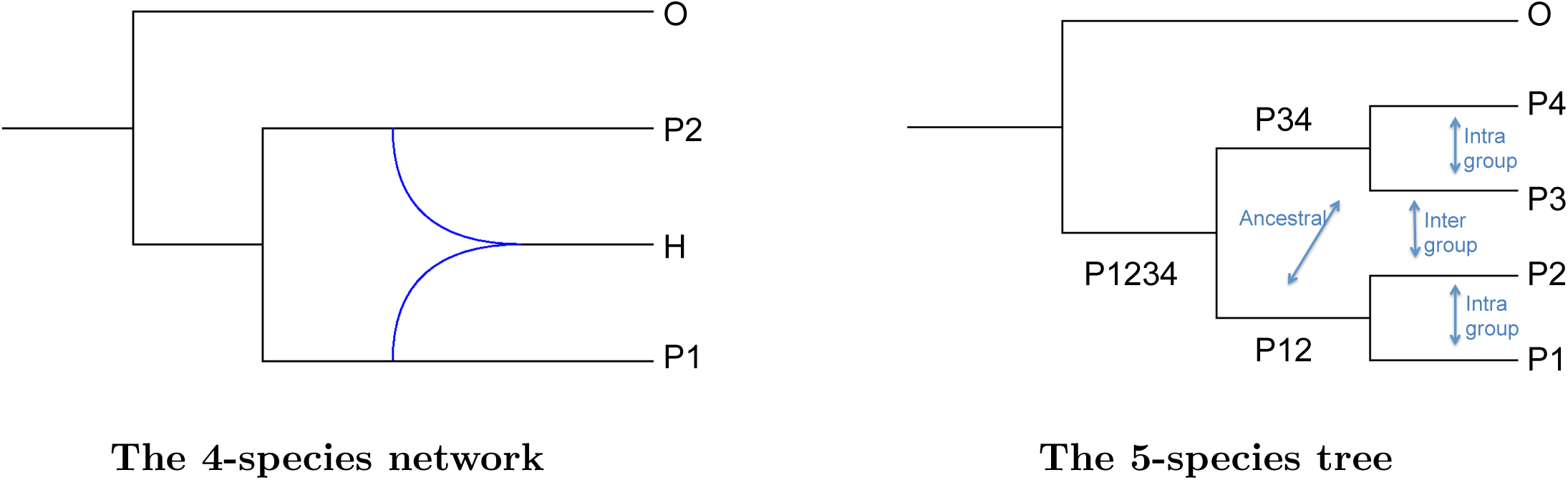
The two phylogenetic networks used for computing SS based on [77] and [97]. On the left-side, the network matches Figure 1 of [77], whereas on the right-side, it coincides with Figure 1b of [97].

Let *f*_1_, *f*_2_, *f*_3_, *f*_4_ be the quantities defined in the following way:

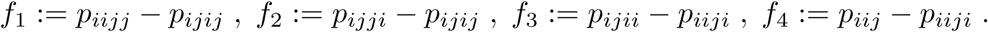

In the same way, *f*^^^_1_, *f*^^^_2_, *f*^^^_3_, *f*^^^_4_ refer to the following estimators:

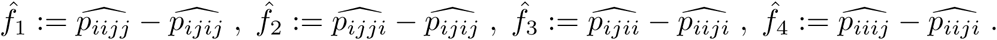

where 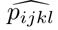 refers to the average over the sites for the site pattern (*i, j, k, l*). Note that we kept here the notations of Kubatko et al. [77]. Given the above definitions we can now precise SS 6-18:

6. **D-statistic by [48, 96]** (cf. equation 1 of [15]), see also [108, 34])

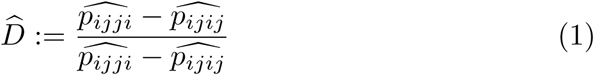

7. **an estimator** 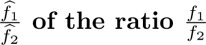 (cf. equation 2 of [15])

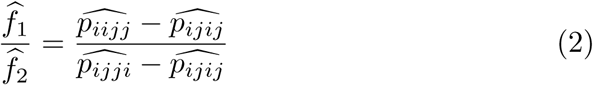

8. **an estimator** 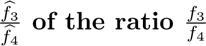 **(cf.equation 5of [77])**

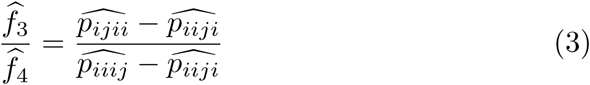

9. **an estimator** 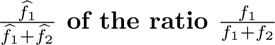 (cf. **equation 3** of [15])

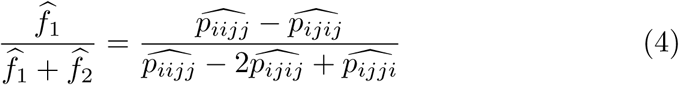

10. **an estimator** 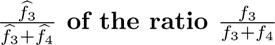

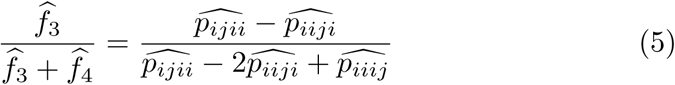

11. **an estimator** 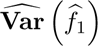 **of Var** (*f*_1_) (cf. equation 9 of [77])

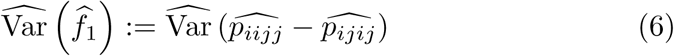

12. **an estimator** 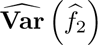 **of Var** (*f*_2_) (cf. equation 10 of [77])

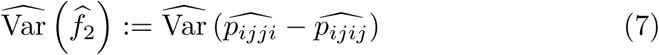

13. **an estimator** 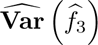 **of Var** (*f*_3_)

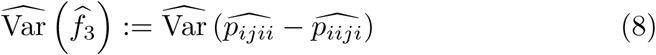

14. **an estimator** 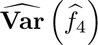 **of Var** (*f*_4_)

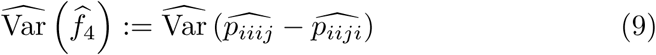

15. **an estimator** 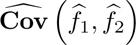 of Cov (*f_1_, f_2_*) (cf. above equation 11 of [77])

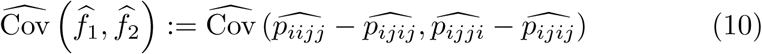

16. **an estimator** 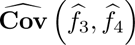 **of Cov** (*f*_3_*, f*_4_)

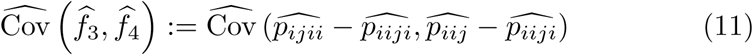

17. the Hils statistic based on *f*_1_ and *f*_2_**, under the null hypothesis of no hybridization** (cf. equation 13 of [77])

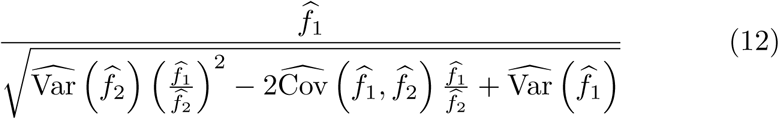

18. the Hils statistic based on *f*_3_ and *f*_4_**, under the null hypothesis of no hybridization** (cf. below equation 13 of [77])

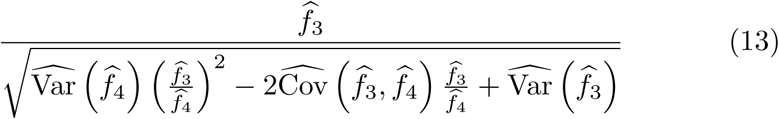

Last, our third kind of SS is motivated by the study on invariants by Pease and Hahn [97], where these authors test the presence of introgressions in a symmetric five-species phylogeny. We consider here four SS (SS 19-22) matching exactly the statistics *D_F O_*, *D_IL_*, *D_F I_*, *D_OL_*introduced in their paper.

In order to discuss briefly the “DFOIL” concept, let us consider the species tree illustrated on the right-side of Figure 4, with **5 species** named *P*_1_, *P*_2_, *P*_3_, *P*_4_ and *O*. Pease and Hahn show that, with the help of their 4 statistics and using the respective sign of these statistics, they are able to detect eight “inter-group” possible introgressions (*P*_1_ *⇒ P*_3_, *P*_3_ *⇒ P*_1_, *P*_1_ *⇒ P*_4_, *P*_4_ *⇒ P*_1_, *P*_2_ *⇒ P*_3_, *P*_3_ *⇒ P*_2_, *P*_2_ *⇒ P*_4_, *P*_4_ *⇒ P*_2_), four “intra-group” introgressions (*P*_1_ *⇒ P*_2_, *P*_2_ *⇒ P*_1_, *P*_3_ *⇒ P*_4_, *P*_4_ *⇒ P*_3_), and four “ancestral” introgressions (*P*_12_ *⇒ P*_3_, *P*_3_ *⇒ P*_12_, *P*_12_ *⇒ P*_4_).

Assuming one lineage per species, let us define *p_ijklm_* as the probability P(*P*_1_ = *i, P*_2_ = *j, P*_3_ = *k, P*_4_ = *l, O* = *m*) at a given site, where (*i, j, k, l, m*) *∈ {−*1, 1*}*^5^. 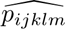 will be the average of the empirical proportions over the sites for the pattern (*i, j, k, l, m*). Although the statistics *D_F O_*, *D_IL_*, *D_F I_*, *D_OL_* are based on Pease and Hahn’s model [97], our SS 19-22 rely on patterns observed in simulated data based on SnappNet’s model.

As mentioned before, these SS do not have to be computed under the true model to be relevant for ABC-RF (cf. Section 3c).

In this context, SS 19-22 are the following:

19. **an estimator** *D̂_F O_* **of the** *D_F O_* statistic (cf. equation 2 of [97])

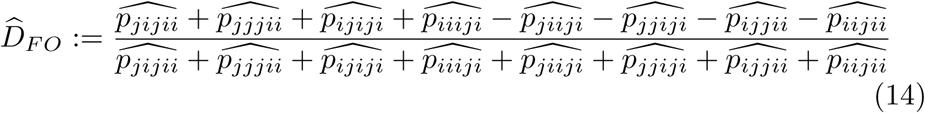

20. **an estimator** *D̂_IL_* **of the** *D_IL_* statistic (cf. equation 3 of [97])

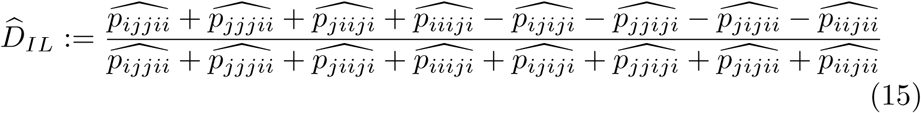

21. **an estimator** *D̂_F I_* **of the** *D_F I_* statistic (cf. equation 4 of [97])

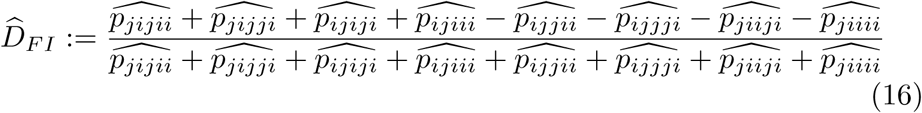

22. **an estimator** *D̂f_OL_* **of the** *D_OL_* statistic (cf. equation 5 of [97])

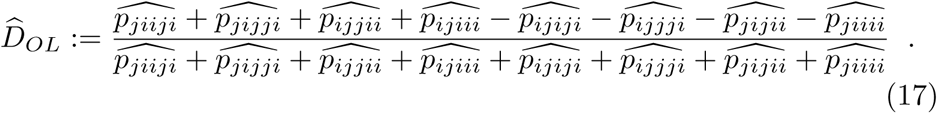

Recall that six phylogenetic networks result from step two of our analysis of rice data, namely Networks 12, 8, 1, 15, 9 and 16, each of them spanning 7 species. This leads to an overall total of 562 SS. More specifically, we considered one SS 1, seven kinds of SS 2, seven kinds of SS 3, one SS 4, seven kinds of SS 5, 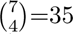 (<u>i.e.</u>, all unordered sets of 4 species out of 7) kinds of SS 6-18 as well as 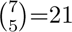 kinds of of SS 19-22.

In order to limit the computational burden for SS 6-18 (based on Kubatko and Chifman [77]), a procedure was adopted. Indeed, Data set 22 can be viewed as a large data set with 10 varieties for each of the 4 cultivar subpopulations and 8 varieties for each of the three wild subpopulations.

Once the 4 species were chosen among the 7 considered species (i.e. 4 cultivar subpopulations and 3 wild subpopulations), we considered a unique assignment of these species to species *O*, *P*_1_, *H*, *P*_2_ of the 4-species network in Figure 4. The same assignment was considered for the 16 networks of interest.

Last, for each assignment, we focused on 5^4^ = 625 random sets of 4 varieties representing each species. Recall that we have 10 (resp. 8) varieties available for the cultivar (resp. wild) species.

We proceeded in the same way for computing SS 19-22 (based on the work of Pease and Hahn [97]): once the 5 species were chosen among the 7 considered species, we considered a single assignment of the 5 species at stake to *O*, *P*_1_, *P*_2_, *P*_3_ and *P*_4_ of the 5 species tree in Figure 4. For each assignment, we focused on 5^5^ = 3125 random sets of 5 varieties. To sum up, the ratio between our sampling and all possibilities was found to be equal to 3.01% for SS 6-18 and to 1.66% for SS 19-22 (cf. Section 4 in Supplementary Material). This sampling strategy allows to reduce considerably the computation burden and gave satisfactory results for our classifier (cf. Section 3c).

#### 2.5.3 Training the Random Forest

For each of the six phylogenetic networks studied at Step 3, we simulated 14,000 data sets. As a consequence, the reference table contains *N*_Ref_ = 84,000 rows. Recall that the number of columns matches the total number of SS, <u>i.e.</u>, 562 SS in our study. Recall that RF consists in aggregating results of multiple randomized CART trees. To build each such tree, a bootstrap sample of size *N*_boot_ = *N*_Ref_ was taken from the reference table, as advised in [100].

The size *n*_try_ of the random sample of variables at each tree node was chosen to be the square root of the total number of SS (cf. package abcrf from Pudlo et al. [100]). The out of bag error rate, also named prior error rate, was computed to evaluate the abcrf algorithm performances. As usual, the majority vote was adopted for classifying scenarios. Overall, the forest contained 1,000 randomized CART trees since the prior error rate stabilized for this number of trees.

## 3 Results

In what follows, we detail results of the three steps of our study on rice data, namely:

(1) The free exploration of the topology space with SnappNet.
(2) The use of SnappNet in a penalized likelihood framework on 16 evolutionary scenarios deduced from Step (1).
(3) The discrimination between the 6 most likely scenarios found at Step (2) thanks to our new hybrid approach, Snarf, relying on ABC-RF and on SnappNet’s estimates.

### (1) Free exploration of the network topology space with SnappNet

SnappNet first ran on data sets 1-17, containing one variety per subpopulation. Two samplings of 12,000 markers were considered for each data set. Figures 5-8 illustrate, the Maximum A Posteriori (MAP) phylogenetic networks obtained by SnappNet. Branch lengths are given in units of expected number of mutations per site. Figures 2-18 in Supplementary Material report for the seventeen data sets the main network topologies sampled by SnappNet during the MCMC steps. An example of MCMC convergence assessment is given for data set 4 in Figure 9 and in Table 1: on the second sampling, the two chains presented an ESS of 9,001, illustrating a very good convergence to the stationary distribution.

**Figure 5.**
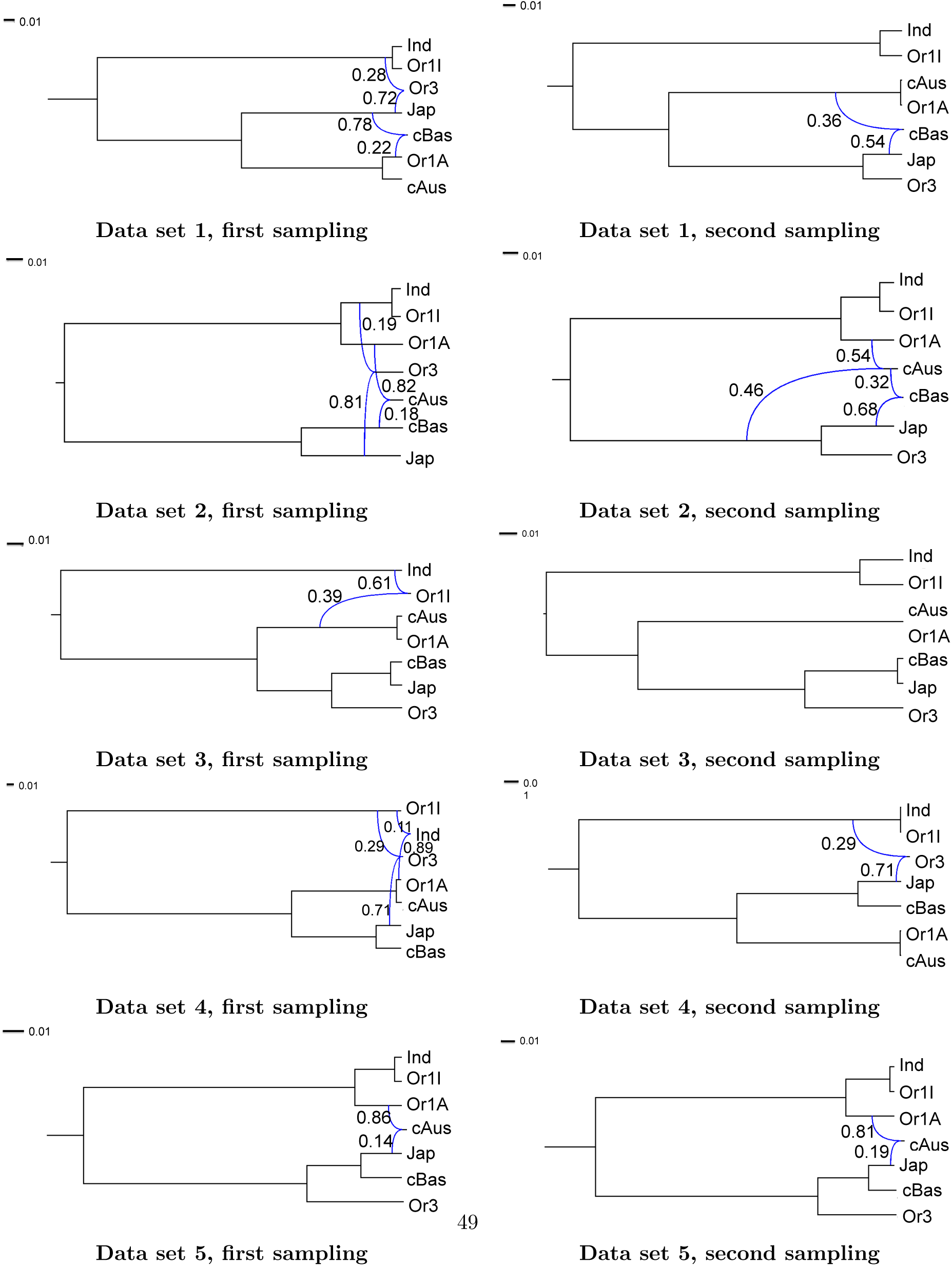
The MAP phylogenetic networks obtained for data sets 1-5 with one variety per subpopulation. The two different samplings of 12k markers along the rice genome are considered here. Inheritance probabilities are reported above reticulation edges and branch lengths are given in units of expected number of mutations per site (see the scale at the top left).

We observed a large panel of scenarios (see Figures 5-8). Among the seventeen data sets, only three (data sets 5, 6 and 12) produced rigorously the same MAP network for the two samplings. However, our Bayesian study presents interesting aspects. First, the expected closer relationships between wild subpopulations and cultivated subpopulations (Jap/Or3, Ind/Or1I, cAus/Or1A) appear in most MAP networks.

**Figure 6.**
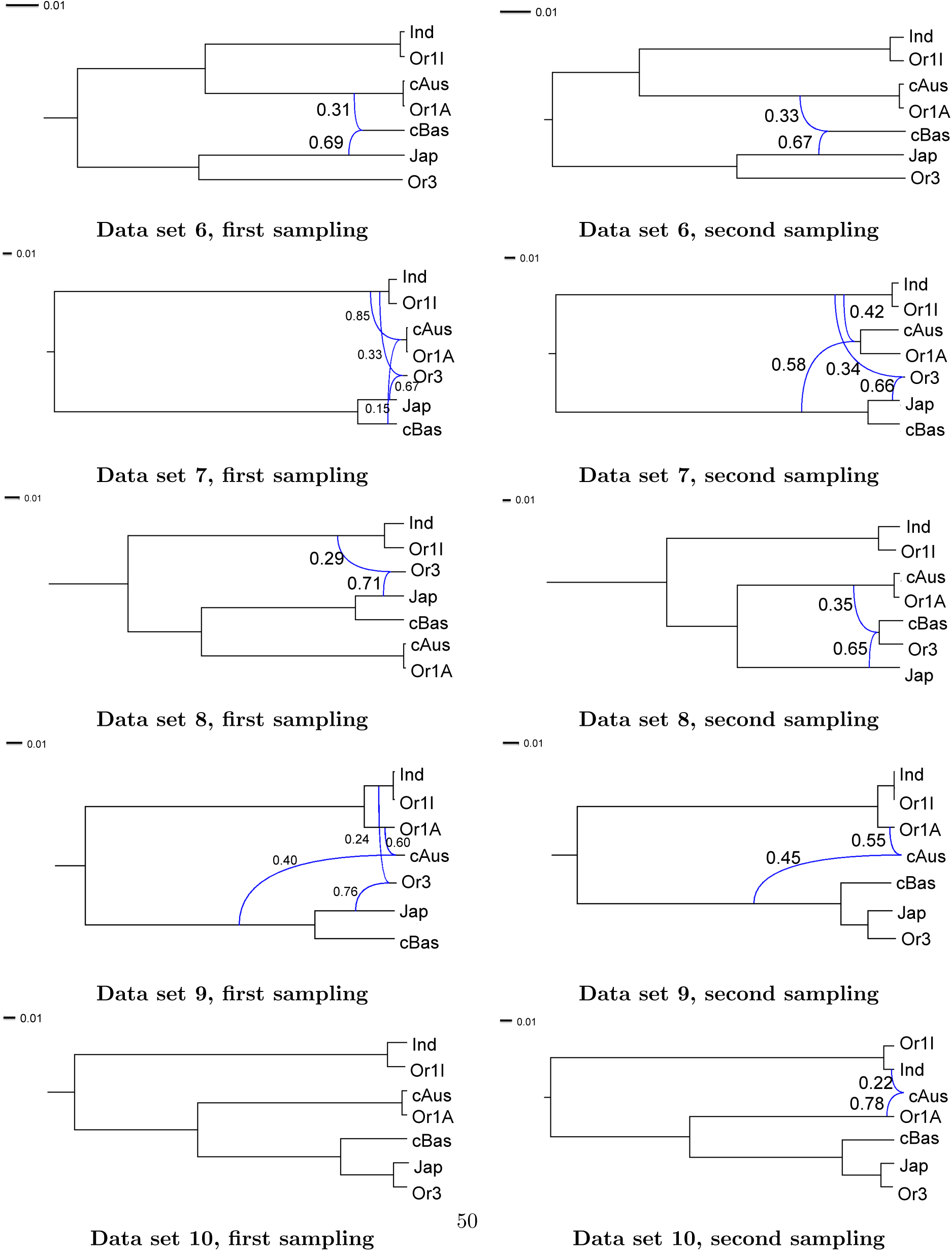
The MAP phylogenetic networks obtained for data sets 6-10 with one variety per subpopulation. The two different samplings of 12k markers along the rice genome are considered here. Inheritance probabilities are reported above reticulation edges and branch lengths are given in units of expected number of mutations per site (see the scale at the top left).

**Figure 7.**
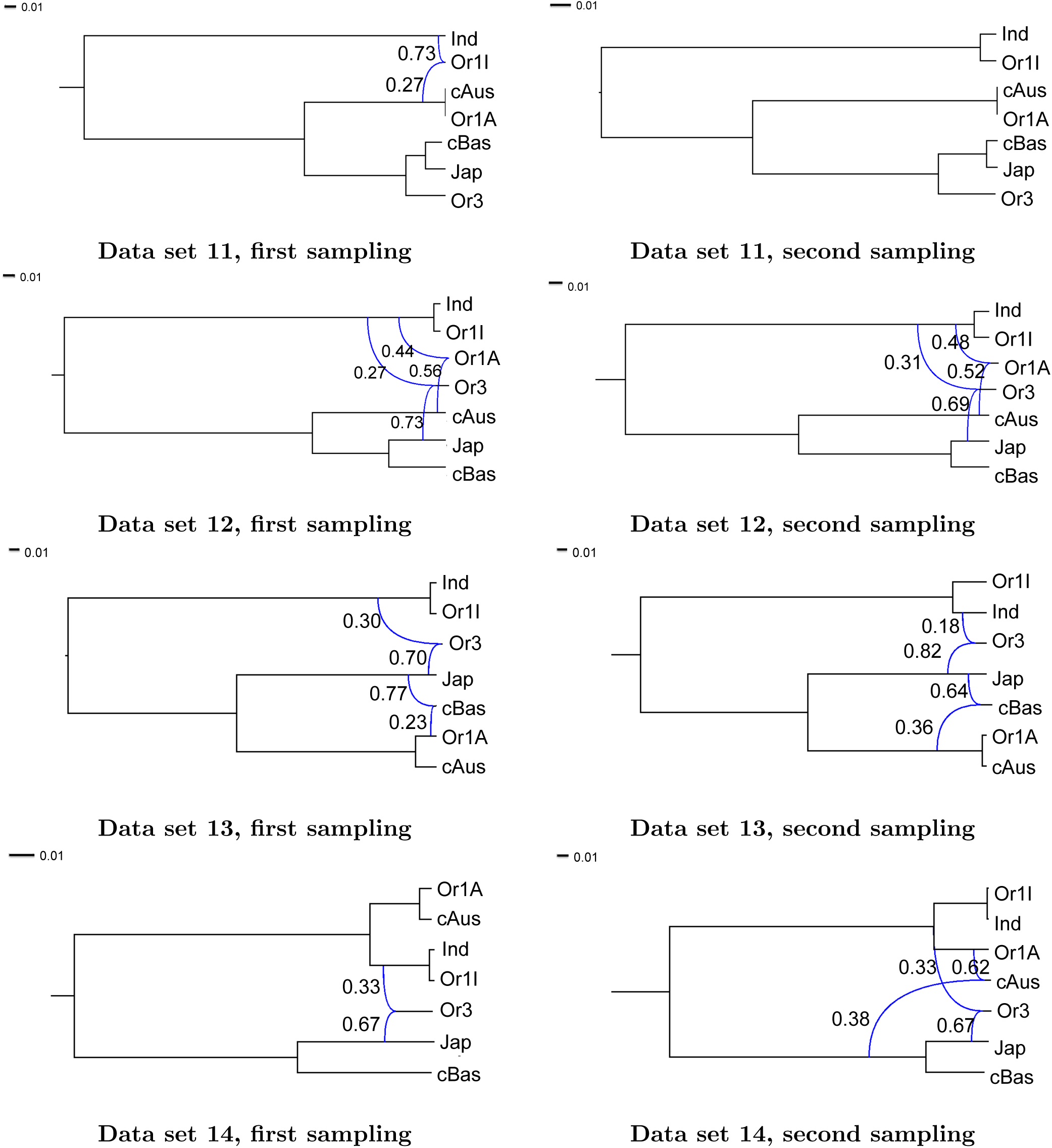
The MAP phylogenetic networks obtained for data sets 11-14 with one variety per subpopulation. The two different samplings of 12k markers along the rice genome are considered here. Inheritance probabilities are reported above reticulation edges and branch lengths are given in units of expected number of mutations per site (see the scale at the top left).

**Figure 8.**
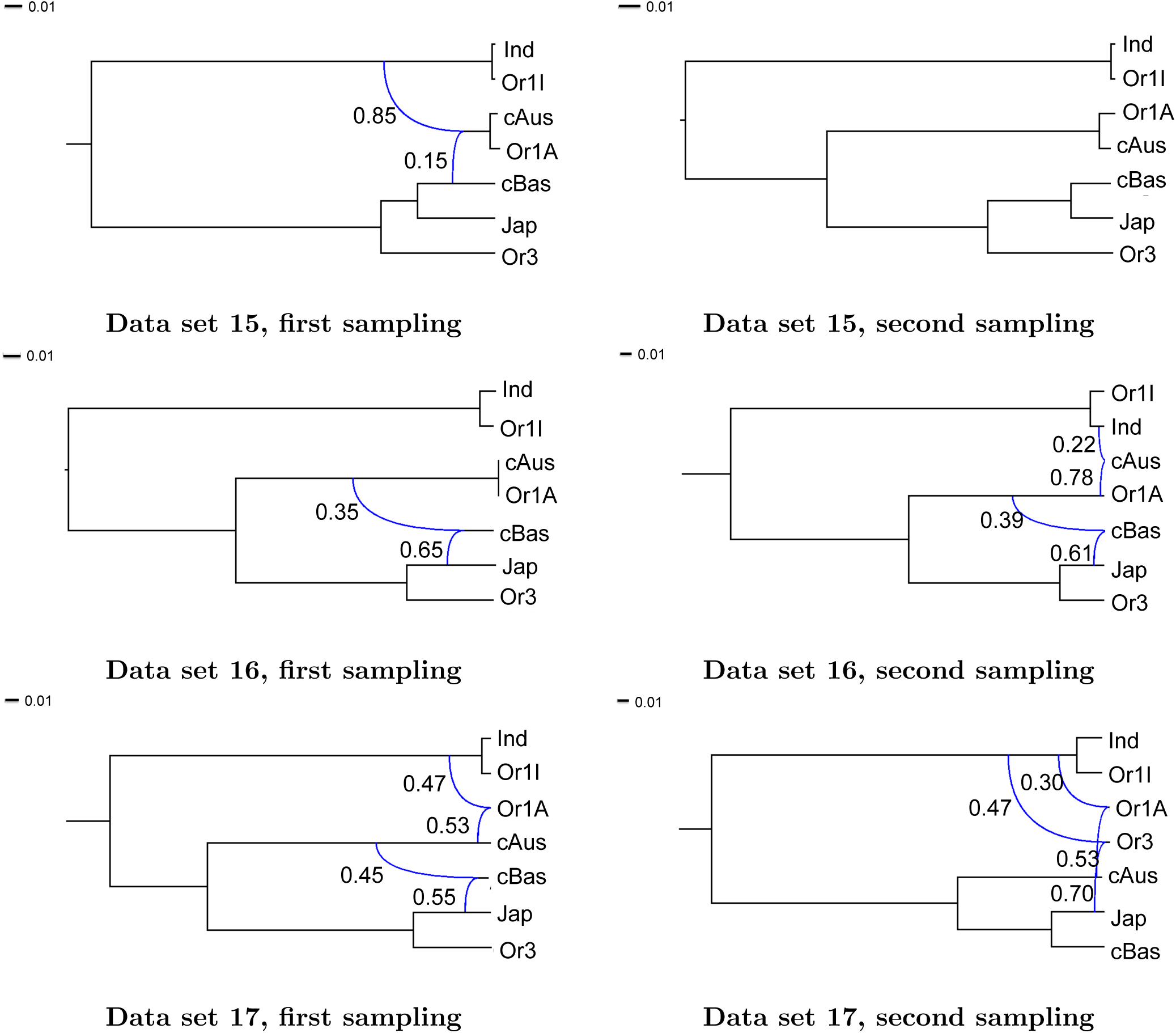
The MAP phylogenetic networks obtained for data sets 15-17 with one variety per subpopulation. The two different samplings of 12k markers along the rice genome are considered here. Inheritance probabilities are reported above reticulation edges and branch lengths are given in units of expected number of mutations per site (see the scale at the top left).

**Figure 9.**
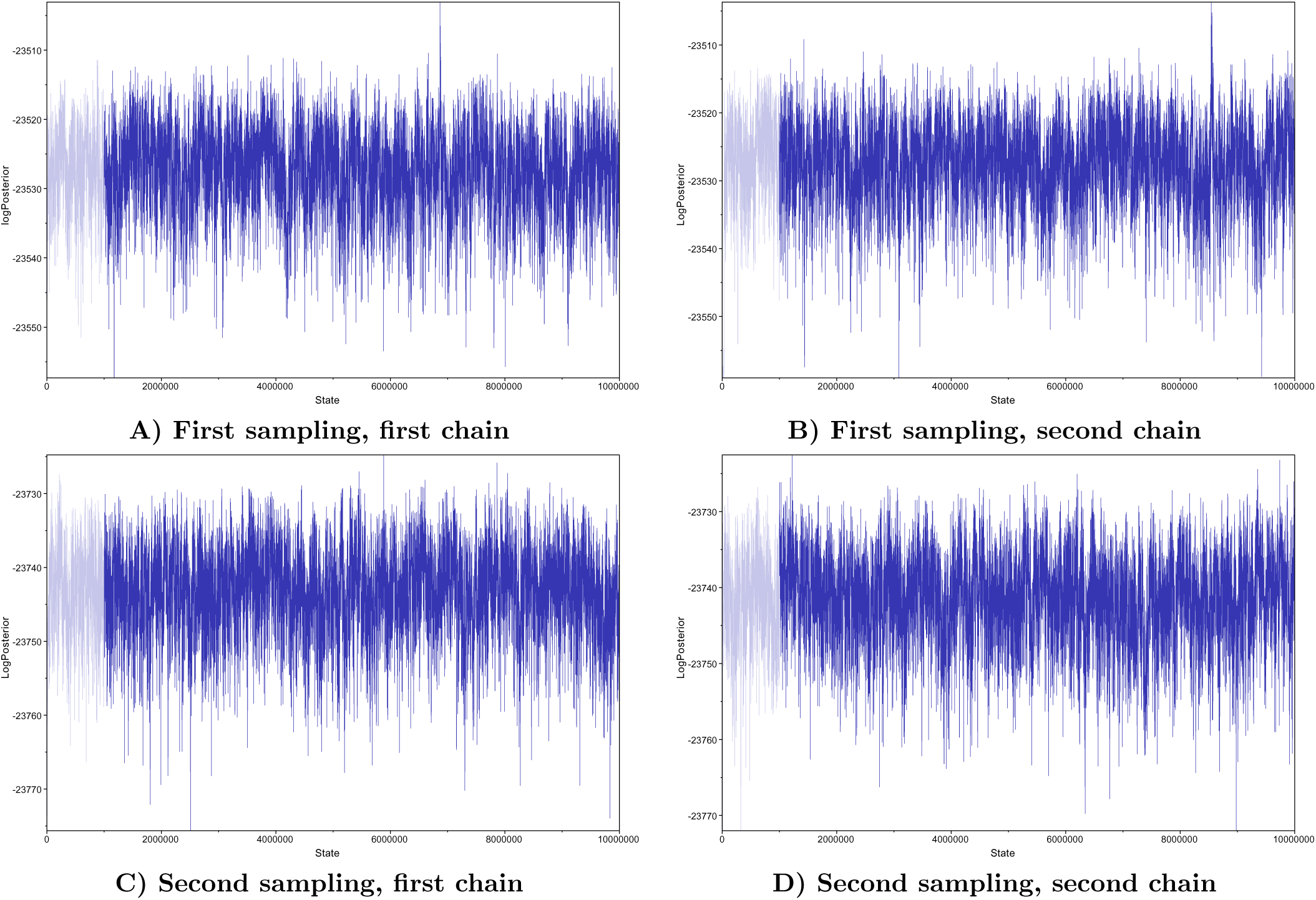
Trace plots obtained according to the Tracer software when data set 4 was analyzed with SnappNet. A) and B) refer to the first sampling of 12,000 SNPs along the whole genome, whereas C) and D) focus on the second sampling. Two chains were considered for each sampling.

These proximities are illustrated by such a couple of subpopulations either being descendant of a same speciation event or being descendant of a reticulate node (<u>i.e.</u>, involved in a hybridization).

A striking result is that many data sets (data set 4, 7 and 12) place Or3 as a hybrid derivative of Japonica x (Ind or Or1I). A predominant contribution of Japonica (close to 70%) is often highlighted. This is noteworthy and possibly representative of a phenomenon that complicates domestication analyses. Indeed, introgressions from cultivars to many wild rice populations have already been reported ([129]). It is a pattern that has been more often found in South and Southeast Asia than in China. Note that in data sets 1-17, the three Or3 representatives come from China.

A signal of hybridization was also found on cAus after analyzing data sets 5 and 9 (see Figures 5 and 6). According to the MAP phylogenetic networks, cAus is either a hybrid Jap x Or1A (data set 5), or a hybrid (Jap,cBas,Or3) x Or1A (data set 9). Although these two data sets incorporate diverse genetic materials, there is an agreement for a larger contribution from Or1A to cAus in the hybridization, as expected.

Data sets 1, 6, 13 and 16 (see Figures 5-8) clearly support the fact that cBas is the consequence of a reticulation involving Jap and the precursor of Or1-A/cAus. This hybridization on cBas is strongly supported by data set 16. The contributions are approximately 63% for Jap and 37% for Or1-A/cAus (see Figure 8). The posterior distribution sampled by SnappNet was concentrated on this scenario in 95.72% (resp. 98.15%) of sampled MCMC observations for the first (resp. second) marker sampling. The same hybridization was found from data set 6, with similar contributions to those obtained from data set 16. Last, data sets 1 and 13 both hesitate between an hybridization Jap x Or1A (cf. first sampling) or Jap x (Or1A x cAus) (cf. second sampling).

Overall, even though consistent signals emerge, our Bayesian analyses with a free exploration of the topology space led to a large spectrum of evolutionary scenarios, depending on the variety choice for each subpopulation. We showed the importance of genetic exchanges since most of the inferred topologies are networks and not trees.

### (2) Study of 16 evolutionary scenarios with SnappNet

To obtain a pertinent global picture, we increased the number of varieties per sub group, by adding extra varieties to the 29 considered in Step 1. We focused exclusively on the 16 network topologies described in Figure 3. Most of these networks present features found in our Bayesian analysis (Step 1), such as the association between wild subpopulations and cultivated subpopulations, the hybridizations on cAus and on cBas. We discarded the hybridization on Or3 since such pattern is peculiar and cannot constitute the backbone of a domestication scenario, supposed to mainly depict how cultivars derived from wild forms. We refer to Section 2.4 for more details on the chosen networks.

With the help of SnappNet, we computed the likelihoods of the 16 networks on data sets 18-21, and we penalized more complex models with the AIC [1] criterion, as in [53].

#### Scanwidth and computational burden

Table 2 reports the number of iterations performed by SnappNet after six months of computation for each studied network. The table also reports the <u>scanwidth</u> associated to each network. Recall that SnappNet provided a break-through in computation speed thanks to a new way to compute the network likelihoods, based on this scanwidth parameter [101, 60]. In a nutshell, the scanwidth measures the tree-likeness of a network’s topology. To get an intuition, imagine scanner lines rising up the network from the leaves up to the root. Initially, each branch above a leaf has its own horizontal scanner line. This set of lines evolves in steps, with each step involving bypassing an internal network node located directly above a branch traversed by a scanner line. Whenever this internal node is a speciation node, the two lines crossing branches below that node are merged into a single line. Whenever, a reticulation node is bypassed, the scanner line crossing the branch below the node is raised locally to cross the two branches above the node instead of the one below. Thus, bypassing an internal node can result either in reducing by one the number of scanner lines or by increasing by one the number of branches crossed by a scanner line. This number of crossed branched is called the <u>width</u> of the line. In contrast to species trees where likelihood can be computed for each branch independently, network likelihoods considered by SnappNet requires some branches to be considered simultaneously in order to compute joint probability distributions. More precisely, SnappNet has to jointly consider all branches crossed by a same scanner line. Hence, to reduce the computational burden as much as possible, one aims at making the scanner lines cross the smallest possible number of branches. The <u>scanwidth of a total order on the internal nodes</u> of the network is defined as the maximum width of a scanner line it induces when traversing the network from the leaves up to the root while bypassing internal nodes in this order. The <u>scanwidth of the network</u> is the smallest scanwidth over all possible orders on its internal nodes. Note that finding an order exactly minimizing this parameter is an NP-hard problem [13] but SnappNet includes a heuristic method that allows for the approximate minimization of this parameter.

According to Table 2, as might be expected, within the allotted time, networks with a scanwidth of 2, performed much more iterations than those with a scanwidth of 3. For instance, the minimum number of iterations was equal to 397,061, resp. 14,651 for networks of scanwidth 2, resp. 3. In the same way, the maximum number of iterations was found to be equal to 3,338,825 and 303,303 for networks of scanwidth 2 and 3, respectively.

#### Topologies of the most likely scenarios

Tables 3-6 that focus on data sets 18-21, report the log likelihoods and the AIC values associated to the different studied networks. For each criterion, the ranking between networks is presented in brackets. Note that Tables 3 and 4 (resp. Tables 5 and 6) differ only in the set of markers sampled along the genome.

Moreover, Table 7 presents a summary on analyses on data sets 18-21. Figure 10, complementary to Table 7, reports inheritance probabilities estimated by SnappNet for the most likely evolutionary scenarios (see Figures 19-20 in Supplementary Material for the details).

**Figure 10.**
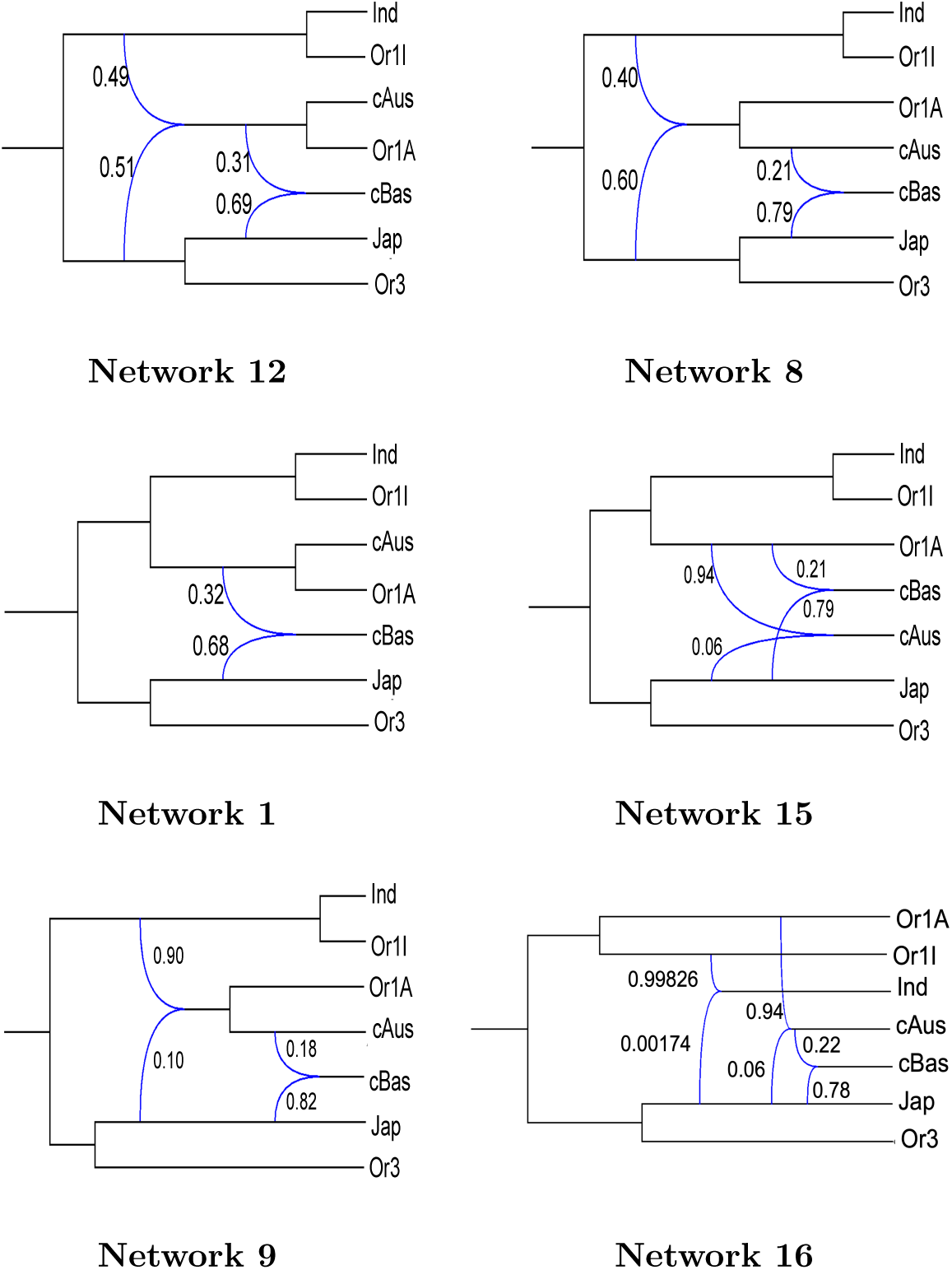
Inheritance probabilities estimated by SnappNet for the most likely evolutionary scenarios. Reported inheritance probabilities are averages on data sets 18, 19, 20 and 21, and on the two samplings.

According to Table 7, Networks 12, 8, 1, 15, 9 and 16 are the most likely scenarios. They a) place the split between the Indica ancestor and Japonica ancestor closest to the root (cf. Figure 10 below and also Figures 19-20 in Supplementary Material) and b) place the origin of the cBasmati lineage in close connection to both Japonica and cAus, either directly (Networks 8, 9, 16) or through related immediate ancestry (Networks 12, 1 and 15). As expected, the most likely networks feature the association between wild and cultivated subpopulations (Jap/Or3, Ind/Or1I, cAus/Or1A).

The least likely scenarios are Networks 6, 7, 14 and 2. Network 6 does not present the associations Ind/Or1I and cAus/Or1A, whereas Network 7 places cAus closer to Japonica than to Indica, which is unusual. In view of our numerical results for Network 14, Or1A contributed to cAus for approximately 95%. Consequently, due to Network 14’s topology, this network considers that the (cAus/Or1A) clade and the (Ind,Or1I) clade are far appart. Last, Network 2 depicts a singular scenario with the emergence of Indica and of a novel wild form at the origin of cAus and Or1A.

On the other hand, Networks 10, 13, 4, 3, 5 and 11 that have been attributed intermediate rankings, can be reasonably discarded for the rest of our study. First, Networks 10, 13, 3 and 5 exhibit a suspicious recent domestication of Japonica that happened after the hybridization at the origin of cBas: this thesis appears highly unrealistic in view of the rice literature. Note also that Network 5 considers that cAus is closer to Japonica than to Indica. Furthermore, Network 4 is <u>displayed</u> by Network 8 but in contrast to the latter, cAus is not connected to Indica anymore, which seems to be unlikely. The missing reticulation edge in Network 4 presented an important weight (0.40) in Network 8 (see Figure 10). Last, according to Network 11, the wild Or1A have received genetic material from Japonica, which seems implausible.

Consequently, it seems reasonable to concentrate exclusively on networks 12, 8, 1, 15, 9 and 16 for a deeper analysis based on a larger data set. Note that beyond these biological considerations, Table 7 shows that there is an important gap in summed ranking between these networks and the following ones.

Before studying in more detail the six most likely networks, let us give precisions on continuous parameters (e.g. population sizes, branch lengths, inheritance probabilities) estimated by SnappNet for the most likely scenarios.

#### Population sizes for the most likely scenarios

Figure 11 in this manuscript and Figures 21-25 in Supplementary Material give all the population sizes and branch lengths associated to these networks. Reported values are averages on data sets 18-21, and on the two samplings. Table 8 presents a summary of these figures by focusing exclusively on wild and cultivars subpopulations. We can notice that overall, population sizes were found larger for wild species than for cultivated rice. Recall that in SnappNet’s mathematical model, *θ* denotes the expected number of mutations between two individuals chosen randomly. In other words, as expected, wild species exhibited more diversity than cultivars. For instance, population sizes for the clade (Or3, Jap) were estimated at (0.2224, 0.2077) for Network 1, at (0.1949, 0.1201) for Network 8 and at (0.1993, 0.1247) for Network 9. In the same way, population sizes associated to the split (Or1A, cAus), were found equal to (0.3302, 0.0314) for Network 12, to (0.1956, 0.1184) for Network 15, and to (0.1866, 0.1217) for Network 16. Surprisingly, population sizes for Or1I were found greater than those of Indica in only half of cases. A possible explanation for Network 16 lies in the fact that it depicts a scenario where Japonica and Or1I contributed together to Indica.

**Figure 11.**
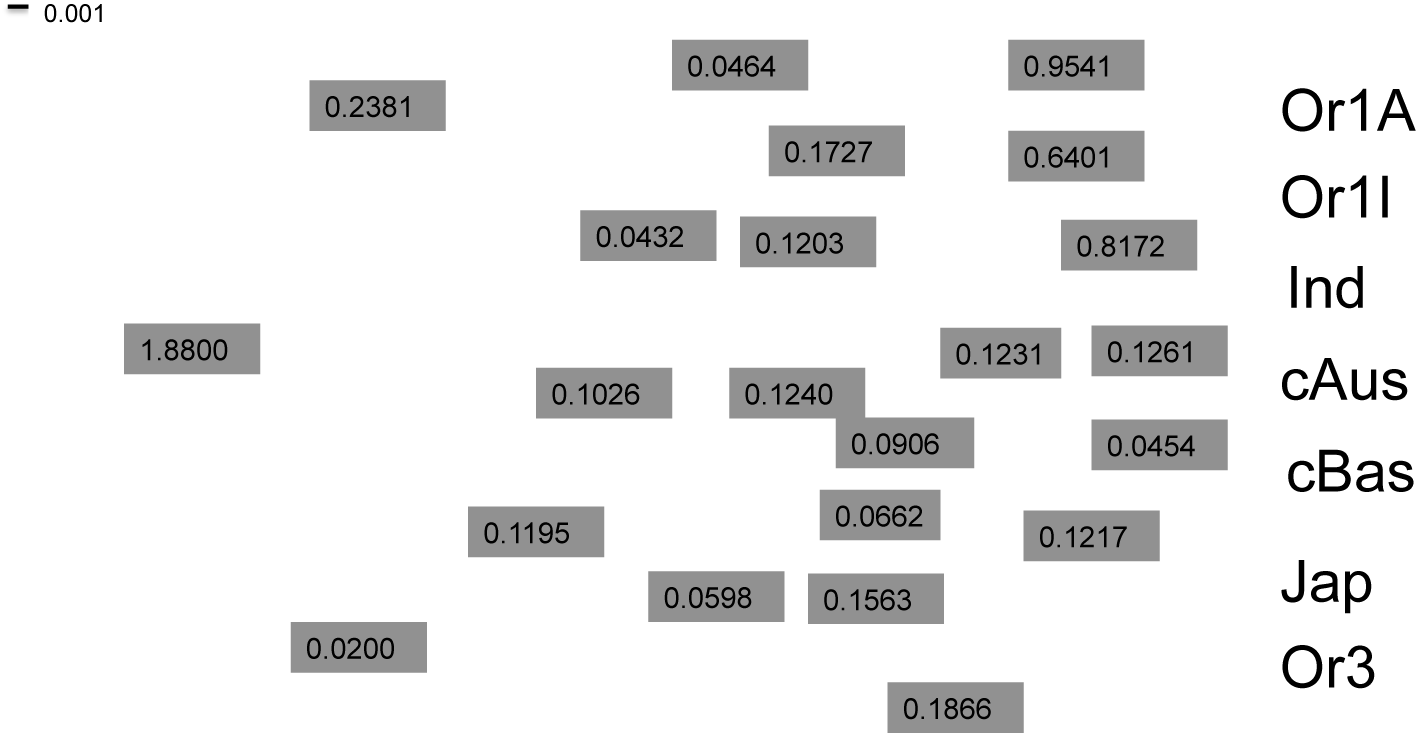
Population sizes and branch lengths estimated by SnappNet for **Network 16**. Population sizes are given in grey boxes, whereas branch lengths are given represented in units of expected number of mutations per site (see the scale at the top left). Reported values are averages on data sets 18, 19, 20 and 21, and on the two samplings.

**Table 8.** Population sizes estimated by SnappNet for the most likely scenarios. The focus is on wild and cultivars subpopulations.

|  | Or1A/cAus | Or3/Jap | Or1I/Ind |
| --- | --- | --- | --- |
| **Net 1** | 0.2227/0.0119 | 0.2224/0.2077 | 0.1161/0.1516 |
| **Net 8** | 0.1028/0.0876 | 0.1949/0.1201 | 0.4930/0.2740 |
| **Net 9** | 0.4428/0.0774 | 0.1993/0.1247 | 0.2661/0.2823 |
| **Net 12** | 0.3302/0.0314 | 0.2208/0.1519 | 0.2434/0.2146 |
| **Net 15** | 0.5032/0.0629 | 0.1956/0.1184 | 0.1420/0.0948 |
| **Net 16** | 0.9541/0.1261 | 0.1866/0.1217 | 0.6401/0.8172 |

#### Inheritance probabilities for the most likely scenarios

For each of the most likely scenarios, Figure 10 gives the average inheritance probabilities over the data sets 18-21 and on the two samplings. Moreover, the probabilities associated to each data set, are reported in Figures 19 and 20 in Supplementary Material. According to the different figures, we can observe that for Networks 1, 8, 9, 15, and 16, the inheritance probabilities were consistent over the different data sets. Surprisingly, for Network 12 that was ranked first, we found significant differences among data sets. The dosage ratios were estimated at 77:23 for data set 18 and at 16:84 for data set 19. Next, a smoothing effect was noticed for data sets 20 and 21. Since these data sets contain a mixture of varieties from data sets 18 and 19, more balanced contributions were obtained. In view of this instability on Network 12, it seems crucial to discriminate between networks thanks to a larger data set.

### (3) Discriminating between the **6** best scenarios with the Snarf hybrid approach

In order to investigate more in detail the six most likely evolutionary scenarios found previously, we present here the analysis of a larger data set thanks to the new hybrid approach, called Snarf, presented in this paper. In this context, Data set 22 includes now 64 varieties.

Recall that Snarf combines SnappNet’s accuracy with the potential of ABC-RF. In our case, for calibrating prior distributions, ABC-RF benefits from SnappNet’s parameter estimations obtained by penalized likelihood at the previous step. Note that branch lengths and instantaneous rates always relied on SnappNet’s estimates, even when our investigated priors were “free” from SnappNet. Indeed, previous studies (e.g. [115]) have shown that branch lengths play a key role in ABC approaches. In what follows, we first assess the performances of this hybrid approach on simulated data, and we concentrate later on real data.

#### Performances of the Snarf hybrid approach on simulated data

In order to make the reading easier, let us briefly discuss the impact of different factors on the performance of our hybrid approach. We refer to Section 5 in Supplementary Material for a full description of the results of this simulation study.

##### Impact of the inheritance prior

Incorporating a very specific prior for the inheritance probability helps to differentiate the evolutionary histories more easily. SnappNet was found rewarding since it allowed to calibrate a more performant prior.

##### Impact of the prior on the instantaneous rates

Choosing a prior on *u* and *v* or considering the MLEs, does not seem to have an impact on performances of the classifier.

##### Impact of the population size prior

Overall, the two investigated priors on population sizes gave similar performances.

##### Impact of the number of trees in the RF

A forest of 1,000 trees seems to be a good fit for all our experiments since the errors were stabilized.

##### The most important SS

It is clear that the Hils statistics [77], that refer to hybridization tests, are the most important contributors to the classifier. Note also that the D-statistic by [48, 96] and the estimators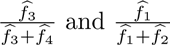 (see [15]) play an important role in our ABC-RF analysis.

##### Impact of the number of rows of the reference table

As expected, the prior error rate decreases when the number of rows of the reference table increases. Since the out of bag error rates (<u>i.e.</u> prior error rates) obtained for *N*_Ref_ =84,000 was already fully satisfactory, we did not explore larger reference tables.

##### Conclusion of the simulation study

Our method enjoyed very good performances on simulated data, whatever the prior under study. Indeed, on average, the prior error rate was found to be equal to 7.05% (cf. Section 5 in Supplementary Material).

#### Real data analysis with the Snarf hybrid approach

In this section, we show how Snarf, using real data, helps to identify the most plausible scenario for tracing the history of rice. We use the ABC-RF classifier once it has been trained on simulated data. Table 9 based on Data set 22, gives the percentage of trees supporting each evolutionary scenario, as a function of all the investigated prior distributions. As before, two samplings are under study. Table 9 also reports the posterior probability of selecting the true model (i.e. the true network), thanks to a supplementary RF. This new RF was trained on the same reference tables as before but differs from the previous RF since it uses the binary model prediction error as dependent variable (cf. [100] for more details).

**Table 9.**
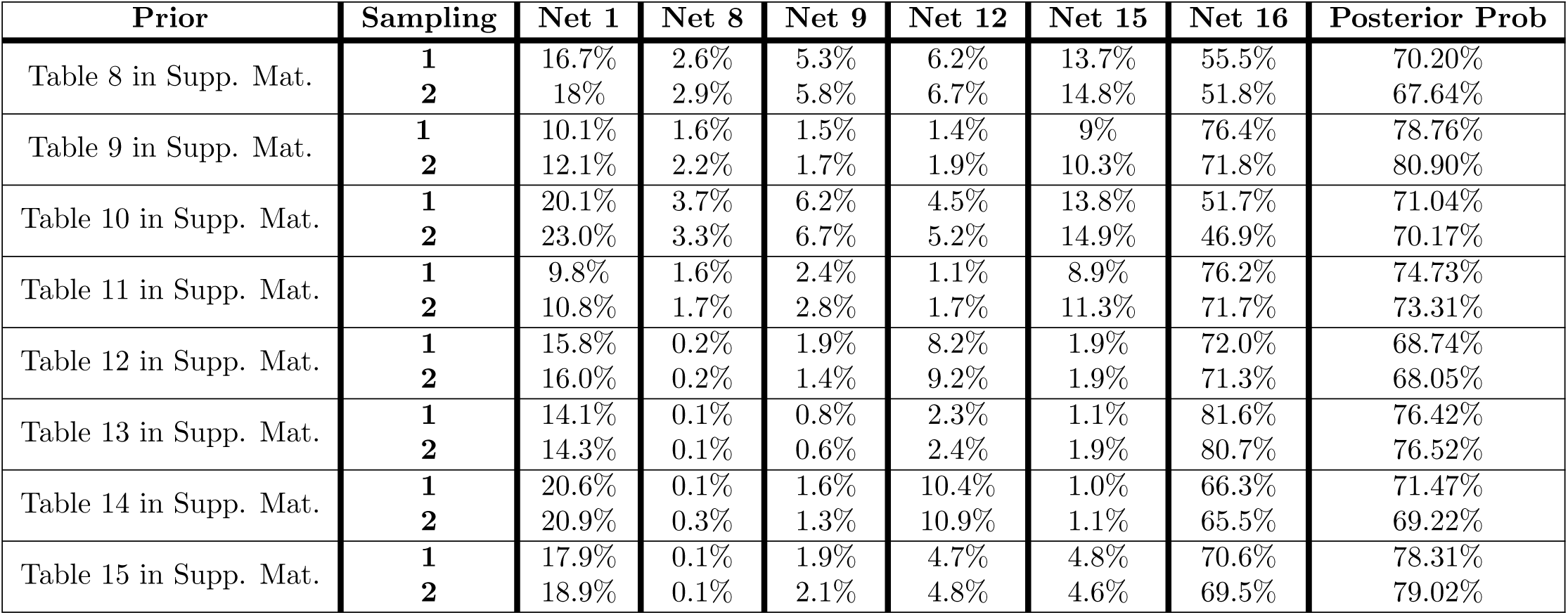
Percentage of the random forest trees supporting each evolutionary scenario as a function of the prior distributions. Values were obtained for two samplings of Data set 22 (real data). The posterior probability of selecting the true network is given in last column.

| Prior | Sampling | Net 1 | Net 8 | Net 9 | Net 12 | Net 15 | Net 16 | Posterior Prob |
| --- | --- | --- | --- | --- | --- | --- | --- | --- |
| Table 8 in Supp. Mat. | 1 | 16.7% | 2.6% | 5.3% | 6.2% | 13.7% | 55.5% | 70.20% |
|  | 2 | 18% | 2.9% | 5.8% | 6.7% | 14.8% | 51.8% | 67.64% |
| Table 9 in Supp. Mat. | 1 | 10.1% | 1.6% | 1.5% | 1.4% | 9% | 76.4% | 78.76% |
|  | 2 | 12.1% | 2.2% | 1.7% | 1.9% | 10.3% | 71.8% | 80.90% |
| Table 10 in Supp. Mat. | 1 | 20.1% | 3.7% | 6.2% | 4.5% | 13.8% | 51.7% | 71.04% |
|  | 2 | 23.0% | 3.3% | 6.7% | 5.2% | 14.9% | 46.9% | 70.17% |
| Table 11 in Supp. Mat. | 1 | 9.8% | 1.6% | 2.4% | 1.1% | 8.9% | 76.2% | 74.73% |
|  | 2 | 10.8% | 1.7% | 2.8% | 1.7% | 11.3% | 71.7% | 73.31% |
| Table 12 in Supp. Mat. | 1 | 15.8% | 0.2% | 1.9% | 8.2% | 1.9% | 72.0% | 68.74% |
|  | 2 | 16.0% | 0.2% | 1.4% | 9.2% | 1.9% | 71.3% | 68.05% |
| Table 13 in Supp. Mat. | 1 | 14.1% | 0.1% | 0.8% | 2.3% | 1.1% | 81.6% | 76.42% |
|  | 2 | 14.3% | 0.1% | 0.6% | 2.4% | 1.9% | 80.7% | 76.52% |
| Table 14 in Supp. Mat. | 1 | 20.6% | 0.1% | 1.6% | 10.4% | 1.0% | 66.3% | 71.47% |
|  | 2 | 20.9% | 0.3% | 1.3% | 10.9% | 1.1% | 65.5% | 69.22% |
| Table 15 in Supp. Mat. | 1 | 17.9% | 0.1% | 1.9% | 4.7% | 4.8% | 70.6% | 78.31% |
|  | 2 | 18.9% | 0.1% | 2.1% | 4.8% | 4.6% | 69.5% | 79.02% |

According to the table, Network 16 is clearly the model selected by ABC-RF, and this holds true regardless of the considered prior. Indeed, the votes for this scenario dominates from 70.20% to 78.76% for the first sampling, and from 67.64% to 80.90% for the second sampling. Moreover, the posterior probability of selecting the true model was relatively high. As a result, there is strong support for an evolutionary history with a single domestication, the one of Japonica. According to Network 16, the association of Japonica with the wild Or1A is at the origin of the cultivar cAus. In the same way, Japonica helped to convert the wild Or1I into Indica. In a nutshell, a precursor of Japonica is at the genesis of the Indica, cBas and cAus modern cultivars. This scenario argues for a single domestication, Japonica, followed by extensive exchanges at the origin of modern cultivars. This is not a surprise, since such a scenario is largely present in the literature (e.g. [61, 140, 27, 53, 138]). In contrast, other studies [56, 29, 130, 69] supported a multiple domestication scenario – as reported here by Networks 1, 15 and 12, 9 and 8, claiming that the domestication of Japonica and Indica occurred independently, in southern China and in India. Note that in our previous study [101], Network 1 was the final scenario suggested by SnappNet in a Bayesian setting. In view of our present more in-depth ABC-RF study (64 varieties vs. only 14 varieties in [101]), Network 1 is only the second preferred scenario. However, it is ranked far behind Network 16, since the votes for Network 1 only ranged from 9.8% to 20.6% for the first sampling, and from 10.8% to 20.9% for the second sampling (see Table 9).

As a consequence, according to the present study, there is a strong evidence that Network 16 depicts the reticulated history of rice domestication. In particular, according to this penalized likelihood study, the oldest reticulations match introgressions from Japonica into Indica and from Japonica into cAus. The inferred contributions are very imbalanced: Japonica is involved in only 0.174% of the Indica genome and in only 6% of the cAus genome. Coming back to the recent hybridization at the origin of the cBasmati lineage, Network 16 features cBasmati as a hybrid derivative of the Japonica and cAus lineages. According to Section 3, the dosage ratio (Japonica vs. cAus) is found to be equal to 78:22 on average (see Figure 10), reflecting as expected a more important contribution of Japonica. Since the contributions are more balanced, cBas can be viewed as the result of an admixture event between Japonica and cAus.

Last, the branch length associated to cBas was found very close to the one leading to cAus. This roughly matches 80% of Indica’s branch length (see Figure 11). Consequently, according to the present study, cBas seems to be a relatively old population. The population sizes were found approximately 3 times larger for Japonica than for cBas (cf. 0.1217 vs 0.0454 in Figure 11), which is not surprising since Japonica presents three subgroups (tropical, subtropical, and temperate).

## Discussion

### Uncertainty regarding the evolutionary history of rice

In this paper, we investigated the rice domestication process in Asia. Rice is a remarkable species due to the existence of reticulate histories involving several episodes of domestication. There are several well established cultivar groups but debate is still unsettled regarding the number of domestications that occurred. Previous studies suggest either a unique domestication ([61, 27, 53, 138, 51]) or multiple domestications [29, 130, 146, 69]. Due to the ancient dissemination of the crop throughout the continent, there is a huge diversity in terms of time, geography and involved (sub)populations. As a result, this domestication process is very hard to infer. The present work offers more insights into rice evolution by proposing a new methodology based on a rich stochastic model, which incorporates introgression events, incomplete lineage sorting and mutations that occur over time.

We introduced a new hybrid approach, Snarf, that is built on SnappNet [101] and on ABC-RF [100]. In particular, our method exploits new features from SnappNet, like penalized likelihood which was not present in [101]. Furthermore, our study is innovative in that it tackles the computational burden of inference using an ABC-RF approach combined with network-specific summary statistics.

### Studies should not be limited to a single variety per subgroup

We started by freely exploring the network topology space, thanks to SnappNet in a Bayesian setting. During this preliminary step, we considered data sets with only a few varieties, <u>i.e.</u> with a single variety per subpopulation. As in our previous study [101], we classified varieties into groups or subpopulations. The wild rice *O.rufipogon* was distributed in three groups, Or1I, Or1A and Or3, reflecting a preferential proximity to Indica, cAus, and Japonica, respectively ([61]). On the basis of these small data sets, we explored freely the whole space of evolutionary scenarios, thanks to SnappNet in a Bayesian framework. This led to a large spectrum of evolutionary histories. For instance, Or3 was found as a hybrid derivative of Japonica x (Ind, Or1I) in a few of our experiments. Since wild forms are allogamous, it is not surprising to obtain scenarios underlying introgression from cultivars into wild forms. This is a pattern that has been more often found in South and Southeast Asia than in China. We also found a more expected signal of hybridization on cBas, and a signal of hybridization on cAus. Note that in the literature, several other studies relied on small data sets. For instance, [28] inferred evolutionary histories thanks to a three population topology involving trios with one representative for each population (cBas, Jap and *O.rufipogon*). According to the authors of this phylogenomic study, cBas and cAus are sister clades to Japonica and Indica, respectively. A signal of gene flow between cAus and cBas was also reported. In our case, we also observed such signal with a contribution of the precursor of Or1A/cAus for 37% to cBas.

Finally, we came to the conclusion that the complexity of rice evolution demanded to analyze a large amount of genetic material to obtain a pertinent global picture. Up to now, most network reconstruction methods, based on an explicit model of evolution, can only handle data sets containing a single individual per species when considering large numbers of SNPs. According to our preliminary study, this sample size is too small to have reliable results on rice evolution.

### A new hybrid approach shows a clear preference for one particular evolutionary scenario

For this purpose, we devised a new hybrid approach that leverages the power of classifiers based on summary statistics to handle much larger data sets. We used this new approach, called Snarf, to distinguish among 16 rice evolutionary scenarios that emerged from the preliminary study. These networks reflect features observed in our Bayesian analysis (step one) and most of them conform with the main features of the genetic structure. Overall, our statistical study revealed that a particular network (<u>i.e.</u>, Network 16) was by far the preferred scenario. It was heavily supported by ABC-RF for different samplings of real data and whatever the considered prior. Note that for ABC-RF, specific SS were implemented within SimSnappNet.

In the preferred scheme, Japonica contributed to other cultivar groups, cAus, cBas and Indica at different steps of the evolution. Japonica was the first to be domesticated, and then it was mobilized for domesticating Indica. Later, it also played a role in the domestication of cAus, and in the birth of cBas. In other words, our study supports the thesis of a single domestication, Japonica. Thus, there is an agreement with a large panel of papers ([61, 140, 27, 53, 138]) arguing that rice underwent a unique domestication in the Yangtze valley. Note that the recent study of [51], based on a simpler model, suggests a scenario close to ours (cf. their Figure 4e).

In addition, our hybrid approach also gives access to the inheritance probabilities, thanks to SnappNet in a penalized likelihood framework. We did not estimate continuous parameters by simulation based approaches (e.g. [107, 110]): estimates obtained by SnappNet must be more accurate since the likelihood calculation relies on the evolutionary model. In view of our stable results across different data sets for Network 16, cBas seems to result of an admixture between Japonica and cAus, with a dosage ratio 78:22 (see Figure 10). We also found a small contribution of Japonica to the emergence of Indica. Given the size of this contribution (0.174%), this can be considered as an introgression (see Figure 10). In the same way, introgressions from Japonica into cAus were detected, at a level of approximately of 6%. The domestication of Japonica has fixed a few alleles that are deleterious for wild populations but favorable for Indica and cAus. As a consequence, Japonica contributed by giving domestication alleles to Indica and to cAus. Then, in light of Table 1 of [113], it seems reasonable that 0.174% (resp. 6%) of the Indica (resp. cAus) genome comes from Japonica. Indeed, the number of domestication loci is usually very low since a too high level of selection can lead to population extinction (see [3] or [2] for a review). To sum up, in our study, we retrieved a few domestication alleles from Japonica in Indica and cAus genomes, on the basis of carefully chosen varieties. This pattern could be seen even more clearly, in future research with a larger sample size. Note that the reticulations events on cAus and on cBas were already observed when we concentrated on small data sets with SnappNet in a Bayesian setting.

Another interesting aspect of Snarf lies in the fact that it gives estimates for branch lengths and population sizes. Then, we have highlighted that cBas was an old group that was not derived from a recent hybridization involving modern Japonica. As a consequence, we observe differences with the recent study of [51] on the origin of cBas. According to our study, the Japonica component of cBas is certainly ancient. In the same way, the cAus component is ancient, and we estimated its origin just after the emergence of cAus, whose origin is an introgression from Japonica.

### The limitations of this study

Our work relies on the NMSC that assumes independent sites. As a result, we considered different marker samplings along the rice genome and checked the robustness of our methods. In the future, it would be very interesting to be able to exploit complete genomes instead of sampling along the genome. In that sense, we would model linkage disequilibrium between sites. It would require to study networks evolving within a phylogenetic network, as suggested in the review by [32]. In this context, a natural stochastic process is the coalescent with recombination [63, 135], or its approximation [88], and theoretical results are given in [120, 59]. Furthermore, ancestral recombination graphs [36, 52] and their approximations [86, 39, 134, 105, 71, 83] are highly linked to this topic (see [95, 79] for a review). Another possible extension of our present work is to consider the model of [42], that generalizes NMSC, by incorporating variations among inheritance probabilities along the genome. This way, it takes into account the fact that selection occurs differently between loci.

A question raised by our study is related to the number of varieties to consider in ABC-RF. Recall that here, we considered a total of 64 varieties, and a maximum of 10 varieties per subpopulation.

For computing SS 6-18 and SS 19-22 based on [77, 97], the current version of SimSnappNet lets the user choose to decide whether considering all varieties or only drawing a few varieties at random. Recall that SS 6-18 and SS 19-22 are based on sets of four and five species, respectively. In our manuscript, in order to limit the computational burden, we considered respectively a total of 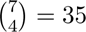 and 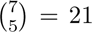 sets of four and five species. Indeed, once the four (resp. five) species were chosen among the 7 species, we considered a unique assignment to species *O*, *P*_1_, *H*, *P*_2_ (resp. *O*, *P*_1_, *P*_2_, *P*_3_ and *P*_4_) of the network (resp. species tree) in Figure 4. If the total number of subpopulations had been small, we could have imagined taking into account all possible assignments. Furthermore, for each set of four (resp. five) species, we focused on 5^4^ = 625 (resp. 5^5^ = 3125) random sets of 4 (resp. 5) varieties. To sum up, the ratio between our sampling and all possibilities was found to be equal to 3.01% for SS 6-18 and to 1.66% for SS 19-22. Overall, we handled 562 SS, and this setting gave satisfactory results for our classifier. Future users should figure out the balance between the number of varieties to draw and the number of sets of four and five species to handle.

Last, the choice of the SS is not exhaustive and new SS could be implemented within SimSnappNet in years to come. According to our ABC-RF analysis, the Hils statistic (see [77]) for testing hybridization was found as the most important contributor to the classifier. This test relying on 4 species was more relevant than the DFOIL one [97] for the presence of introgressions in a symmetric five-taxon phylogeny.

### Future work

In this paper, we focused on the rice domestication process in Asia but the Snarf hybrid approach is general and could be applied to other kinds of data (human, animals, other plants…). In the future, we could envision two other hybrid approaches, cousins of Snarf. The first one would combine SnappNet and deep learning, whereas the second one would combine a Pseudo Likelihood (PL) method and abcrf with the help of SimSnappNet and its new SS dedicated to phylogenetic networks.

## Acknowledgments

The authors thank Celine Scornavacca for having suggested a few summary statistics based on phylogenetic networks. We are grateful to the genotoul bioinformatics platform Toulouse Occitanie (Bioinfo Genotoul, https://doi.org/10.15454/1.5572369328961167E12) for providing computing resources.

## Data Availability

All data files are available from the github public repository located at https://github.com/rabier/SNARF. Note that, imputed data of wild forms were downloaded from the link located at the bottom of the following webpage http://viewer.shigen.info/oryzagenome2detail/downloads/index.xhtml

