## Supplementary Material for "A hybrid approach combining a phylogenetic method and Approximate Bayesian Computation Random Forest for phylogenetic network inference: application to the rice domestication process in Asia"

August 5, 2026

#### Contents


**1 Supplementary informations on rice real data** **2**

**2 Free exploration of the topology space with SnappNet** **9**

**3 Study of 16 evolutionary scenarios with SnappNet** **22**

**4 Extra informations on SS6-18 and SS19-22** **28**

4.1 Computing SS6-18 . . . . . . . . . . . . . . . . . . . . . . . . . . . 28

4.2 Computing SS19-22 . . . . . . . . . . . . . . . . . . . . . . . . . . . 28

**5 Discriminating between the 6 best scenarios with the Snarf hybrid method** **29**

### 1 Supplementary informations on rice real data

**Table 1. Description of the 29 varieties studied. These varieties are either representative cultivars spanning the four main rice subpopulations (Indica, Japonica, circum Aus and circum Basmati), or representative of the wild subpopulations (Or1I, Or1A, Or3).**

| Subpopulations | Variety ID | Country | Variety name |
|---|---|---|---|
| *circum Aus* | IRIS-313-11058 | Bangladesh | AUS 329 |
| | IRIS 313-11737 | India | CHUNDI |
| | IRIS-313-10852 | India | ARC 7336 |
| | IRIS-313-11027 | Pakistan | JHONA 101 |
| *circum Basmati* | IRIS-313-11062 | Bangladesh | BEGUNBICHI 33 |
| | IRIS-313-11825 | India | HANSRAJ |
| | IRIS-313-8326 | India | JC1 |
| | IRIS-313-11258 | India | ARC 13502 |
| | IRIS-313-10851 | India | ARC 7296 |
| | IRIS-313-12094 | Bangladesh | ARC KASHA |
| *Indica* | IRIS-313-11819 | Myanmar | PADINTHUMA |
| | IRIS-313-11089 | Cambodia | SRAU THMOR |
| | IRIS-313-11796 | China | DU GEN CHUAN |
| | IS-313-11646 | India | NCS771 A |
| | CX270 | Taiwan | TAICHUNGNATIVE1 |
| | IRIS-313-11741 | SriLanka | HERATH BANDA |
| *Japonica* | B204 | China | LONGHUAMAOHU |
| | IRIS-313-11924 | Thailand | NAM JAM |
| | IRIS-313-10577 | Philippines | IFUGAO RICE |
| | IRIS-313-11691 | Bhutan | SHANGYIPA |
| | IRIS-313-7883 | Indonesia | GANIGI |
| | B269 | China | YUEFU |
| *Or1I* | W1117 | India | W1117 |
| | W1559 | Thailand | W1559 |
| *Or1A* | W0574 | Malaya | W0574 |
| | W1747 | India | W1747 |
| *Or3* | W3042 | China | W3042 |
| | W3048 | China | W3048 |
| | W3073 | China | W3073 |

**Table 2. Data sets 1-17 that contain one variety per subpopulation. Varieties were chosen from Table 1 in order to build each data set.**

| Subpopulations | Dataset | Variety ID | Country | Variety name |
|---|---|---|---|---|
| *circum Aus* | 1,3,8,12,13,17 | IRIS-313-11058 | Bangladesh | AUS 329 |
| | 2,7,15 | IRIS 313-11737 | India | CHUNDI |
| | 4,6,9,11 | IRIS-313-10852 | India | ARC 7336 |
| | 5,10,14,16 | IRIS-313-11027 | Pakistan | JHONA 101 |
| *circum Basmati* | 1,13 | IRIS-313-11062 | Bangladesh | BEGUNBICHI 33 |
| | 2,12,17 | IRIS-313-11825 | India | HANSRAJ |
| | 3,7,11,15 | IRIS-313-8326 | India | JC1 |
| | 4,8,10,16 | IRIS-313-11258 | India | ARC 13502 |
| | 5,9,14 | IRIS-313-10851 | India | ARC 7296 |
| | 6 | IRIS-313-12094 | Bangladesh | ARC KASHA |
| *Indica* | 1,12,13,17 | IRIS-313-11819 | Myanmar | PADINTHUMA |
| | 2,11 | IRIS-313-11089 | Cambodia | SRAU THMOR |
| | 3,10,16 | IRIS-313-11796 | China | DU GEN CHUAN |
| | 4,7,15 | IS-313-11646 | India | NCS771 A |
| | 5,8,14 | CX270 | Taiwan | TAICHUNGNATIVE1 |
| | 6,9 | IRIS-313-11741 | SriLanka | HERATH BANDA |
| *Japonica* | 1,7,14 | B204 | China | LONGHUAMAOHU |
| | 2,8 | IRIS-313-11924 | Thailand | NAM JAM |
| | 3,9 | IRIS-313-10577 | Philippines | IFUGAO RICE |
| | 4,12,13,16 | IRIS-313-11691 | Bhutan | SHANGYIPA |
| | 5,11,15 | IRIS-313-7883 | Indonesia | GANIGI |
| | 6,10,17 | B269 | China | YUEFU |
| *Or1I* | 1,4,8,10,12,13,16,17 | W1117 | India | W1117 |
| | 2,3,5,6,7,9,11,14,15 | W1559 | Thailand | W1559 |
| *Or1A* | 1,3,4,6,7,8,10,11,13,15,16 | W0574 | Malaya | W0574 |
| | 2,5,9,12,14,17 | W1747 | India | W1747 |
| *Or3* | 1,4,9,11,13 | W3042 | China | W3042 |
| | 2,5,7,10,12,14,15,16,17 | W3048 | China | W3048 |
| | 3,6,8 | W3073 | China | W3073 |

**Table 3. Data sets 18 and 19 that contain a few varieties per subpopulation. Varieties were chosen from Table 1 in order to build each data set. These data sets are used in the analysis of the 16 phylogenetic networks.**

| Subpopulations | Dataset | Variety ID | Country | Variety name |
|---|---|---|---|---|
| *circum Aus* | 18 | IRIS-313-11058<br>IRIS 313-11737 | Bangladesh<br>India | AUS 329<br>CHUNDI |
| | 19 | IRIS-313-10852<br>IRIS-313-11027 | India<br>Pakistan | ARC 7336<br>JHONA 101 |
| *circum Basmati* | 18 | IRIS-313-11062<br>IRIS-313-11825<br>IRIS-313-8326 | Bangladesh<br>India<br>India | BEGUNBICHI 33<br>HANSRAJ<br>JC1 |
| | 19 | IRIS-313-11258<br>IRIS-313-10851<br>IRIS-313-12094 | India<br>India<br>Bangladesh | ARC 13502<br>ARC 7296<br>ARC KASHA |
| *Indica* | 18 | IRIS-313-11819<br>IRIS-313-11796<br>IRIS-313-11089 | Myanmar<br>China<br>Cambodia | PADINTHUMA<br>DU GEN CHUAN<br>SRAU THMOR |
| | 19 | IS-313-11646<br>CX270<br>IRIS-313-11741 | India<br>Taiwan<br>SriLanka | NCS771 A<br>TAICHUNGNATIVE1<br>HERATH BANDA |
| *Japonica* | 18 | B204<br>IRIS-313-11924<br>IRIS-313-10577 | China<br>Thailand<br>Philippines | LONGHUAMAOHU<br>NAM JAM<br>IFUGAO RICE |
| | 19 | IRIS-313-11691<br>IRIS-313-7883<br>B269 | Bhutan<br>Indonesia<br>China | SHANGYIPA<br>GANIGI<br>YUEFU |
| *Or1I* | 18,19 | W1117<br>W1559 | India<br>Thailand | W1117<br>W1559 |
| *Or1A* | 18,19 | W0574<br>W1747 | Malaya<br>India | W0574<br>W1747 |
| *Or3* | 18,19 | W3042<br>W3048<br>W3073 | China<br>China<br>China | W3042<br>W3048<br>W3073 |

**Table 4. Same as Table 3 but the focus is on data sets 20 and 21. Data sets 20 and 21 are built from data sets 18 and 19: these data sets contain a mixture of varieties from data sets 18 and 19.**

| Subpopulations | Dataset | Variety ID | Country | Variety name |
|---|---|---|---|---|
| *circum Aus* | 20 | IRIS-313-11058 | Bangladesh | AUS 329 |
| | | IRIS-313-10852 | India | ARC 7336 |
| | 21 | IRIS 313-11737 | India | CHUNDI |
| | | IRIS-313-11027 | Pakistan | JHONA 101 |
| *circum Basmati* | 20 | IRIS-313-11062 | Bangladesh | BEGUNBICHI 33 |
| | | IRIS-313-11825 | India | HANSRAJ |
| | | IRIS-313-8326 | India | JC1 |
| | 21 | IRIS-313-11258 | India | ARC 13502 |
| | | IRIS-313-10851 | India | ARC 7296 |
| | | IRIS-313-12094 | Bangladesh | ARC KASHA |
| *Indica* | 20 | IRIS-313-11819 | Myanmar | PADINTHUMA |
| | | IRIS-313-11796 | China | DU GEN CHUAN |
| | | IRIS-313-11741 | SriLanka | HERATH BANDA |
| | 21 | IS-313-11646 | India | NCS771 A |
| | | CX270 | Taiwan | TAICHUNGNATIVE1 |
| | | IRIS-313-11089 | Cambodia | SRAU THMOR |
| *Japonica* | 20 | B204 | China | LONGHUAMAOHU |
| | | IRIS-313-11691 | Bhutan | SHANGYIPA |
| | | IRIS-313-10577 | Philippines | IFUGAO RICE |
| | 21 | IRIS-313-11924 | Thailand | NAM JAM |
| | | IRIS-313-7883 | Indonesia | GANIGI |
| | | B269 | China | YUEFU |
| *Or1I* | 20,21 | W1117 | India | W1117 |
| | | W1559 | Thailand | W1559 |
| *Or1A* | 20,21 | W0574 | Malaya | W0574 |
| | | W1747 | India | W1747 |
| *Or3* | 20,21 | W3042 | China | W3042 |
| | | W3048 | China | W3048 |
| | | W3073 | China | W3073 |

**Table 5. Cultivars contained in data set 22. Data set 22 is used in the ABC-RF study.**

| Subpopulations | Variety ID | Country | Variety name |
|---|---|---|---|
| *circum Aus* | IRIS-313-11058 | Bangladesh | AUS 329 |
| | IRIS-313-10852 | India | ARC 7336 |
| | IRIS 313-11737 | India | CHUNDI |
| | IRIS-313-11027 | Pakistan | JHONA 101 |
| | IRIS-313-11191 | Srilanka | RANRUWAN |
| | CX227 | Japon/Banglasdesh | KASALATH |
| | IRIS-313-10020 | Srilanka | HODARAWALA |
| | IRIS-313-10718 | Srilanka | KARUTHA SEENATI |
| | IRIS-313-10871 | India | ARC 11777 |
| | IRIS-313-11055 | Banglasdesh | AUS 299 |
| *circum Basmati* | IRIS-313-11062 | Bangladesh | BEGUNBICHI 33 |
| | IRIS-313-11825 | India | HANSRAJ |
| | IRIS-313-8326 | India | JC1 |
| | IRIS-313-11258 | India | ARC 13502 |
| | IRIS-313-10851 | India | ARC 7296 |
| | IRIS-313-12094 | Bangladesh | ARC KASHA |
| | CX149 | India | KARNAL LOCAL |
| | IRIS-313-11270 | India | ARC 14663 |
| | IRIS-313-11289 | India | ARC 18578 |
| | CX110 | Japan | UP15 |
| *Indica* | IRIS-313-11819 | Myanmar | PADINTHUMA |
| | IRIS-313-11796 | China | DU GEN CHUAN |
| | IRIS-313-11741 | SriLanka | HERATH BANDA |
| | IS-313-11646 | India | NCS771 A |
| | CX270 | Taiwan | TAICHUNGNATIVE1 |
| | IRIS-313-11089 | Cambodia | SRAU THMOR |
| | IRIS-313-11665 | China | JIN HUA 258 |
| | CX150 | Philippines | Chorofa |
| | IRIS-313-10975 | Bangladesh | KOLA MUCHI |
| | IRIS-313-11118 | Vietnam | SOC NAU(BIEN THE) |
| *Japonica* | B204 | China | LONGHUAMAOHU |
| | IRIS-313-11691 | Bhutan | SHANGYIPA |
| | IRIS-313-10577 | Philippines | IFUGAO RICE |
| | IRIS-313-11924 | Thailand | NAM JAM |
| | IRIS-313-7883 | Indonesia | GANIGI |
| | B269 | China | YUEFU |
| | B001 | China | HEIBIAO |
| | IRIS-313-12349 | Laos | MAK KHEUA DENG |
| | IRIS-313-10582 | Philippines | ARC 7336 |
| | IRIS-313-11044 | Malaysia | BUKU |

**Table 6. Wild varieties contained in data set 22. Data set 22 is used in the ABC-RF study.**

| Subpopulations | Variety ID | Country | Variety name |
|---|---|---|---|
| *Or1I* | W1117 | India | W1117 |
| | W1559 | Thailand | W1559 |
| | W0178 | Thailand | W0178 |
| | W2316 | Vietnam | W2316 |
| | W2061 | Bangladesh | W2061 |
| | W1723 | China | W1723 |
| | W0639 | Burma | W0639 |
| | W0638 | Burma | W0638 |
| *Or1A* | W1740 | India | W1740 |
| | W0106 | India | W0106 |
| | W0574 | Malaya | W0574 |
| | W1105 | India | W1105 |
| | W1747 | India | W1747 |
| | W1853 | Thailand | W1853 |
| | W1551 | Thailand | W1551 |
| | W1690 | Thailand | W1690 |
| *Or3* | W3065 | China | W3065 |
| | W0621 | Burma | W0621 |
| | W1782 | India | W1782 |
| | W1739 | India | W1739 |
| | W3073 | China | W3073 |
| | W3048 | China | W3048 |
| | W3042 | China | W3042 |
| | W3037 | China | W3037 |

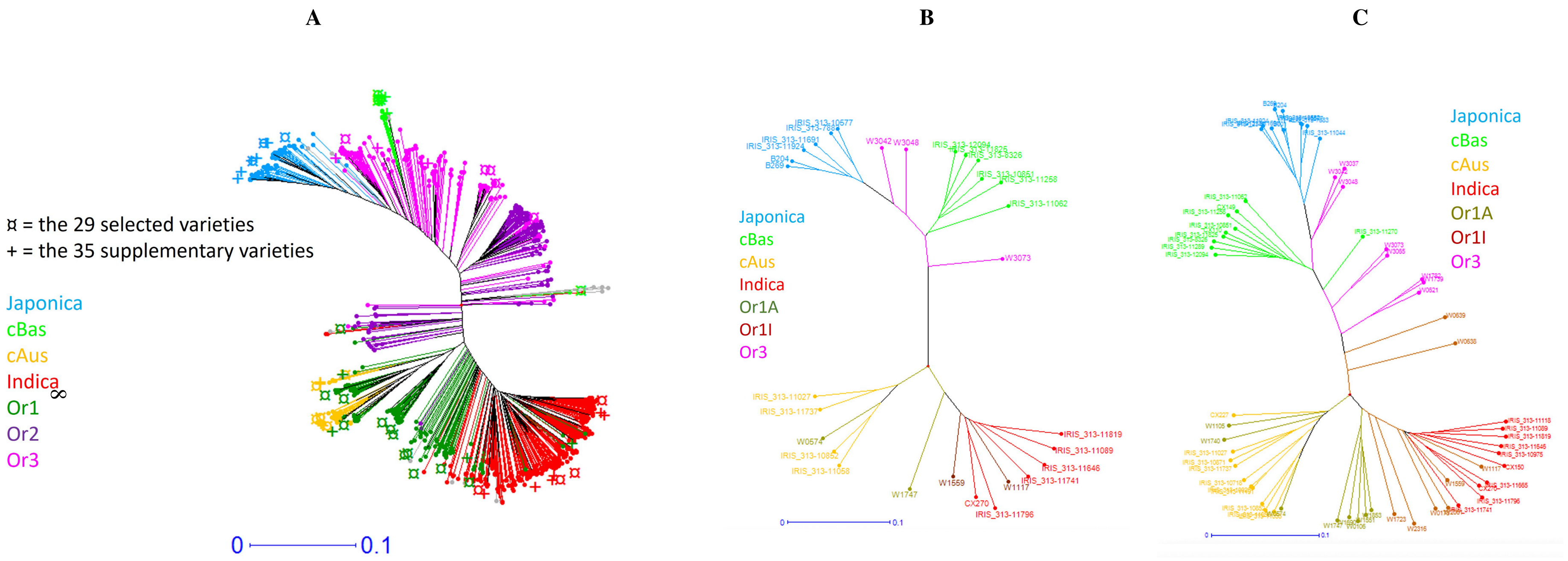


**Figure 1.** Same as Figure 2 of the main manuscript except that the names of the accessions are specified. A: the 899 accessions based on 2.48 million SNPs as described in [9]. B: the 29 selected accessions. C: the 64 selected accessions.

### 2 Free exploration of the topology space with SNAPPNET

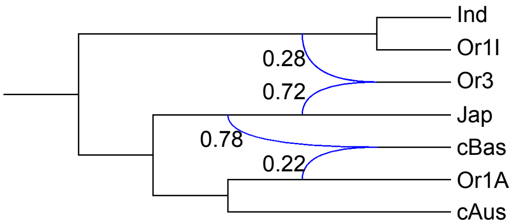


**First topology, first sampling**
49.52%

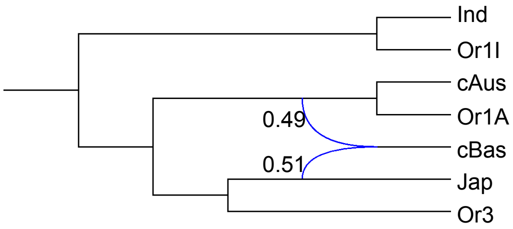


**Second topology, first sampling**
47.10%

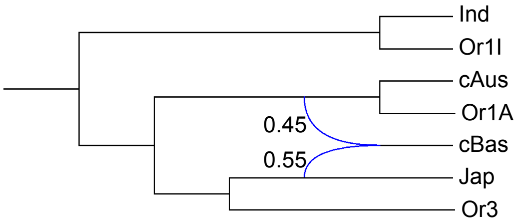


**Single topology, second sampling**
100%

**Figure 2.** The main topologies sampled by SNAPPNET when **data set 1** was analyzed. Reported inheritance probabilities for each topology are averages on sampled observations. The two different samplings of 12k markers along the rice genome are considered here.

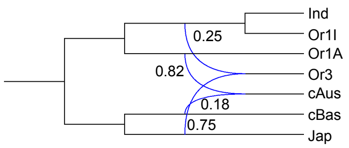


**First topology, first sampling**
80.42%

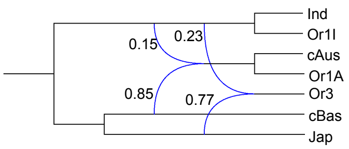


**Second topology, first sampling**
15.08%

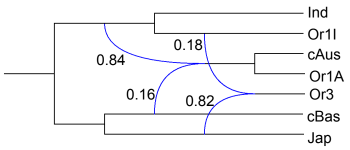


**Third topology, first sampling**
2.16%%

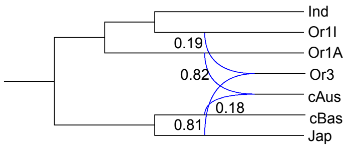


**Fourth topology, first sampling**
1.11%

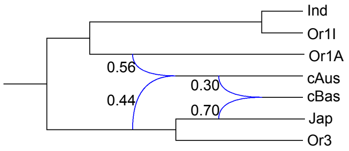


**Fifth topology, first sampling**
1.02%

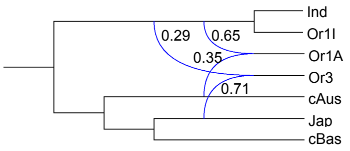


**Sixth topology, first sampling**
17.14%

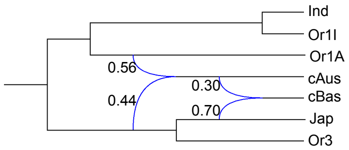


**First topology, second sampling**
47.63%

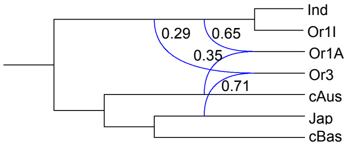


**Second topology, second sampling**
17.14%

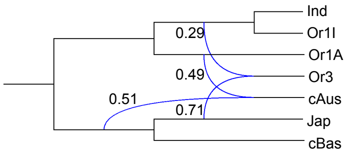


**Third topology, second sampling**
14.80%

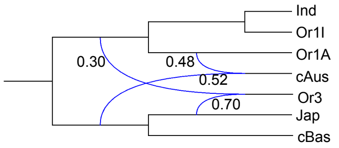


**Fourth topology, second sampling**
11.75%

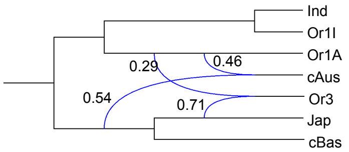


**Fifth topology, second sampling**
2.04%

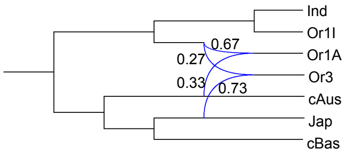


**Sixth topology, second sampling**
1.96%

**Figure 3.** The topologies sampled by SnappNet when **data set 2** was analyzed. Reported inheritance probabilities for each topology are averages on sampled observations. The two different samplings of 12k markers along the rice genome are considered here.

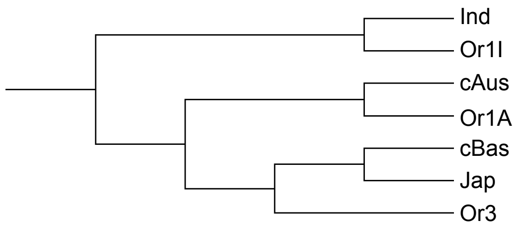


**First topology, first sampling**
92.53%

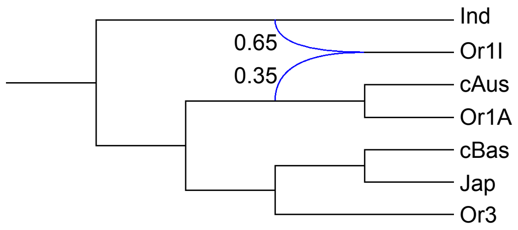


**Second topology, first sampling**
7.39%

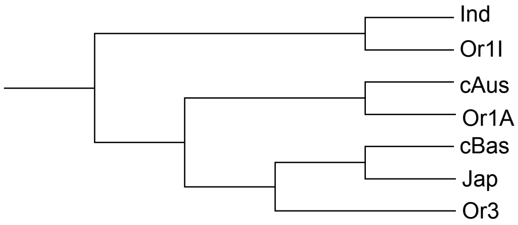


**Single topology, second sampling**
100%

**Figure 4.** The main topologies sampled by SnappNet when **data set 3** was analyzed. Reported inheritance probabilities for each topology are averages on sampled observations. The two different samplings of 12k markers along the rice genome are considered here.

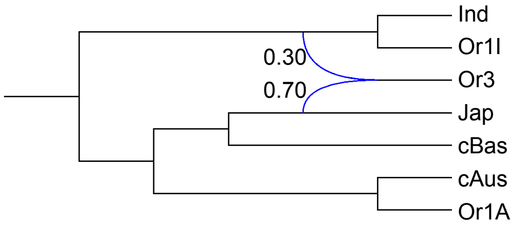


**Main topology, first sampling**
99.41%

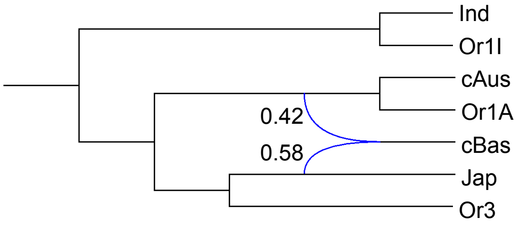


**First topology, second sampling**
50%

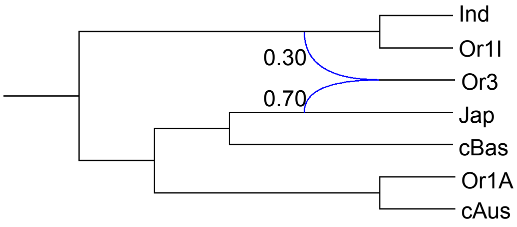


**Second topology, second sampling**
50%

**Figure 5.** The main topologies sampled by SnappNet when **data set 4** was analyzed. Reported inheritance probabilities for each topology are averages on sampled observations. The two different samplings of 12k markers along the rice genome are considered here.

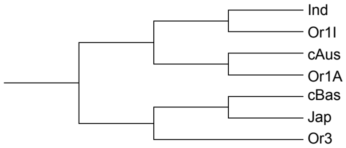


**First topology, First sampling**
92.53%

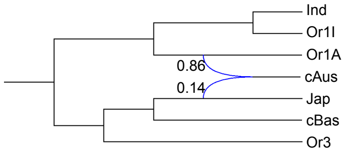


**Second topology, first sampling**
3.78%

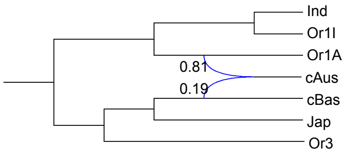


**First topology, second sampling**
46.91%

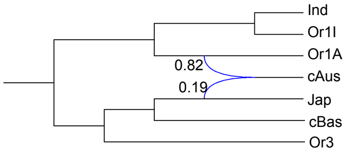


**Second topology, second sampling**
40.20%

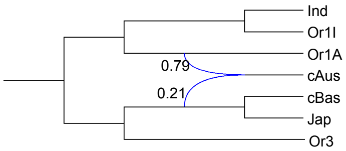


**Third topology, second sampling**
12.23%

**Figure 6.** The main topologies sampled by SNAPPNET when **data set 5** was analyzed. Reported inheritance probabilities for each topology are averages on sampled observations. The two different samplings of 12k markers along the rice genome are considered here.

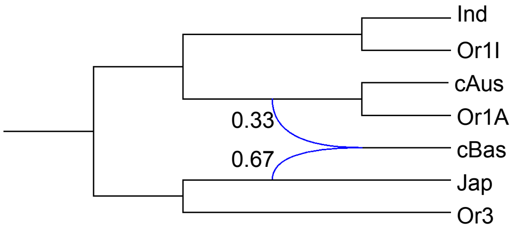


**Single topology, first sampling**
100%

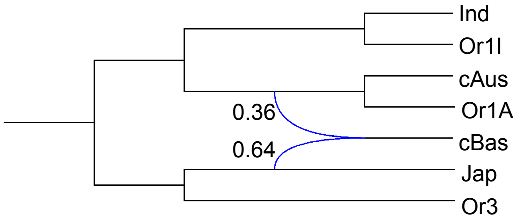


**Main topology, second sampling**
99.70%

**Figure 7.** The main topologies sampled by SNAPPNET when **data set 6** was analyzed. Reported inheritance probabilities for each topology are averages on sampled observations. The two different samplings of 12k markers along the rice genome are considered here.

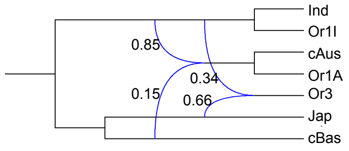


**First topology, first sampling**
55.01%

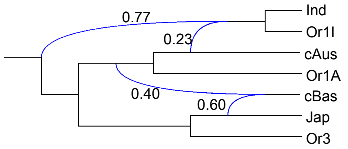


**Second topology, first sampling**
26.36%

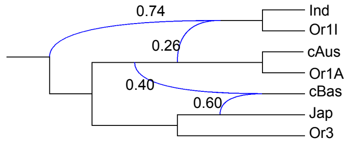


**Third topology, first sampling**
10.99%

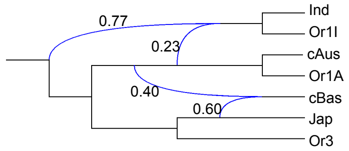


**Fourth topology, first sampling**
5.61%

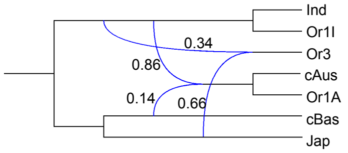


**Fifth topology, first sampling**
1.55%

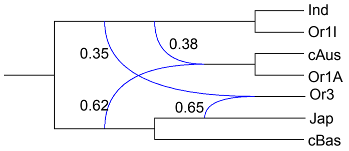


**First topology, second sampling**
71.92%

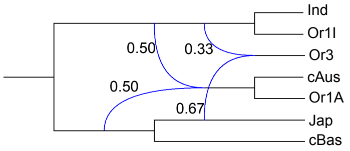


**Second topology, second sampling**
18.11%

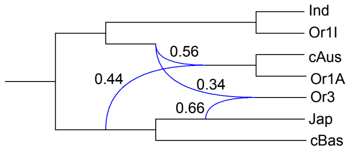


**Third topology, second sampling**
5.80%

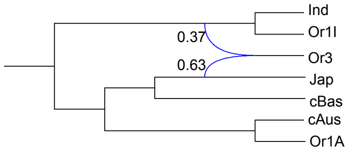


**Fourth topology, second sampling**
2.62%

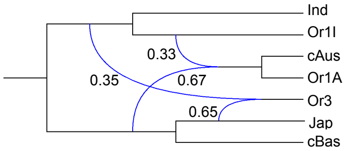


**Fifth topology, second sampling**
1.28%

**Figure 8.** The main topologies sampled by SnappNet when **data set 7** was analyzed. Reported inheritance probabilities for each topology are averages on sampled observations. The two different samplings of 12k markers along the rice genome are considered here.

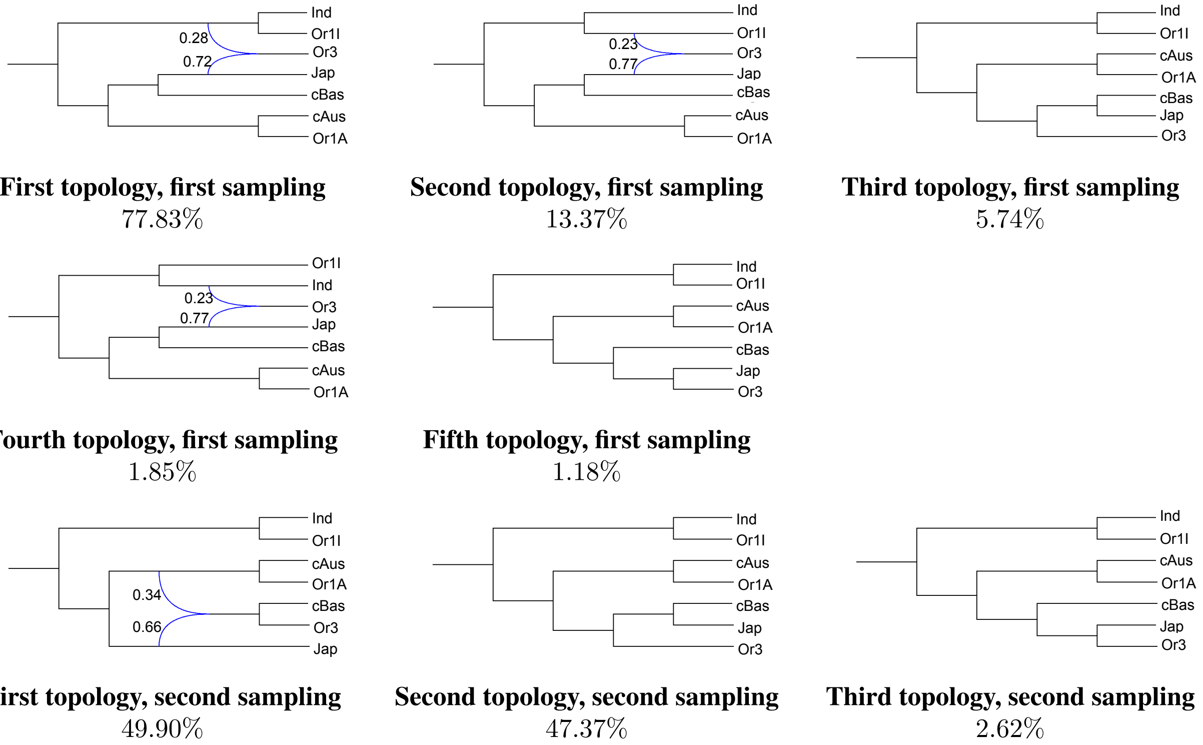


**First topology, first sampling**
77.83%

**Second topology, first sampling**
13.37%

**Third topology, first sampling**
5.74%

**Fourth topology, first sampling**
1.85%

**Fifth topology, first sampling**
1.18%

**First topology, second sampling**
49.90%

**Second topology, second sampling**
47.37%

**Third topology, second sampling**
2.62%

**Figure 9.** The main topologies sampled by SnappNet when **data set 8** was analyzed. Reported inheritance probabilities for each topology are averages on sampled observations. The two different samplings of 12k markers along the rice genome are considered here.

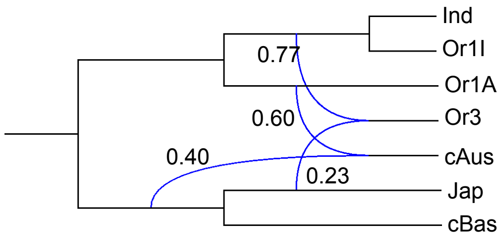


**First topology, first sampling**
71.72%

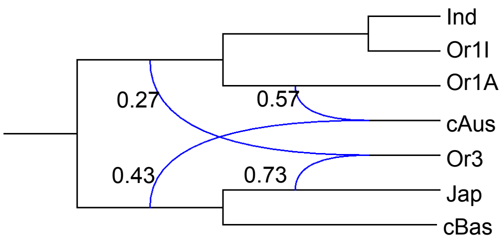


**Second topology, first sampling**
26.90%

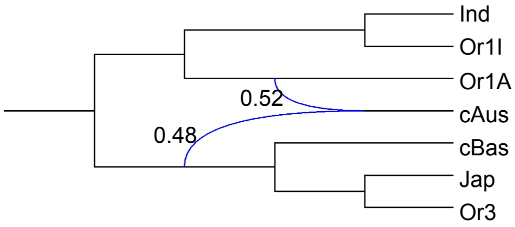


**Single topology, second sampling**
100%

**Figure 10.** The main topologies sampled by SnappNet when **data set 9** was analyzed. Reported inheritance probabilities for each topology are averages on sampled observations. The two different samplings of 12k markers along the rice genome are considered here.

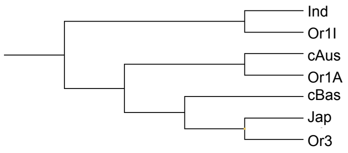


**Topology, first sampling**
99.86%

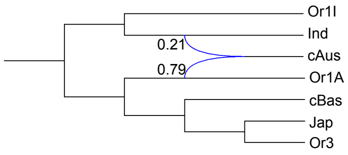


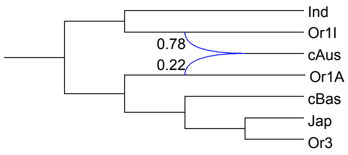


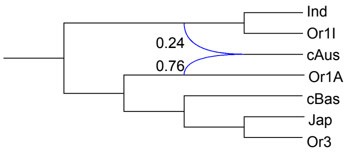


**First topology, second sampling** 86.40%

**Second topology, second sampling** 9.23%

**Third topology, second sampling** 4.07%

**Figure 11.** The main topologies sampled by SnappNet when **data set 10** was analyzed. Reported inheritance probabilities for each topology are averages on sampled observations. The two different samplings of 12k markers along the rice genome are considered here.

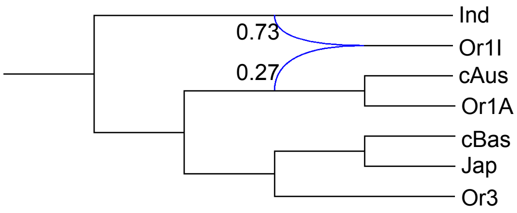


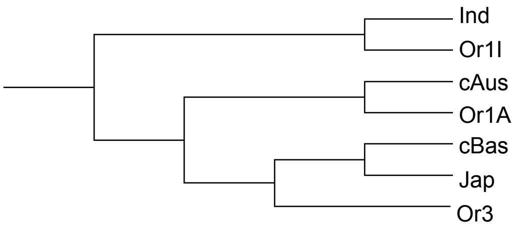


**Main topology, first sampling** 99.45%

**Single topology, second sampling** 100%

**Figure 12.** The main topologies sampled by SnappNet when **data set 11** was analyzed. Reported inheritance probabilities for each topology are averages on sampled observations. The two different samplings of 12k markers along the rice genome are considered here.

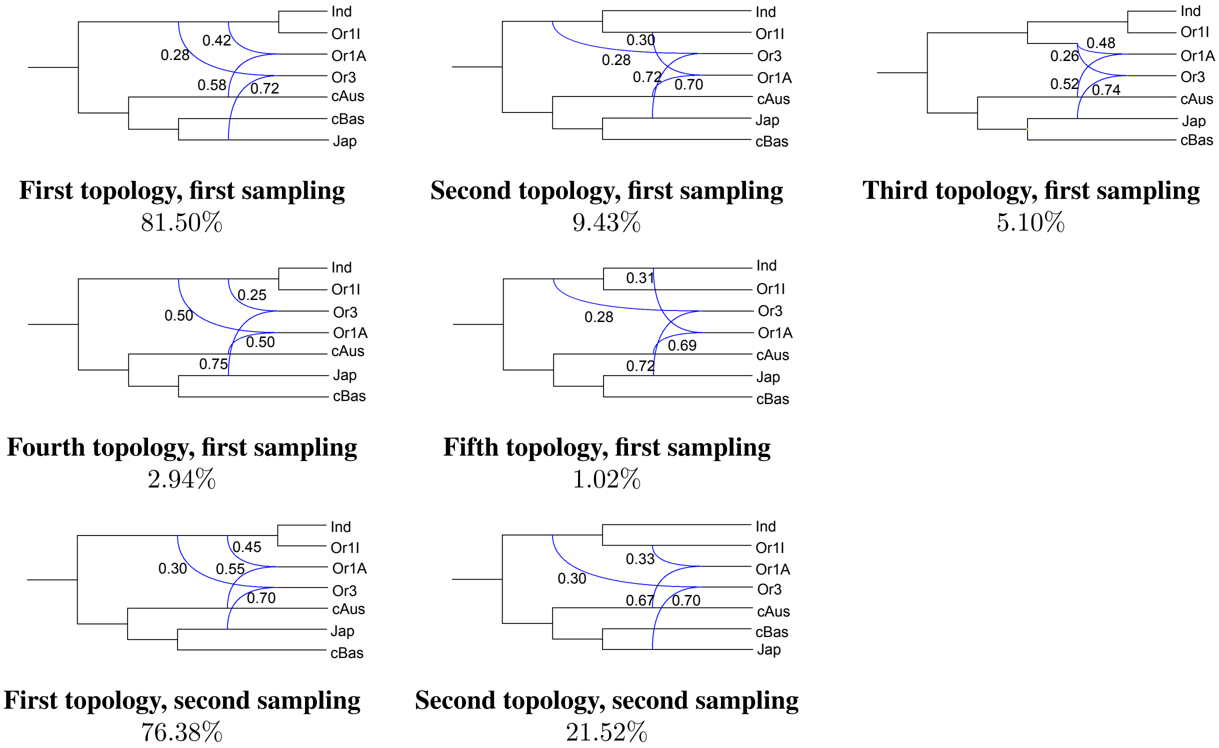


**Figure 13.** The topologies sampled by SnappNet when **data set 12** was analyzed. Reported inheritance probabilities for each topology are averages on sampled observations. The two different samplings of 12k markers along the rice genome are considered here.

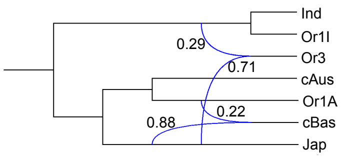


**First topology, first sampling**
87.71%

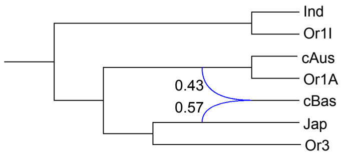


**Second topology, first sampling**
3.40%

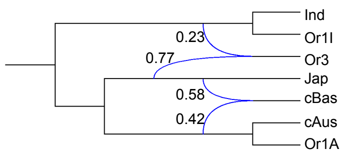


**Third topology, first sampling**
2.97%

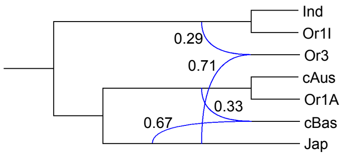


**Fourth topology, first sampling**
2.14%

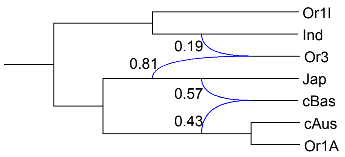


**Fifth topology, first sampling**
1.77%

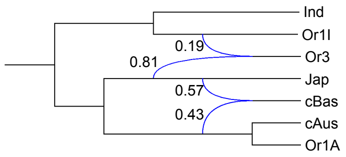


**Sixth topology, first sampling**
1.08%

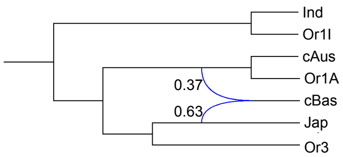


**First topology, second sampling**
48.99%

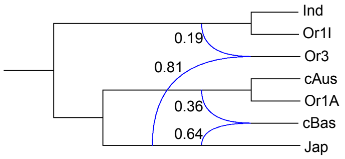


**Second topology, second sampling**
15.06%

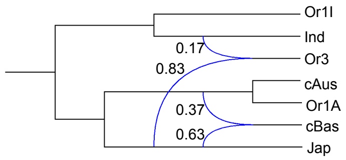


**Third topology, second sampling**
15.06%

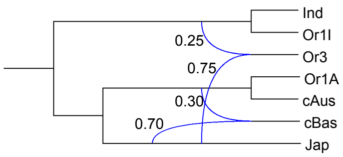


**Fourth topology, second sampling**
7.73%

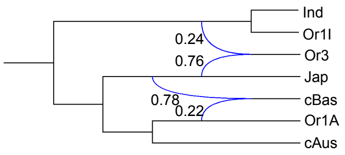


**Fifth topology, second sampling**
6.66%

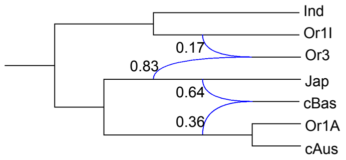


**Sixth topology, second sampling**
3.92%

**Figure 14.** The topologies sampled by SnappNet when **data set 13** was analyzed. Reported inheritance probabilities for each topology are averages on sampled observations. The two different samplings of 12k markers along the rice genome are considered here.

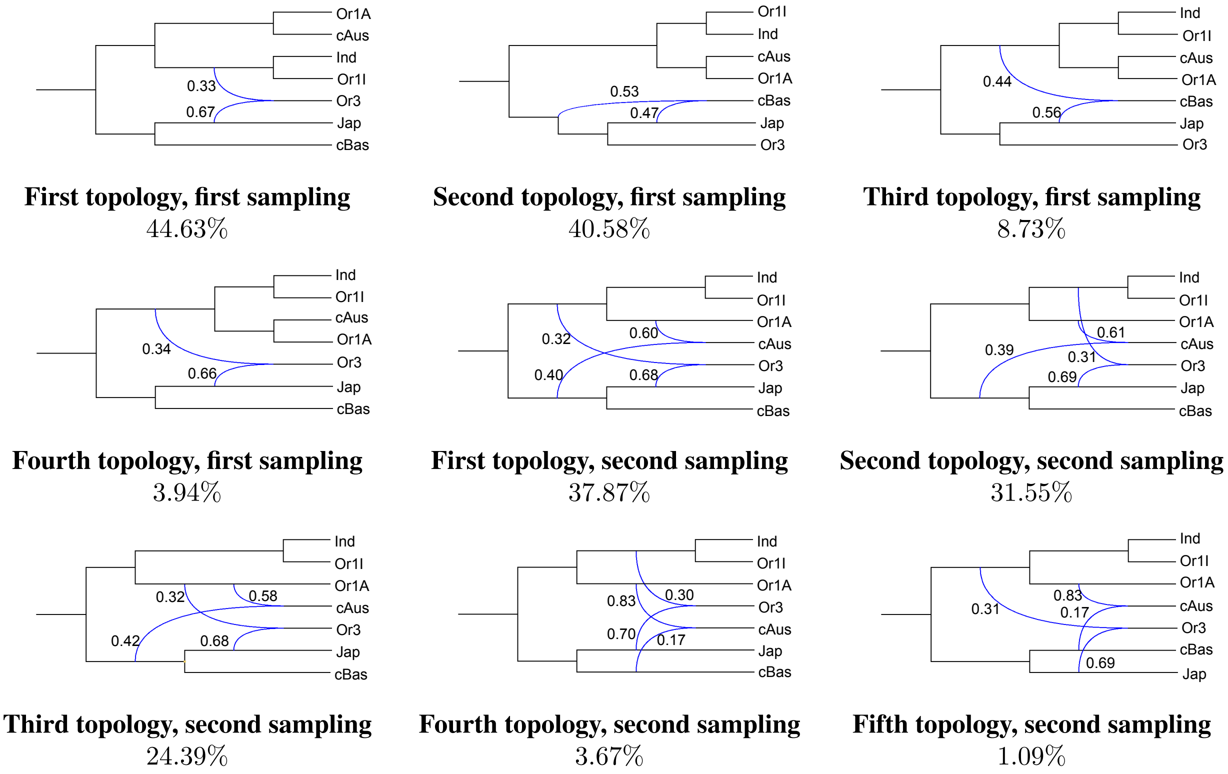


**Figure 15.** The topologies sampled by SnappNet when **data set 14** was analyzed. Reported inheritance probabilities for each topology are averages on sampled observations. The two different samplings of 12k markers along the rice genome are considered here.

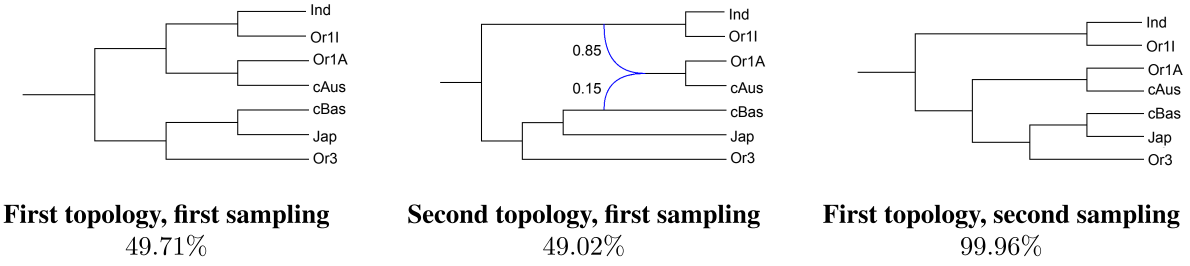


**Figure 16.** The topologies sampled by SnappNet when **data set 15** was analyzed. Reported inheritance probabilities for each topology are averages on sampled observations. The two different samplings of 12k markers along the rice genome are considered here.

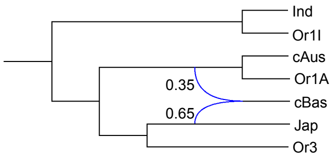


**First topology, first sampling**
95.72%

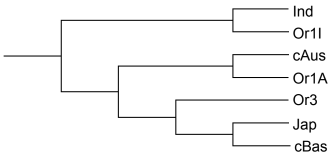


**Second topology, first sampling**
3.98%

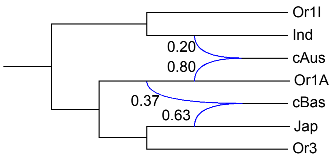


**First topology, second sampling**
78.99%

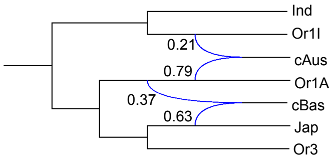


**Second topology, second sampling**
10.41%

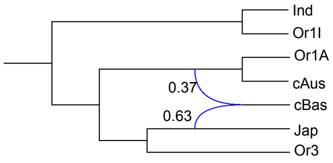


**Third topology, second sampling**
4.62%

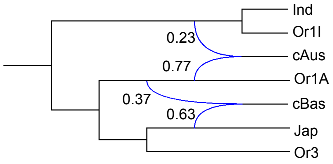


**Fourth topology, second sampling**
4.13%

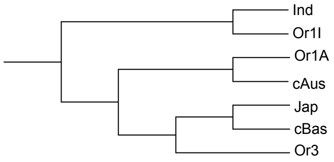


**Fifth topology, second sampling**
1.77%

**Figure 17.** The topologies sampled by SNAPPNET when **data set 16** was analyzed. Reported inheritance probabilities for each topology are averages on sampled observations. The two different samplings of 12k markers along the rice genome are considered here.

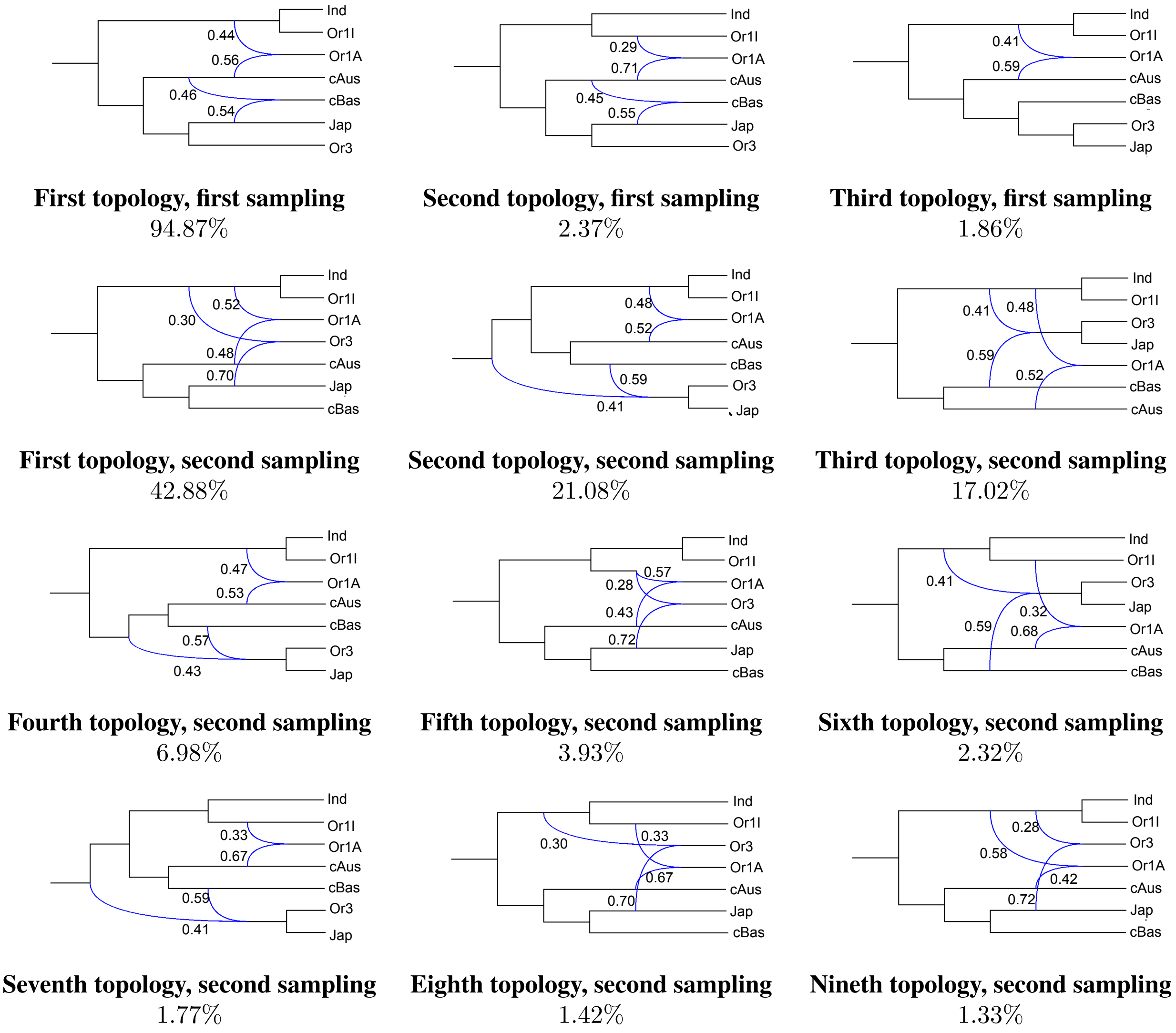


**First topology, first sampling** 94.87%

**Second topology, first sampling** 2.37%

**Third topology, first sampling** 1.86%

**First topology, second sampling** 42.88%

**Second topology, second sampling** 21.08%

**Third topology, second sampling** 17.02%

**Fourth topology, second sampling** 6.98%

**Fifth topology, second sampling** 3.93%

**Sixth topology, second sampling** 2.32%

**Seventh topology, second sampling** 1.77%

**Eighth topology, second sampling** 1.42%

**Nineth topology, second sampling** 1.33%

**Figure 18.** The topologies sampled by SNAPPNET when **data set 17** was analyzed. Reported inheritance probabilities for each topology are averages on sampled observations. The two different samplings of 12k markers along the rice genome are considered here.

### 3 Study of 16 evolutionary scenarios with SNAPPNET

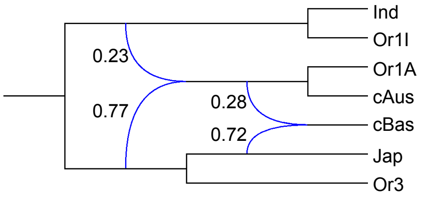


**Network 12, data set 18**

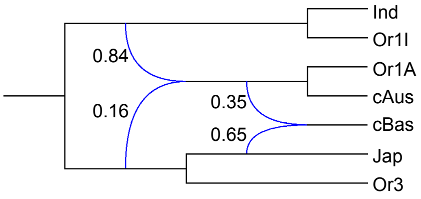


**Network 12, data set 19**

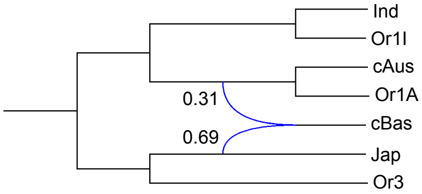


**Network 1, data set 18**

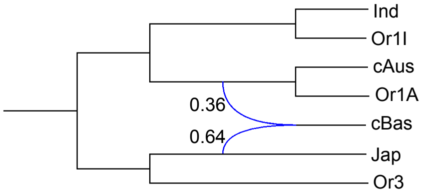


**Network 1, data set 19**

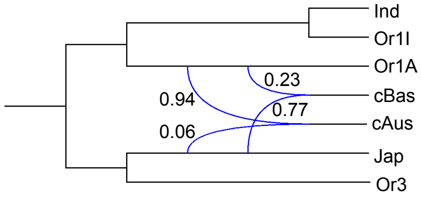


**Network 15, data set 18**

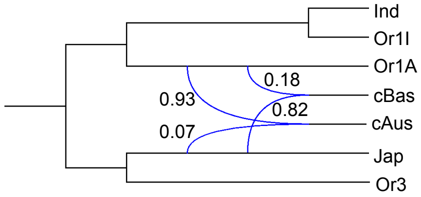


**Network 15, data set 19**

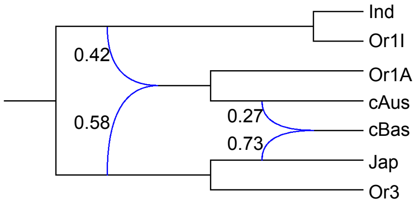


**Network 8, data set 18**

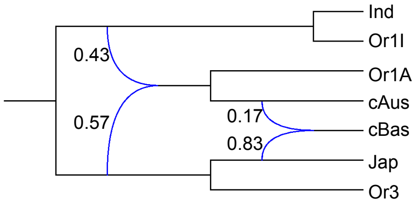


**Network 8, data set 19**

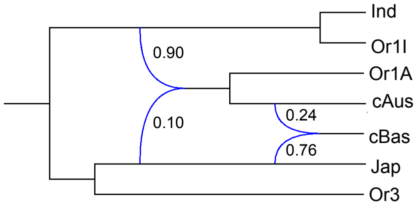


**Network 9, data set 18**

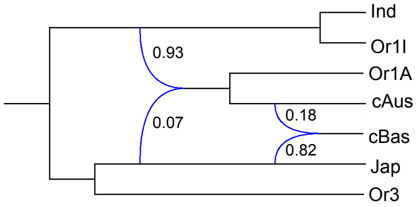


**Network 9, data set 19**

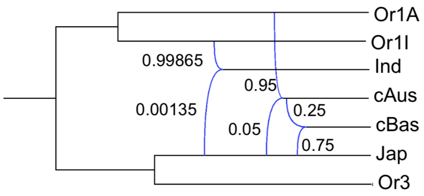


**Network 16, data set 18**

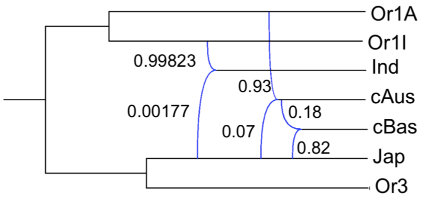


**Network 16, data set 19**

**Figure 19.** Inheritance probabilities estimated by SNAPPNET for the most likely evolutionary scenarios. The focus is on data sets 18 and 19. For each data set, reported inheritance probabilities are averages on the two samplings.

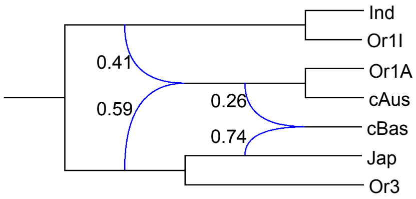


**Network 12, data set 20**

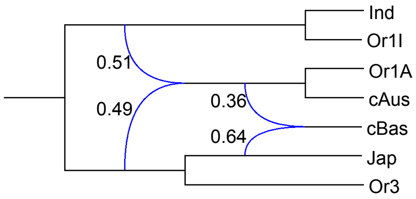


**Network 12, data set 21**

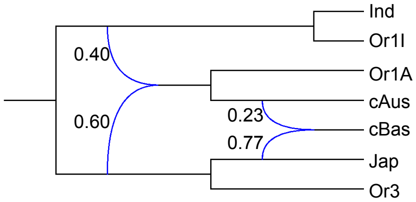


**Network 8, data set 20**

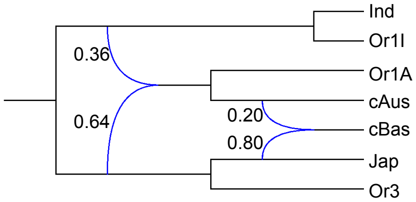


**Network 8, data set 21**

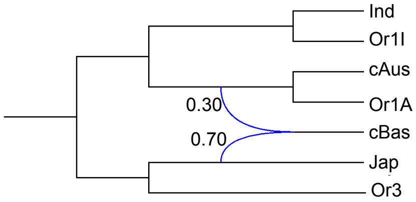


**Network 1, data set 20**

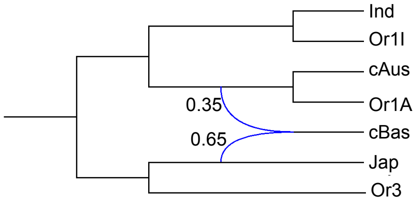


**Network 1, data set 21**

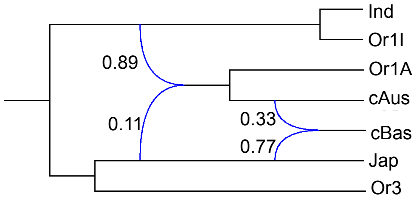


**Network 9, data set 20**

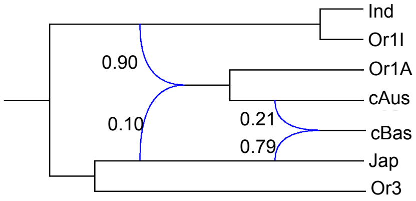


**Network 9, data set 21**

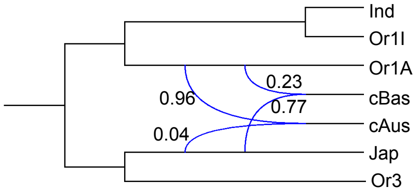


**Network 15, data set 20**

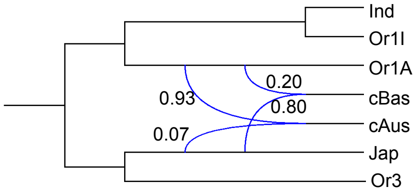


**Network 15, data set 21**

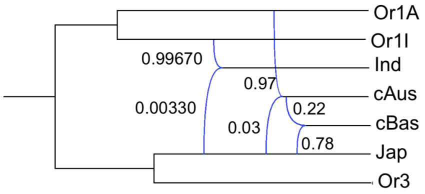


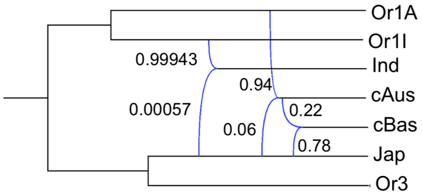




**Network 16, data set 20**

**Network 16, data set 21**

**Figure 20.** Same as Figure 19 except that the focus is on data sets 20 and 21.

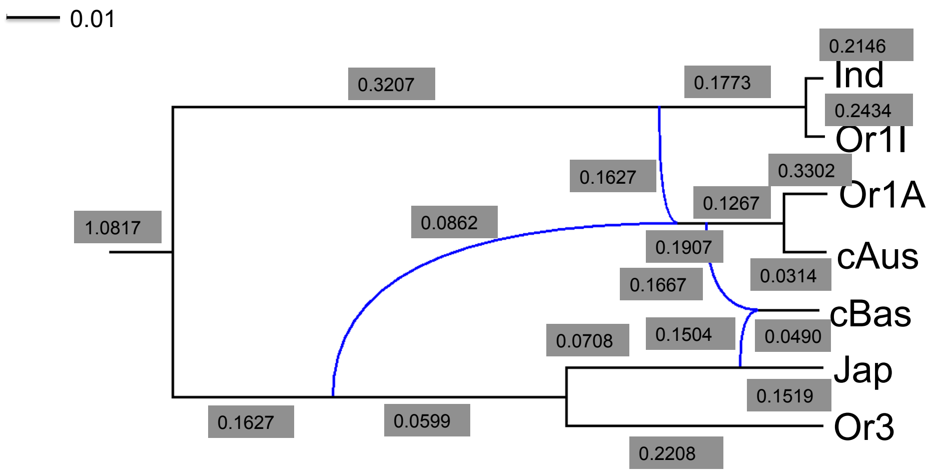


**Figure 21.** Population sizes and branch lengths estimated by SNAPPNET for **Network 12**. Population sizes are given in grey boxes, whereas branch lengths are are given in units of expected number of mutations per site (see the scale at the top left). Reported values are averages on data sets 18, 19, 20 and 21, and on the two samplings.

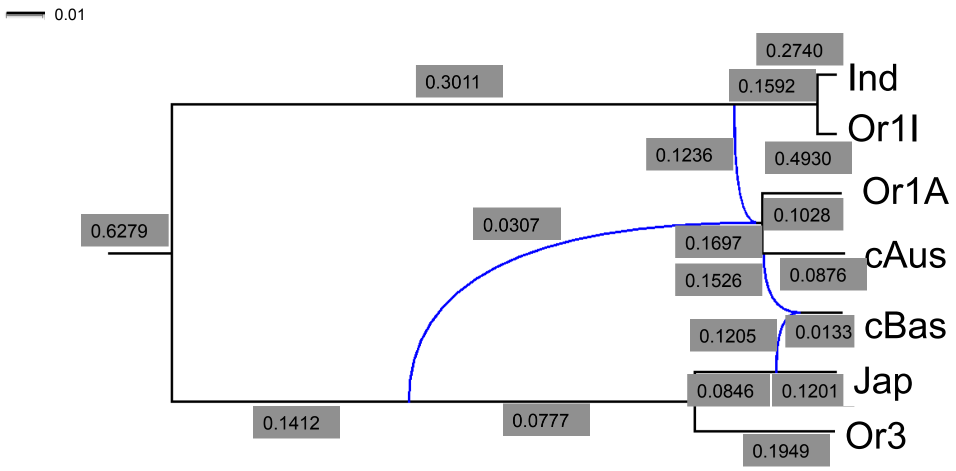


**Figure 22.** Population sizes and branch lengths estimated by SNAPPNET for **Network 8**. Population sizes are given in grey boxes, whereas branch lengths are are given in units of expected number of mutations per site (see the scale at the top left). Reported values are averages on data sets 18, 19, 20 and 21, and on the two samplings.

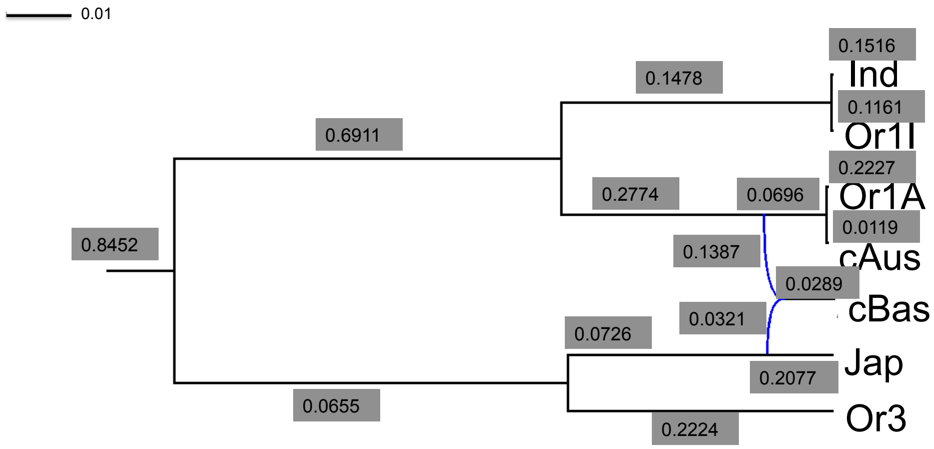


**Figure 23.** Population sizes and branch lengths estimated by SnappNet for **Network 1**. Population sizes are given in grey boxes, whereas branch lengths are are given in units of expected number of mutations per site (see the scale at the top left). Reported values are averages on data sets 18, 19, 20 and 21, and on the two samplings.

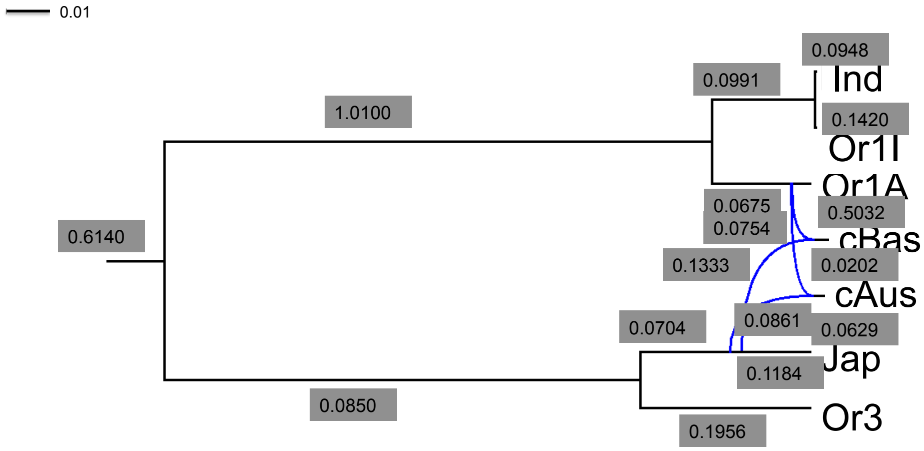


**Figure 24.** Population sizes and branch lengths estimated by SnappNet for **Network 15**. Population sizes are given in grey boxes, whereas branch lengths are are given in units of expected number of mutations per site (see the scale at the top left). Reported values are averages on data sets 18, 19, 20 and 21, and on the two samplings.

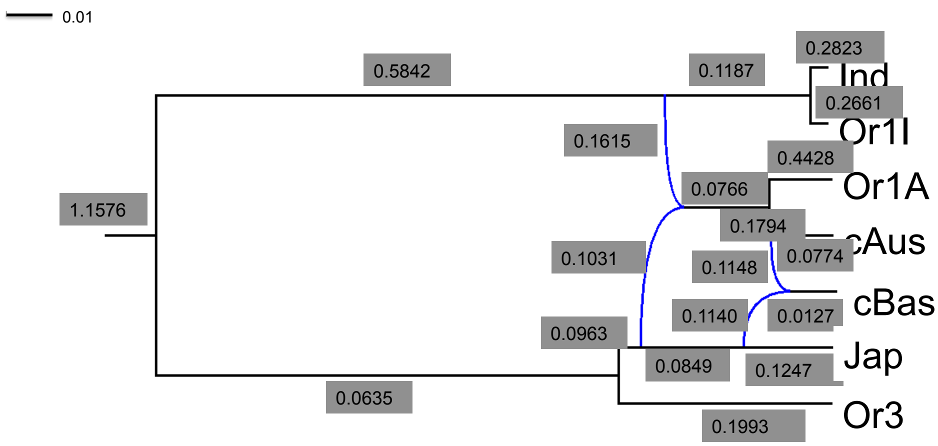


**Figure 25.** Population sizes and branch lengths estimated by SNAPPNET for **Network 9**. Population sizes are given in grey boxes, whereas branch lengths are are given in units of expected number of mutations per site (see the scale at the top left). Reported values are averages on data sets 18, 19, 20 and 21, and on the two samplings.

**Table 7. Rates $u$ and $v$ estimated by Maximum Likelihood with SNAPPNET. Average over Data sets 18-21 (including the two samplings). The focus is on the six best scenarios.**

| Network | $u$ | $v$ |
|---|---|---|
| 1 | 0.554975 | 5.04750 |
| 8 | 0.55405 | 5.12530 |
| 9 | 0.55457 | 5.08090 |
| 12 | 0.55393 | 5.13499 |
| 15 | 0.55450 | 5.08716 |
| 16 | 0.55430 | 5.10405 |

### 4 Extra informations on SS6-18 and SS19-22

#### 4.1 Computing SS6-18

As mentioned in the main manuscript, there are $\binom{7}{4}$=35 sets of 4 species chosen among 7 species. Let us enumerate these different sets of 4 species:

- $\binom{4}{4} = 1$ set with only cultivars
- $\binom{4}{1} = 4$ sets with one cultivar and three wilds,
- $\binom{4}{2} \times \binom{3}{2} = 18$ sets with 2 cultivars and two wilds
- $\binom{4}{3} \times \binom{3}{1} = 12$ sets with 3 cultivars and one wild.

Recall that data set 22 includes 10 varieties for each of the four cultivar subpopulations and 8 varieties for each of the three wild subpopulations. Therefore, there is a total of $\binom{4}{4} \times 10^4 + \binom{4}{1} \times 10 \times 8^3 + \binom{4}{1} \times \binom{2}{3} \times 10^2 \times 8^2 + \binom{4}{1} \times \binom{2}{3} \times 10^3 \times 8 = 241,680$ possible sets of 4 varieties. Furthermore, following [4], let us assume that the outgroup $O$ is fixed. There are 3 possibilities for labelling the hybrid species (either $P_1$, $P_2$ or $H$) of the network in Figure 4 of the main manuscript. Then, the total number of possibilities is $241,680 \times 3 = 725,040$. As described in the main manuscript, we handled $\binom{7}{4} = 35$ species sets and 625 combinations of varieties for each set, that is to say a total of $35 \times 625 = 21,875$ cases. Overall, the ratio between our sampling and all possibilities is $\frac{21,875}{725,040} \approx 3.01\%$. Note that this ratio becomes $\frac{21,875}{241,680 \times 2} = 4.53\%$ as soon as we consider only two ways of labelling the hybrid species, as suggested in [4].

#### 4.2 Computing SS19-22

In the same way, there are $\binom{7}{5}$=21 sets of 5 species chosen among 7 species. Let us enumerate these different sets of 5 species:

- $\binom{4}{4} \times \binom{3}{1} = 3$ set with 4 cultivars and one wild
- $\binom{4}{3} \times \binom{3}{2} = 12$ sets with 3 cultivars and two wilds
- $\binom{4}{2} \times \binom{3}{3} = 6$ sets with 2 cultivars and three wilds

There is a total of $\binom{4}{4}\binom{3}{1} \times 10^4 \times 8 + \binom{4}{3} \times \binom{3}{2} \times 10^3 \times 8^2 + \binom{4}{2} \times \binom{3}{3} \times 10^2 \times 8^3 = 1,315,200$ possible sets of 5 varieties. Assuming that the outgroup $O$ is fixed, the 3 possible ways of labelling the leaves of the species tree (Figure 4 of the main manuscript) are : $((P_1, P_2), (P_3, P_4))$, $((P_1, P_3), (P_2, P_4))$ and $((P_1, P_4), (P_2, P_3))$. Then, there are $1,315,200 \times 3 = 3,945,600$ combinations. As explained in the main manuscript, we handled $\binom{7}{5} = 21$ species sets and $3,125$ combinations of varieties for each set, that is to say we considered a total of $21 \times 3,125 = 65,625$ cases. Overall, the ratio between our sampling and all possibilities is $\frac{65,625}{3,945,600} = 1.66\%$.

### 5 Discriminating between the $6$ best scenarios with the SNARF hybrid method

##### Preliminaries

As mentioned before, although SNAPPNET tremendously reduced the computational burden as compared to `MCMCBiMarkers`, the likelihood computation can still be costly with SNAPPNET as soon as we consider many varieties per species (cf. [7]). In contrast, ABC-RF allows to analyze large data sets. First, the problem's dimension is transformed from $n \times p$ to $N_{\text{Ref}} \times$ nbSS, where $n$, $p$, $N_{\text{Ref}}$ and nbSS denote respectively the number of varieties, the number of SNPs, the number of rows of the reference table and the total number of SS. Recall that the number of rows denote the total number of simulations (cf. Section 2.5.3 of the main text). In our case, the dimension changed from $64\times$12,000 to 84,000$\times 562$. Moreover, when the number of variables is large, RF is relatively robust to noisy variables and RF does not deteriorate as soon as the fraction of relevant variables is large enough (cf. [3]). In that sense, ABC-RF can easily handle many SS in a satisfactory manner.

In our study, the reference tables, associated to the different combination of priors and required for the ABC-RF learning step, were obtained thanks to the SIMSNAPPNET simulator [7], in which we implemented specific SS (cf. Section 2.5 of the main text). The SS we propose are mainly based on phylogenetic invariants ([4, 6]) and are dedicated to phylogenetic networks. Besides, they are computed under the NMSC model.

##### Performances of the SNARF hybrid approach on simulated data

In machine learning, a confusion matrix is a tool that enables the evaluation of the performance of a classifier: each row of the matrix refers to the true class whereas each column refers to the the predicted class (see [3]). The confusion matrices obtained by ABC-RF on the basis of reference tables associated to the different priors, are given in Tables 8-15. Each table reports the number of votes for each network, on the basis of 1,000 trees. Moreover, it also gives the so-called misclassification rate. This rate refers to the proportion of incorrectly classified objects: it can be viewed as the prior error rate of the classifier. Table 16 is a summary of all the investigated priors. In what follows, we study the impact of different factors on the performance of our hybrid approach.

Impact of the inheritance prior:
Two kinds of prior for the inheritance probability $\gamma$, were considered: the prior is either (a) a specific uniform prior (Table 8) based on SNAPPNET, or (b) a general uniform prior (Table 9) "free" from SNAPPNET (cf. Section 2.5.1 of the main text).

To begin with, let us focus on the specific uniform prior (Table 8). The misclassification rate was found to be equal to $16.13\%$ and $14.30\%$, for Networks 8 and 9, respectively. In contrast, the error rates for Networks 1, 12, 15 and 16 were found equal to $4.28\%$, $1.44\%$, $1.30\%$ and $1.85\%$, respectively. Then, Networks 8 and 9 seem to be hard to discriminate, whereas Networks 1, 12, 15 and 16 were easily recovered by ABC-RF. It can be explained by the fact that the evolutionary scenarios underlined by Networks

8 and 9 are very close. The two networks present a topology with two hybridization nodes on top of each other, and have the same number of edges. The error rate of 16.13% (resp. 14.30%) for Network 8 (resp. Network 9) is due to the fact that 978 (resp. 1,399) trees were classified as Network 9 (resp. Network 8) instead of Network 8 (resp. Network 9). Thus, the classifier hesitates mainly between these two scenarios. Note also the fair performances of the classifier for Network 1 and Network 12, despite the fact that Network 1 is displayed by Network 12.

Moving on to the general inheritance prior (Table 9), the prior error remained low for Networks 1, 12, 15 and 16, and it was still approximately equal to 16% for Network 8. Surprisingly, the error rate for Network 9 increased from 14.30% to 19.14%. In other words, it seems that considering a larger parameter space for $\gamma$ makes things more difficult for recovering Network 9. With this new prior, the inheritance probability corresponding to the oldest reticulation node of Network 9 was drawn from ~~a~~ $U(0.05, 0.95)$, whereas it was previously sampled with the specific prior from ~~a~~ $U(0.05, 0.20)$.

Overall, on the basis of all investigated scenarios, the prior error rate was found smaller for SnappNet's prior (6.49%, see Table 8) than for the prior "free" from SnappNet (7.50%, see Table 9). **Incorporating a very specific prior for the inheritance probability helps to differentiate the evolutionary histories more easily. SnappNet was found rewarding since it allowed to calibrate a more performant prior.**

Impact of the prior on the instantaneous rates:

We studied the impact of the prior on the instantaneous rates (Tables 10-11) as compared to the use of the Maximum Likelihood Estimates (MLEs) (Tables 8-9). When using the specific inheritance prior (Tables 8 and 10), we observed similar misclassification rates: the prior error rate was found to be equal to 6.49% on the basis of MLEs (Table 8) and equal to 6.62% with a prior on $u$ and $v$ (Table 10). In the same way, for the general inheritance prior (Tables 9 and 11), we observed very close error rates: 7.50% with MLEs (Table 9) and 7.75% with a prior on instantaneous rates (Table 11).

**Choosing a prior on $u$ and $v$ or considering the MLEs, does not seem to have an impact on performances of the classifier (see the similar misclassification rates between Tables 10 and 8, and between Tables 11 and 9).** Note that in both cases, instantaneous rates always relied on SnappNet's estimates since (a) the prior was tuned thanks to SnappNet and (b) the MLEs were obtained by SnappNet. We can also mention that with the prior on instantaneous rates (Tables 10-11), we observed the same behavior as previously without prior (Tables 8-9): the inheritance prior had a large effect on Network 9 since the error rate increased from 13.99% (cf. Table 10, specific inheritance prior) to 19.72% (cf. Table 11, general inheritance prior).

Impact of the population size prior:

We compared a general $\Gamma(1, 0.1)$ prior on population sizes (Tables 12-15), to a prior specific to SnappNet (Tables 8-11). We can notice that Networks 8 and 9 were recovered more easily with the general prior. Indeed, on average, the prior error rate decreased from 16.13% (resp. 16.68%) to 5.55% (resp. 10.31%) for Network 8 (resp. Network 9). In contrast, it became more difficult to recover Network 12 with the general prior than with SnappNet's prior. Indeed, the following increases in terms of error rate were observed: an increase either (a) from 1.44% to 16.77% (Table 8 vs Table 12), either (b) from 4.84% vs 18.69% (Table 9 vs Table 13), either (c) from 1.51% to 16.84% (Table 10 vs Table 14), or (d) from 4.76% to 17.89% (Table 11 vs Table 15).

**Overall, the two investigated priors on population sizes gave similar performances**: the average error rate was found to be equal to 7.01% for the general prior, and to 7.09% for SnappNet's prior.

Impact of the number of trees in the forest:

We let the number of trees vary, in order to find the optimal number of trees to grow the forest. According to Figures 26- 27, we can observe the decrease of the error when the number of trees increases. **A forest of 1,000 trees seems to be a good fit for all our experiments since the errors were stabilized.** Consequently, we considered more trees than the 500 trees advised by the package .

The most important SS:

The most important SS are illustrated in Figures 28- 29. It is clear that **the Hils statistics [4] are the most important contributors to the classifier**. It is not surprising since they refer to hybridization tests. Note also that **the D-statistic by [2, 5] and the estimators $\frac{\widehat{f_3}}{\widehat{f_3}+\widehat{f_4}}$ and $\frac{\widehat{f_1}}{\widehat{f_1}+\widehat{f_2}}$ (see [1]) play an important role in our ABC-RF analysis**.

Impact of the number of rows of the reference table:

Table 17 gives, for each prior described in Table 16, the misclassification rate as a function of the number of simulations used to build the reference table. For instance, for the first prior, the misclassification rate was found to be equal to (a) 7.82% for $N_{\text{Ref}}$ =21,000, (b) to 7.03% for $N_{\text{Ref}}$ =42,000 and (c) to 6.49% for $N_{\text{Ref}}$ =84,000. We observed globally the same values whatever the prior under study. **As expected, the prior error rate decreases when the number of rows of the reference table increases** (see Table 17). Since **the out of bag error rates (i.e. prior error rates) obtained for $N_{\text{Ref}}$ =84,000 was** already **fully satisfactory**, we did not explore larger reference tables.

Conclusion of the simulation study:

**To conclude, our method enjoyed very good performances on simulated data, whatever the prior under study**. Indeed, on average, the prior error rate was found to be equal to 7.05%.

SnappNet's prior on the inheritance probability was found the most rewarding but we have to keep in mind that even when our investigated priors were "free" from SnappNet, branch lengths and instantaneous rates always relied on SnappNet's estimates. This might explain the fair accuracy in all settings. We refer to [8] for the importance of branch lengths in ABC approaches. Last, we found that our hybrid approach should be tuned with $N_{\text{Ref}}$ =84,000 and with 1,000 trees.

**Algorithm 1: ChangeBranchLength**

**Input:** network $N$
**Output:** network $N$
Sample an internal tree node $\nu$ at random from the network $N$
// $\nu$ can not be the root, and not the leaf
**if** <u>$\nu$ is reticulation node</u> **then**
  // $\nu$ has 2 parents, denoted parent1 and parent2
  // $\nu$ has one child, denoted child

  // generate a random number $u$
  // from the following Uniform distribution
  $u \sim \mathbb{U}\left[-\text{dist}(\nu, \text{child1}); \min(\text{dist}(\nu, \text{parent1}), \text{dist}(\nu, \text{parent2}))\right]$
  // update the branch length between $\nu$ and child
  dist($\nu$, child) = dist($\nu$, child) + $u$
  // update the branch length between $\nu$ and parent2
  dist($\nu$, parent2) = dist($\nu$, parent2) - $u$
  // update the branch length between $\nu$ and parent1
  dist($v$, parent1) = dist($v$, parent1) - $u$
**else**
  // $\nu$ is not a reticulation node
  // $\nu$ has one parent, denoted parent
  // $\nu$ has two children, denoted child1 and child2

  // generate a random number $u$
  // from the following Uniform distribution
  $u \sim \mathbb{U}\left[-\min(\text{dist}(\nu, \text{child1}), \text{dist}(\nu, \text{child2})); \text{dist}(\nu, \text{parent})\right]$
  // update the branch length between $\nu$ and child1
  dist($\nu$, child1) = dist($\nu$, child1) + $u$
  // update the branch length between $\nu$ and child2
  dist($\nu$, child2) = dist($\nu$, child2) + $u$
  // update the branch length between $\nu$ and parent
  dist($\nu$, parent) = dist($\nu$, parent) - $u$
**return** $N$

**Table 8. Confusion matrix obtained by ABC-RF, when the inheritance probability $\gamma$ was drawn from the uniform distribution $U(\max(0.05, \widetilde{\gamma} - 0.1), \min(0.95, \widetilde{\gamma} + 0.1))$, where $\widetilde{\gamma}$ denotes the MLE of $\gamma$ obtained with SnappNet. No prior was considered for the instantaneous rates $u$ and $v$ (MLE values). The error rate, given in last column, denotes the misclassification rate for each scenario. The size $N_{\text{Ref}}$ of the reference table was 84,000 before filtering for missing data and 72,406 after filtering. The out of bag error rate was found to be equal to $6.49\%$.**

| | Net 1 | Net 8 | Net 9 | Net 12 | Net 15 | Net 16 | Error Rate |
|---|---|---|---|---|---|---|---|
| **Net 1** | 10,565 | 290 | 57 | 3 | 94 | 28 | 4.28% |
| **Net 8** | 354 | 10,055 | 978 | 567 | 5 | 30 | 16.13% |
| **Net 9** | 75 | 1,399 | 10,330 | 197 | 25 | 27 | 14.30% |
| **Net 12** | 30 | 87 | 58 | 12,357 | 0 | 5 | 1.44% |
| **Net 15** | 45 | 6 | 9 | 1 | 11,709 | 93 | 1.30% |
| **Net 16** | 99 | 62 | 28 | 1 | 49 | 12,688 | 1.85% |

**Table 9. Same as Table 8 except that the inheritance probability $\gamma$ was drawn from the uniform distribution $U(0.05, 0.95)$. The size $N_{\text{Ref}}$ of the reference table was 84,000 before filtering for missing data and 72,627 after filtering. The out of bag error rate was found to be equal to $7.50\%$.**

| Network ID | Net 1 | Net 8 | Net 9 | Net 12 | Net 15 | Net 16 | Error Rate |
|---|---|---|---|---|---|---|---|
| **Net 1** | 10,727 | 292 | 25 | 10 | 9 | 3 | 3.06% |
| **Net 8** | 370 | 10,086 | 964 | 601 | 2 | 0 | 16.11% |
| **Net 9** | 67 | 1,533 | 9,765 | 710 | 4 | 0 | 19.14% |
| **Net 12** | 100 | 383 | 112 | 11,946 | 0 | 0 | 4.84% |
| **Net 15** | 69 | 4 | 1 | 0 | 11858 | 19 | 0.77% |
| **Net 16** | 35 | 9 | 7 | 3 | 113 | 12,800 | 1.29% |

**Table 10. Same as Table 8 except that the rate $u$ was drawn from uniform distribution $U(\max(0, \widetilde{u} - 0.0005), \widetilde{u} + 0.0005)$, where $\widetilde{u}$ refers to the MLE of $u$ obtained with SnappNet. The size $N_{\text{Ref}}$ of the reference table was 84,000 before filtering for missing data and 72,458 after filtering. The out of bag error rate was found to be equal to $6.62\%$.**

| Network ID | Net 1 | Net 8 | Net 9 | Net 12 | Net 15 | Net 16 | Error Rate |
|---|---|---|---|---|---|---|---|
| **Net 1** | 10,526 | 309 | 58 | 4 | 129 | 30 | 4.79% |
| **Net 8** | 375 | 9,917 | 1,090 | 557 | 9 | 22 | 17.15% |
| **Net 9** | 79 | 1,376 | 10,366 | 197 | 17 | 17 | 13.99% |
| **Net 12** | 39 | 103 | 41 | 12,319 | 0 | 2 | 1.48% |
| **Net 15** | 39 | 6 | 3 | 0 | 11,748 | 80 | 1.07% |
| **Net 16** | 89 | 57 | 29 | 1 | 41 | 12,783 | 1.67% |

**Table 11. Same as Table 9 except that the rate $u$ was drawn from uniform distribution $U(\max(0, \widetilde{u} - 0.0005), \widetilde{u} + 0.0005)$, where $\widetilde{u}$ refers to the rate estimated by maximum likelihood with SNAPPNET. The size $N_{\text{Ref}}$ of the reference table was 84,000 before filtering for missing data and 72,531 after filtering. The out of bag error rate was found to be equal to $7.75\%$.**

| Network ID | Net 1 | Net 8 | Net 9 | Net 12 | Net 15 | Net 16 | Error Rate |
|---|---|---|---|---|---|---|---|
| **Net 1** | 10,696 | 304 | 24 | 21 | 29 | 2 | 3.26% |
| **Net 8** | 392 | 10,039 | 949 | 676 | 0 | 0 | 16.73% |
| **Net 9** | 72 | 1,592 | 9,678 | 711 | 2 | 0 | 19.72% |
| **Net 12** | 102 | 408 | 114 | 11,887 | 0 | 0 | 4.99% |
| **Net 15** | 60 | 3 | 1 | 0 | 11,770 | 23 | 0.73% |
| **Net 16** | 25 | 13 | 13 | 1 | 103 | 12,841 | 1.19% |

**Table 12. Same as Table 8 except that the prior on $\theta$ is $\Gamma(1, 0.1)$. The size $N_{\text{Ref}}$ of the reference table was 84,000 before filtering for missing data and 75,974 after filtering. The out of bag error rate was found to be equal to $6.26\%$.**

| Network ID | Net 1 | Net 8 | Net 9 | Net 12 | Net 15 | Net 16 | Error Rate |
|---|---|---|---|---|---|---|---|
| **Net 1** | 11,754 | 230 | 26 | 573 | 75 | 47 | 7.49% |
| **Net 8** | 67 | 12,108 | 216 | 187 | 16 | 0 | 3.86% |
| **Net 9** | 80 | 523 | 11,784 | 295 | 4 | 9 | 7.18% |
| **Net 12** | 1,332 | 613 | 167 | 10,610 | 5 | 22 | 16.77% |
| **Net 15** | 21 | 7 | 3 | 3 | 12,443 | 15 | 0.39% |
| **Net 16** | 120 | 26 | 11 | 34 | 25 | 11,523 | 1.69% |

**Table 13. Same as Table 9 except that the prior on $\theta$ is $\Gamma(1, 0.1)$. The size $N_{\text{Ref}}$ of the reference table was 84,000 before filtering for missing data and 75,512 after filtering. The out of bag error rate was found to be equal to $7.83\%$.**

| Network ID | Net 1 | Net 8 | Net 9 | Net 12 | Net 15 | Net 16 | Error Rate |
|---|---|---|---|---|---|---|---|
| **Net 1** | 11,724 | 235 | 48 | 477 | 16 | 8 | 6.27% |
| **Net 8** | 71 | 11,644 | 600 | 226 | 3 | 0 | 7.17% |
| **Net 9** | 127 | 1,009 | 10,898 | 568 | 0 | 2 | 13.53% |
| **Net 12** | 1,434 | 593 | 340 | 10,305 | 1 | 1 | 18.69% |
| **Net 15** | 33 | 7 | 1 | 5 | 12,331 | 11 | 0.46% |
| **Net 16** | 37 | 1 | 17 | 4 | 40 | 12,695 | 0.77% |

**Table 14. Same as Table 10 except that the prior on $\theta$ is $\Gamma(1, 0.1)$. The size $N_{\text{Ref}}$ of the reference table was 84,000 before filtering for missing data and 75,958 after filtering. The out of bag error rate was found to be equal to $6.35\%$.**

| Network ID | Net 1 | Net 8 | Net 9 | Net 12 | Net 15 | Net 16 | Error Rate |
|---|---|---|---|---|---|---|---|
| **Net 1** | 11,665 | 232 | 16 | 592 | 91 | 64 | 7.86% |
| **Net 8** | 76 | 12,073 | 202 | 209 | 15 | 0 | 3.99% |
| **Net 9** | 75 | 529 | 11,784 | 291 | 3 | 13 | 7.17% |
| **Net 12** | 1,324 | 625 | 171 | 10,602 | 6 | 21 | 16.84% |
| **Net 15** | 29 | 8 | 2 | 9 | 12,438 | 9 | 0.45% |
| **Net 16** | 136 | 17 | 5 | 26 | 30 | 12,570 | 1.67% |

**Table 15. Same as Table 11 except that $\theta$ was drawn from a $\Gamma(1, 0.1)$. The size $N_{\text{Ref}}$ of the reference table was 84,000 before filtering for missing data and 75,943 after filtering. The out of bag error rate was found to be equal to $7.61\%$.**

| Network ID | Net 1 | Net 8 | Net 9 | Net 12 | Net 15 | Net 16 | Error Rate |
|---|---|---|---|---|---|---|---|
| **Net 1** | 11,858 | 209 | 49 | 458 | 14 | 10 | 5.87% |
| **Net 8** | 66 | 11,688 | 597 | 242 | 2 | 0 | 7.20% |
| **Net 9** | 107 | 1,041 | 10,937 | 539 | 3 | 0 | 13.38% |
| **Net 12** | 1,367 | 563 | 356 | 10,503 | 1 | 1 | 17.89% |
| **Net 15** | 20 | 7 | 1 | 8 | 12,513 | 8 | 0.35% |
| **Net 16** | 51 | 0 | 20 | 2 | 33 | 12,669 | 0.83% |

**Table 16. Summary of all the combinations of priors investigated in this study. Recall that the parameters $\gamma$, $u$ and $\theta$, denote respectively, the inheritance probability, the instantaneous mutation rate and the population size. $\widetilde{\gamma}$, $\widetilde{\theta}$ and $\widetilde{u}$ denote the MLEs of $\gamma$, $u$ and $\theta$, obtained with SnappNet. $U(.)$ and $\Gamma(.)$ refer respectively to the Uniform and Gamma distributions.**

| Prior in \ Parameter | $\gamma$ | $u$ | $\theta$ |
|---|---|---|---|
| **Table 8** | $U(\max(0.05, \widetilde{\gamma} - 0.1), \min(0.95, \widetilde{\gamma} + 0.1))$ | No prior ($\widetilde{u}$) | $\Gamma(1, \widetilde{\theta})$ |
| **Table 9** | $U(0.05, 0.95)$ | No prior ($\widetilde{u}$) | $\Gamma(1, \widetilde{\theta})$ |
| **Table 10** | $U(\max(0.05, \widetilde{\gamma} - 0.1), \min(0.95, \widetilde{\gamma} + 0.1))$ | $U(\max(0, \widetilde{u} - 0.0005), \widetilde{u} + 0.0005)$ | $\Gamma(1, \widetilde{\theta})$ |
| **Table 11** | $U(0.05, 0.95)$ | $U(\max(0, \widetilde{u} - 0.0005), \widetilde{u} + 0.0005)$ | $\Gamma(1, \widetilde{\theta})$ |
| **Table 12** | $U(\max(0.05, \widetilde{\gamma} - 0.1), \min(0.95, \widetilde{\gamma} + 0.1))$ | No prior ($\widetilde{u}$) | $\Gamma(1, 0.1)$ |
| **Table 13** | $U(0.05, 0.95)$ | No prior ($\widetilde{u}$) | $\Gamma(1, 0.1)$ |
| **Table 14** | $U(\max(0.05, \widetilde{\gamma} - 0.1), \min(0.95, \widetilde{\gamma} + 0.1))$ | $U(\max(0, \widetilde{u} - 0.0005), \widetilde{u} + 0.0005)$ | $\Gamma(1, 0.1)$ |
| **Table 15** | $U(0.05, 0.95)$ | $U(\max(0, \widetilde{u} - 0.0005), \widetilde{u} + 0.0005)$ | $\Gamma(1, 0.1)$ |

**Table 17. Misclassification rate as a function of the size $N_{\text{Ref}}$ of the reference table (before filtering for missing data), and as a function of the prior distribution.**

| Prior \ $N_{\text{Ref}}$ | 21,000 | 42,000 | 84,000 |
|---|---|---|---|
| **Table 8** | 7.81% | 7.03% | 6.49% |
| **Table 9** | 8.55% | 7.81% | 7.50% |
| **Table 10** | 7.69% | 7.18% | 6.62% |
| **Table 11** | 8.66% | 8.39% | 7.74% |
| **Table 12** | 7.30% | 6.79% | 6.26% |
| **Table 13** | 8.60% | 8.40% | 7.83% |
| **Table 14** | 7.26% | 6.84% | 6.35% |
| **Table 15** | 8.80% | 8.17% | 7.61% |

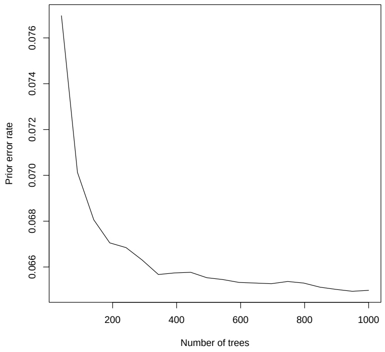


**No prior for $u$ and $v$**
$\gamma \sim U(\max(0.05, \widetilde{\gamma} - 0.1), \min(0.95, \widetilde{\gamma} + 0.1))$

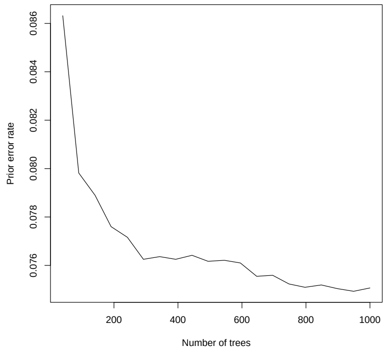


**No prior for $u$ and $v$**
$\gamma \sim U(0.05, 0.95)$

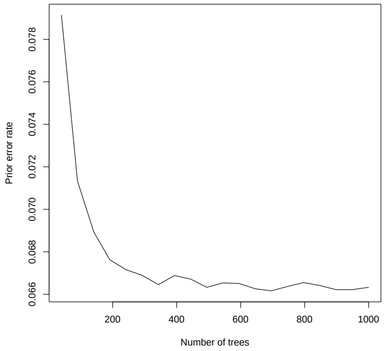


$u \sim U(\max(0, \widetilde{u} - 0.0005), \widetilde{u} + 0.0005)$
$\gamma \sim U(\max(0.05, \widetilde{\gamma} - 0.1), \min(0.95, \widetilde{\gamma} + 0.1))$

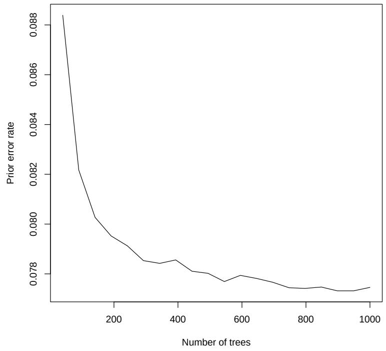


$u \sim U(\max(0, \widetilde{u} - 0.0005), \widetilde{u} + 0.0005)$
$\gamma \sim U(0.05, 0.95)$

**Figure 26.** Prior error rate as a function of the number of trees in the forest, and as a function of the considered priors. In all cases, $\theta$ was drawn from a $\Gamma(1, \widetilde{\theta})$. The focus is only on priors associated to Tables 8-11.

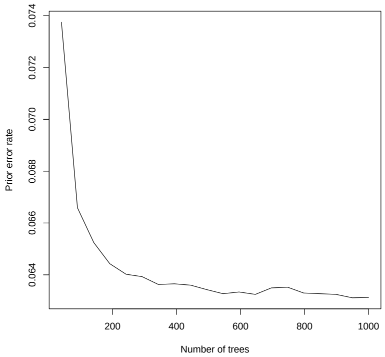


**No prior for $u$ and $v$**
$\gamma \sim U(\max(0.05, \widetilde{\gamma} - 0.1), \min(0.95, \widetilde{\gamma} + 0.1))$

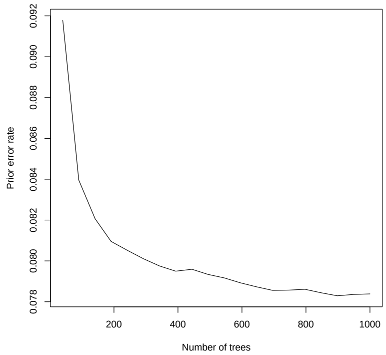


**No prior for $u$ and $v$**
$\gamma \sim U(0.05, 0.95)$

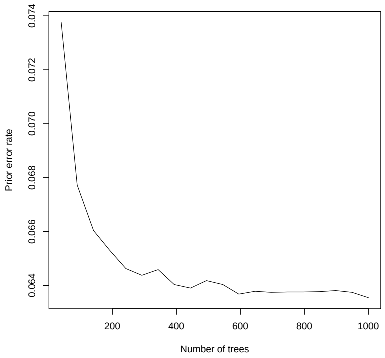


$u \sim U(\max(0, \widetilde{u} - 0.0005), \widetilde{u} + 0.0005)$
$\gamma \sim U(\max(0.05, \widetilde{\gamma} - 0.1), \min(0.95, \widetilde{\gamma} + 0.1))$

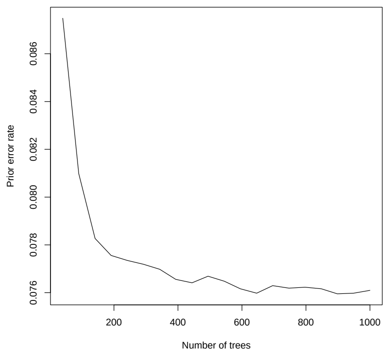


$u \sim U(\max(0, \widetilde{u} - 0.0005), \widetilde{u} + 0.0005)$
$\gamma \sim U(0.05, 0.95)$

**Figure 27.** Prior error rate as a function of the number of trees in the forest, and as a function of the considered priors. In all cases, $\theta$ was drawn from a $\Gamma(1, 0.1)$. The focus is only on priors associated to Tables 12-15.

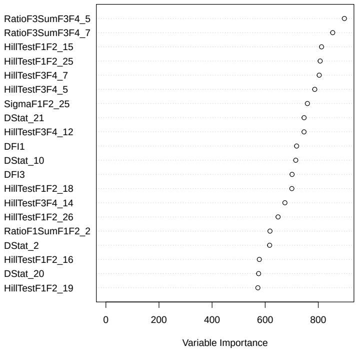


**No prior for $u$ and $v$**
$\gamma \sim U(\max(0.05, \widetilde{\gamma} - 0.1), \min(0.95, \widetilde{\gamma} + 0.1))$

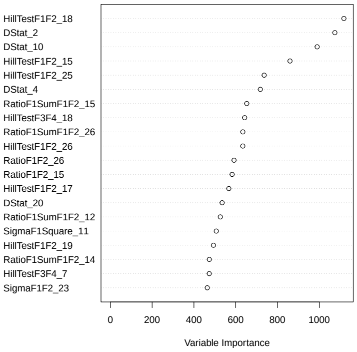


**No prior for $u$ and $v$**
$\gamma \sim U(0.05, 0.95)$

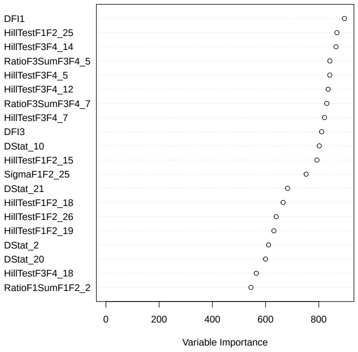


$u \sim U(\max(0, \widetilde{u} - 0.0005), \widetilde{u} + 0.0005)$
$\gamma \sim U(\max(0.05, \widetilde{\gamma} - 0.1), \min(0.95, \widetilde{\gamma} + 0.1))$

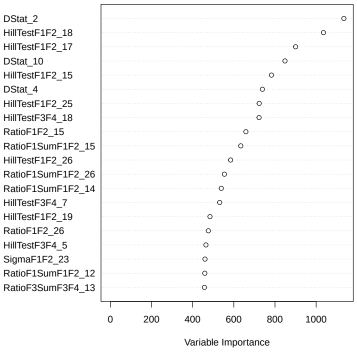


$u \sim U(\max(0, \widetilde{u} - 0.0005), \widetilde{u} + 0.0005)$
$\gamma \sim U(0.05, 0.95)$

**Figure 28.** Contributions of the twenty most important SS as a function of the considered priors. In all cases, $\theta$ was drawn from a $\Gamma(1, \widetilde{\theta})$. The focus is only on priors associated to Tables 8-11.

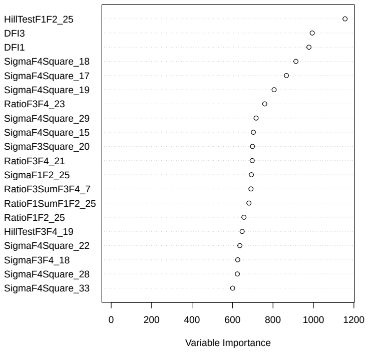


**No prior for** $u$ **and** $v$
$\gamma \sim U(\max(0.05, \widetilde{\gamma} - 0.1), \min(0.95, \widetilde{\gamma} + 0.1))$

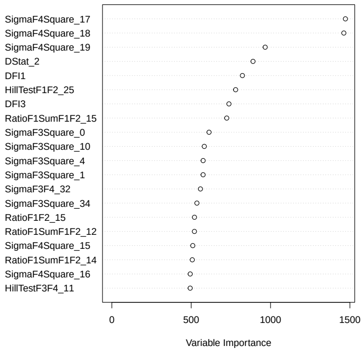


**No prior for** $u$ **and** $v$
$\gamma \sim U(0.05, 0.95)$

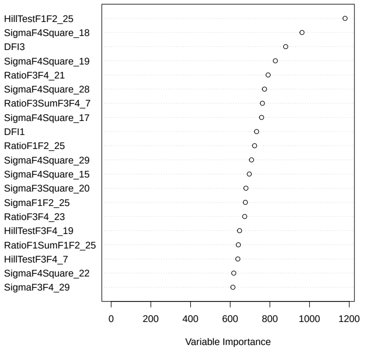


$u \sim U(\max(0, \widetilde{u} - 0.0005), \widetilde{u} + 0.0005)$
$\gamma \sim U(\max(0.05, \widetilde{\gamma} - 0.1), \min(0.95, \widetilde{\gamma} + 0.1))$

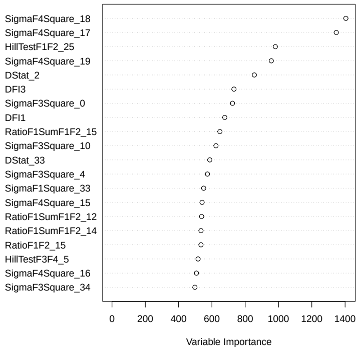


$u \sim U(\max(0, \widetilde{u} - 0.0005), \widetilde{u} + 0.0005)$
$\gamma \sim U(0.05, 0.95)$

**Figure 29.** Contributions of the twenty most important summary statistics as a function of the considered priors. In all cases, $\theta$ was drawn from a $\Gamma(1, 0.1)$. The focus is only on priors associated to Tables Tables 12-15.
